# Frontal Vulnerability vs. Temporal Resilience within the core Language Network: Neuro-compensatory mechanisms underline differential language aging trajectories

**DOI:** 10.1101/2025.04.29.651293

**Authors:** Partika Jain, Azman Akhter, Arpan Banerjee

**Author notes:** Corresponding Author: Arpan Banerjee.

## Abstract

Age-related decline underlies cognitive functions such as sensorimotor control, executive functioning, memory, and language production (LP), whereas language comprehension (LC) tends to remain intact or improve across healthy adult lifespan. This preservation likely stems from structural and functional integrity within core language network (cLAN) regions. To investigate this hypothesis, we analyzed the relationships among brain’s resting-state functional connectivity (FC), structural connectivity (SC), and language behaviour (LC and LP) using a cross-sectional cohort of healthy adults (N = 652; M/F = 322/330; aged 18–88) from the Cambridge Centre for Ageing and Neuroscience (Cam-CAN) dataset. Six cognitive tasks assessing LC and LP were employed, with neuroimaging measures focused on region-specific connections within the cLAN. Using generalized additive mixed models (GAMMs), complex brain-behaviour interactions were identified. Behavioral analyses revealed established age-related dichotomy, LC abilities in vocabulary and proverb comprehension improved and in syntactic and semantic comprehension remained stable, whereas LP tasks, e.g., verbal fluency, picture priming, and tip of tongue exhibited significant decline across the lifespan. SC exhibited decline in both intra-and inter-hemispheric fronto-temporal and frontal lobe connections, contrasted by preserved or enhanced temporal lobe connectivity, supporting a pattern of frontal vulnerability concomitant with temporal resilience. Age-related FC patterns demonstrated overall preservation, reflecting compensatory mechanisms to sustain functional integrity despite structural degradation. GAMM analyses revealed complex relationships between brain connectivity and language performance across age. Thus, integrating knowledge of brain structure, function, and language abilities, we identified the brain network mechanisms associated with dichotomous language behavior along lifespan.

**Significance:** Understanding age-related changes in language abilities is vital for early detection of cognitive decline and intervention development. This study reveals heterogeneous language aging trajectories: language production often declines, while comprehension remains stable or improves. Notably, comprehension tasks with high executive and working memory demands also show resilience, suggesting compensatory mechanisms beyond crystallized knowledge. Using multimodal neuroimaging in a large adult lifespan cohort, we link behavioral patterns to distinct network dynamics within the core language system. We uncover relative vulnerability in frontal brain regions, with pronounced structural decline in fronto-temporal pathways, contrasted by resilience in temporal connectivity, which is preserved or strengthened. These findings highlight regionally specific vulnerabilities and compensatory functional adaptations, refining current models and providing a framework to guide research promoting healthy cognitive aging.

## 1. Introduction

While age-related decline is ubiquitous in most cognitive functions, language behavior exhibits an intriguing idiosyncrasy which is unanswered in existing literature. Language production (LP) often deteriorates with age (Baciu et al., 2021; Burke & Shafto, 2004; Diaz et al., 2016; H. Zhang & Diaz, 2023a), e.g, older adults experience difficulties of stuttering, increased retrieval failures, more disfluent speech, and tip-of-the-tongue experiences (Burke & Shafto, 2004; Shafto et al., 2007). In contrast, language comprehension (LC) tend to remain stable or even improve (Ferré et al., 2019; Salthouse, 2014; Shafto & Tyler, 2014; Sommers et al., 2011; Verhaegen & Poncelet, 2013). Yet, under specified conditions of complexity elderly exhibit reduced comprehension, underscoring that resilience is relative rather than absolute (Goral et al., 2011; Kemper et al., 2004; Obler et al., 1991). Additionally, language abilities may be the earliest cognitive functions that are impaired during neurodegenerative conditions (Mascali et al., 2018; Rafiq et al., 2022; van Boxtel & Lawyer, 2021). Thus, mapping lifespan-scale trajectory of language abilities on brain metrics may accelerate identification of biomarkers for disorders such as Alzheimer’s and dementia.

Brain mapping using resting-state functional connectivity (rsFC) exhibits 80% shared variance with activity during cognitive tasks in similar areas (Cole et al., 2014), age-related decline across whole brain (Chan et al., 2017; Jiang et al., 2022; Khalilian et al., 2024) and also within the extended language network (Bolognini & Martin, 2023; Ferré et al., 2019; Hoffman & Morcom, 2018; H. Zhang et al., 2021; H. Zhang & Diaz, 2023b). Emerging evidences suggest existence of Core language network (cLAN), comprising frontotemporal regions (Fedorenko et al., 2024) that demonstrate remarkable preservation of structural and functional integrity across the adult lifespan (Billot et al., 2024; Campbell et al., 2016; Pistono et al., 2021; Rafiq et al., 2022; Shafto & Tyler, 2014). Few have examined rsFC in cLAN in younger and older adults and their relationship with behavioral performance (Ferré et al., 2019; Gertel et al., 2020; H. Zhang et al., 2021). Interestingly, Ferré et al. (2019) observed no interaction between the rsFC and language behavior scores, potentially due to their reliance on linear methods (Bolognini & Martin, 2023).

Fewer studies have examined age-related changes in structural connectivity (SC; Damoiseaux, 2017; Davis et al., 2009a; Fletcher et al., 2018; Henriques et al., 2023; Pistono et al., 2021; Westlye et al., 2010), particularly within the cLAN. Age-related decline has been reported across several measures, including white matter volume, gray matter density, cortical thickness, fractional anisotropy (FA), and other microstructural properties of white matter tracts (Davis et al., 2009b; Fletcher et al., 2018; Henriques et al., 2023; Pistono et al., 2021; Westlye et al., 2010). Fronto-temporal connections of the dorsal pathway showed more pronounced decline in FA values across age (Yang et al., 2014). Houston et al., 2019 showed that white matter integrity, particularly in the frontal-occipital fasciculus (IFOF) and superior longitudinal fasciculus (SLF) may be critical in language performance. However, the study involved a cohort with a mean age of 40, at which global connectivity strength and efficiency among myelin-weighted connections are known to peak (Bosticardo et al., 2024), and hence may not adequately capture the progressive decline in structural connectivity prevalent in elderly.

Ongoing debate rages in literature whether the neurofunctional changes represent compensatory mechanisms (Cabeza et al., 2018; Davis et al., 2008; Naik et al., 2017) or reflect decline processes such as de-differentiation (Diaz et al., 2014; Li et al., 2001; H. Zhang & Diaz, 2023a). How do these mechanisms support age-related language behavior trajectories within the cLAN? Current study addressed this question by integrating epidemiological, cognitive, neurofunctional, and structural data from a large, age-homogeneous adult cohort, and focused on detailed nonlinear regression analyses between cLAN regions (Billot et al., 2024; Branco et al., 2020; Bulut, 2023; Fedorenko & Blank, 2020; Shafto & Tyler, 2014) and language behavior across adult lifespan.

## 2. Materials and Methods

### 2.1 Participants

The neuroimaging data of cognitively healthy participants (N = 652; M/F = 322/330; age range: 18-88) were obtained from a cross-sectional population-based adult cohort at the Cambridge Centre for Ageing and Neuroscience repository (CamCAN, Stage 2). All participants underwent a series of neuropsychological tests with strict exclusion criteria (Shafto et al., 2014; Taylor et al., 2017), including cognitive impairment (Mini-Mental State Examination [MMSE] score <25, severe memory deficit), communication difficulties (inability to hear 35 dB at 1000 Hz in either ear, poor vision < 20/100, and poor fluency in English), self-reported substance abuse, and experiencing any serious health problems. The education score of each participant was derived by assigning a difficulty level to each education stage provided by the UK government’s official website (https://www.gov.uk/what-different-qualification-levels-mean/list-of-qualification-levels) and then aggregating these values over the participant’s lifespan. All participants provided written informed consent, and this study was approved by the Research Ethics Committee of Cambridgeshire 2 (now East of England-Cambridge Central). Subsequently, only participants with complete neuroimaging data comprising T1, resting state fMRI (rs-fMRI), and structural T1 and diffusion weighted MRI (DWI) data were considered in the current study (N = 642). Additionally, 33 participants were excluded because of poor data quality and previously reported artifacts (for details see: https://camcan-archive.mrc-cbu.cam.ac.uk/dataaccess/issues.php) Considering that left- and right-handed participants may have hemispherically opposite functional dominance (Knecht et al., 2000) and organization (Tejavibulya et al., 2022), only 558 (Male = 276, age range: 18-88) right-handed participants were included in the subsequent analysis.

### 2.2 Language behaviour tasks

Six cognitive tests, conducted externally from the scanner, were employed to assess two primary language domains: language comprehension and production (LC and LP). These assessments comprised three tasks each for LC and LP. Where applicable, each language task was further categorized into multiple subdomains (accuracy and response time measures) based on linguistic complexity. Specifically, the LC domain included the following subdomains: Vocabulary (Vocab), Proverb comprehension (Proverb), and Sentence comprehension task (Synsem), which was further subdivided into different subdomains: Syntactically complex sentences (Syn_comp), Semantically complex sentences (Sem_comp), and their associated response time measures (RT_Syn_comp and RT_Sem_comp). Accuracy in the Syn_comp task was quantified by deriving a composite measure that integrates accuracy rates across different sentence continuation types. This approach allowed for the evaluation of performance under conditions of both expected and unexpected linguistic structures. Similar composite measures were constructed to assess performance in the other Synsem subdomains, providing a comprehensive evaluation of syntactic and semantic comprehension abilities.

The LP domain comprised Verbal Fluency (VF), Tip-of-the-Tongue (ToT), and the Picture Priming task, which was further subdivided into different subdomains High Phonological similarity (High_phon), Low Phonological similarity (Low_phon), High Semantic similarity (High_sem), Low Semantic similarity (Low_sem) based on semantic and phonological similarities between prime and the target, and their corresponding response time measures (RT_high_phon, RT_low_phon, RT_high_sem, RT_low_sem). For each of these subdomains, performance was quantified by comparing responses under each condition with a neutral or unrelated baseline condition. For example, accuracy in the High_sem condition was calculated by assessing the improvement in accuracy when the prime was highly semantically related to the target, relative to when the prime and target were unrelated. This approach allowed us to isolate the specific effects of semantic and phonological similarity on production performance. To facilitate consistent interpretation across all measures, the ToT scores and response time variables were sign-reversed so that higher values uniformly indicated better performance. A detailed structure of the tasks and their corresponding subdomains is provided in Table 1, with comprehensive details available in (Shafto et al., 2014).

### 2.3 MRI data acquisition

The structural (e.g., T1, Diffusion weighted imaging (DWI)) and functional data (Resting-state functional magnetic resonance imaging (rs-fMRI)) were collected at a single site (MRC Cognition and Brain Sciences Unit) using 3 Tesla by CamCAN consortium in a Siemens scanner with a 32-channel head coil, where head movement was restricted with memory foam cushions. A high-resolution 3D structural T1-weighted image was acquired using the Magnetization-Prepared Rapid Gradient-Echo (MPRAGE) sequence with a Generalized Autocalibrating Partially Parallel Acquisition (GRAPPA) acceleration factor, repetition time (TR) = 2250 ms, echo time (TE) = 2.99 ms, inversion time (TI) = 900 ms, flip angle α = 9°, field of view (FOV) = 256 × 240 × 192 mm, isotropic resolution = 1 mm, acquisition time of 4 min and 32 s. rs-fMRI scans were undertaken with echo planar imaging (EPI) sequences comprising of 261 volumes with eye-closed condition. Each volume consisted of 32 axial slices with slice thickness 3.7 mm, TR = 1,970 ms, TE = 30 ms, flip angle α = 78°, FOV = 192 × 192 mm^2^, resolution = 3 × 3 × 4.44 mm acquired in descending order for 8 mins and 40 s.

Diffusion-weighted MR images (DWI) were obtained using a twice-refocused spin-echo (TRSE) echo-planar imaging sequence to reduce distortions caused by Eddy currents. The imaging scheme comprised 30 diffusion gradient directions for each of the two nonzero b-values (1,000 and 2,000 s/mm²) and three images with zero b-values (0 s/mm²). The other imaging parameters were as follows: 66 axial slices, isotropic resolution = 2 mm, TR/TE = 9100/104 ms, acquisition time = 10 min, and 2 s.

### 2.4 rs-fMRI preprocessing

Resting-state functional imaging data of each participant were preprocessed and denoised using the CONN toolbox version 2022a (https://www.nitrc.org/projects/conn ; Whitfield-Gabrieli & Nieto-Castanon, 2012) in conjunction with Statistical Parametric Mapping (SPM 12) and Matlab 2022a. The first five scans from each run were disregarded to allow the magnetization to reach a steady state (Caballero-Gaudes & Reynolds, 2017). Functional realignment was performed by the co-registration and resampling of all functional scans to a reference image using b-spline interpolation (Andersson et al., 2001). Temporal misalignment between different slices of functional data was corrected using a slice-timing correction (STC) procedure (Henson et al., 1999). Outlier detection was performed with the help of Artefact Detection Tool (ART, https://www.nitrc.org/projects/artifact_detect/), in which framewise displacement above 0.9 mm or global blood-oxygen-level-dependent (BOLD) signal changes above 5 standard deviations were flagged as potential outliers and a new reference image was generated by averaging across all scans excluding the outliers. Subsequently, the functional and structural data were independently normalized into a standard Montreal Neurological Institute (MNI) template and segmented into gray matter, white matter, and Cerebrospinal fluid (CSF) using a unified segmentation and normalization method (Ashburner & Friston, 2005). The output data were resampled to a default 180 × 216 × 180 mm bounding box using fourth-order spline interpolation, with 2 mm and 1 mm isotropic voxels for functional and anatomical data, respectively.

### 2.5 Denoising

Physiological fluctuations have been proven to be a considerable contributor to the noise present in blood-oxygen-level-dependent (BOLD) signals (Krüger & Glover, 2001; Liu et al., 2006). To mitigate this issue, an additional preprocessing step was implemented, which involves an anatomical component-based noise correction method (aCompCor ; Behzadi et al., 2007). This algorithm utilizes principal component analysis (PCA) to characterize signals of non-neural origin, such as white matter and cerebrospinal fluid (5 components from each), and then introduces them as covariates in a general linear model (GLM) as an estimate of the physiological noise signal space. Concurrently, 12 motion parameters (three translations, three rotations, and their first-order derivatives) and a variable number of noise components from each identified outlier scan were also regressed from the BOLD signal of each voxel. A simultaneous band-pass filtering (.009 Hz – .08 Hz) was applied, using a discrete cosine transform operation, to prevent any frequency mismatch in the nuisance regression strategy (Hallquist et al., 2013). The use of the aCompCor method significantly improves the sensitivity and specificity of functional connectivity estimates; thus, global signal regression was not applied (Chai et al., 2012; Muschelli et al., 2014).

### 2.6 Core Language Network (cLAN) selection

To evaluate the variation in FC within a defined cLAN across age, a functional connectivity analysis was carried out using a FreeSurfer-based atlas, Desikan Killiany (Desikan et al., 2006). This automatic parcellation, based on hemispheric-specific anatomical features, has been shown to show high accuracy compared to manual labelling (Desikan et al., 2006), and has also been used in various language-related studies (Nilakantan et al., 2017; Papanicolaou et al., 2017; Qi et al., 2019). Recent reviews have identified language regions in frontal and temporal regions serves as the Core Language network (referred to as cLAN throughout this study; Fedorenko et al., 2024) which is supported by functional localizer based probabilistic atlases on considerable number of participants (Lipkin et al., 2022). Converging evidence indicates that the core frontotemporal language network remains largely stable throughout the adult lifespan (Shafto & Tyler, 2014; Billot et al., 2024). Landmark reviews and recent large-scale imaging studies consistently show that, unlike domain-general and executive networks which exhibit marked age-related decline and dedifferentiation, the core language network preserves its selectivity, lateralization, and internal coherence (Rafiq et al., 2022; Zhang et al., 2023; Billot et al., 2024). Additionally, functional connectivity within this network is maintained or even modestly enhanced despite structural gray matter loss, supporting stable language function in aging (Pistono et al., 2021). Thus, focusing on the well-validated core language network provides an essential baseline for understanding language aging.

Accordingly, we selected canonical cLAN parcels aligned with prior research (Billot et al., 2024; Bolognini & Martin, 2023; Branco et al., 2020; Bulut, 2023; Fedorenko et al., 2024; Fedorenko & Blank, 2020; Rafiq et al., 2022; H. Zhang et al., 2021). The language-related region of interests (ROIs) included temporal regions, namely the Middle Temporal Gyrus (MTG), Superior Temporal Gyrus (STG), and superior temporal sulcus (STS), and inferior frontal regions, namely Pars Opercularis (ParsO) and Pars Triangualris (ParsT). These ROIs were delineated bilaterally, yielding a total of 10 LAN-ROIs: 5 in left-hemisphere (LMTG, LSTG, LSTS, LParsO, LParsT) and 5 in right-hemisphere ROIs (RMTG, RSTG, RSTS, RParsO, RParsT). Note that these regions of the frontal lobe are also known in the Brodmann (BA)-based terminology as BA 44 and BA 45, making up the classical Broca’s area.

### 2.7 Functional connectome

For evaluating the functional connectivity (FC) within core language network (cLAN), mean regional blood-oxygen-level-dependent (BOLD) time series were estimated for each individual region of interest (ROI) within the cLAN. FC between two distinct brain ROIs was determined by calculating the partial correlation coefficients between the respective mean regional BOLD time series, accounting for the signals in all other brain regions. This approach provides a more precise depiction of direct functional interactions (Marrelec et al., 2006) by eradicating the influence of common sources and results in sparser FC matrices, compared to the conventional Pearson correlation approach (Smith et al., 2011; Zalesky et al., 2012). To draw statistical comparisons, the resulting connectivity values were then normalized using Fisher’s r-to-z transformation. The resultant FC matrix of 10 LAN-ROIs comprised 10 left-hemispheric FC edges, 10 right-hemispheric FC edges and 25 inter-hemispheric FC edges, yielding a total of 45 LAN FC edges (as represented in Figure 4).

### 2.8 DWI preprocessing

We developed a custom shell script (https://github.com/PartikaJain/CBDL_DWI_script) to preprocess the diffusion weighted imaging (DWI) data, which included the use of different software packages: MRtrix3 (https://www.mrtrix.org/ ; Tournier et al., 2019), FMRIB Software Library (FSL) (https://fsl.fmrib.ox.ac.uk/fsl/fslwiki ; Smith et al., 2004; Woolrich et al., 2009), Freesurfer (https://surfer.nmr.mgh.harvard.edu/ ; Fischl, 2012), and ANTs (https://github.com/stnava ; Tustison et al., 2010). Primarily, DWI data were de-noised with the function *dwidenoise*, which relies on the theory of Marchenko-Pastur Principal Component Analysis (MP-PCA ; Veraart et al., 2016), and then Gibb’s ringing artifacts were removed using the sub-voxel shift method (Kellner et al., 2016). Motion and distortion correction was applied using *dwifslpreproc* function which implements FSL integrated tools: *topup* tool was used for susceptibility-induced distortion correction (Andersson et al., 2003; S. M. Smith et al., 2004) and *eddy* for within-shell and between-shell motion and eddy current-induced distortion correction (Andersson et al., 2016). Bias field correction (Tustison et al., 2010) was applied using the MRtrix function *dwibiascorrect* with *theants* option to improve the brain mask estimation.

### 2.9 Fibre orientation distribution (FOD)

Initially, response functions for distinct tissue types (white matter, gray matter, and cerebrospinal fluid) were estimated using the *dwi2response* function with the Dhollander algorithm, which is specifically designed for multi-shell data (Dhollander et al., 2019). Subsequently, the orientation of fibres in each voxel was estimated using *dwi2fod* function with the multi-shell multi-tissue constrained spherical deconvolution (MSMT-CSD) algorithm (Jeurissen et al., 2014), which have demonstrated superior performance compared to diffusion tensor imaging (DTI) in regions with crossing-fibres (Farquharson et al., 2013; Tournier et al., 2008). Finally, to ensure the comparability of FODs across participants, global intensity normalization was conducted using the *mtnormalise* function.

### 2.10 Tractography

Anatomically Constrained Tractography (ACT) was applied to reduce false positives and improve the biological plausibility of streamline generation (Smith et al., 2012). To make the data suitable for ACT, each T1-weighted image was segmented into five tissue types (cortical gray matter, subcortical gray matter, white matter, cerebrospinal fluid, and pathological tissue) with FSL function *5ttgen* and then registered to the diffusion image with six degrees-of-freedom (DOF) using *flirt* tool from FSL (Greve & Fischl, 2009; Jenkinson et al., 2002). A streamline seeding mask was defined between the interface of white matter and gray matter using *the 5ttgmwmi* function. Hereafter, *the-act* and *-backtrack* options were adopted with *the tckgen* function to anatomically constrain the extent of streamline propagation and allow resampling of a streamline if it terminates in an unexpected location (e.g., cerebrospinal fluid). A probabilistic tractography algorithm was used, as it integrates fibre orientation distribution (FODs) more accurately by considering second-order tensor information, leading to improved streamline propagation (Tournier et al., 2010). For each participant, 10 million tracts were generated with a specified maximum tract length (250 mm) and cut-off FOD amplitude (0.06) from an iterative procedure with random placement of seeds within the mask. These tracts were further refined by using *tcksift2* function, which determines appropriate cross-sectional area multipliers for each streamline. This algorithm optimizes the streamline weights to ensure that the reconstructed fibre volumes correspond accurately with those estimated from the underlying diffusion signal, thereby providing a more reliable and accurate measure of fibre connectivity (Smith et al., 2015). Moreover, the tractograms were visually inspected to ensure comprehensive coverage of the white matter and to confirm that the streamlines were confined within the white matter boundaries.

### 2.11 Structural connectome

Each participant’s anatomical image was initially segmented into distinct brain regions utilizing the same atlas employed during the functional connectivity analysis (Desikan et al., 2006), with *the recon-all* command (Fischl et al., 2004). The atlas-based parcellation image, along with streamline weights and tractography data, was then utilized to generate a symmetric connectivity matrix using the MRtrix function *tck2connectome* (Tournier et al., 2019). The resulting connectivity matrix was scaled with the inverse of node volumes, eliminating biases due to varying brain region sizes and facilitating more accurate and comparable measurements of connectivity strength across different regions (Hagmann et al., 2008). Inter-subject connection density normalization was achieved by adjusting the streamline weights with a proportionality coefficient, such that each matrix entry denotes the total intra-axonal cross-sectional area of the axons associated with the fiber bundle of interest, referred to as the fiber bundle capacity (FBC ; Smith et al., 2022). The resultant SC matrix of 10 LAN-ROIs comprised 10 left-hemispheric SC edges, 10 right-hemispheric SC edges and 25 inter-hemispheric SC edges, yielding a total of 45 LAN SC edges (as represented in Figure 3).

### 2.12 Statistical modeling and testing

In this study, we implemented linear mixed-effects (LMEs) modeling, a widely recognized and robust statistical framework for analyzing neuroimaging and behavioral data, particularly in studies involving repeated measures and hierarchical structures (Chen et al., 2013, 2014; Chua et al., 2015; Jones et al., 2013). This approach effectively controls for Type I errors while maintaining high statistical power (Galbraith et al., 2010; Riha et al., 2020; Scandola & Tidoni, 2021). Unlike traditional linear regression models, LME models account for both fixed effects (e.g., age, functional connectivity (FC), and structural connectivity (SC)) and random effects (e.g., inter-subject variability in gender and education), making them particularly well-suited for the design of the current study. Both gender and education were considered categorical variables; education included 32 distinct levels, each representing a specific educational attainment stage. These levels were derived by assigning a difficulty rating to each education stage and aggregating them across an individual’s academic record (see Section 2.1). As neither gender nor education was the primary variable of interest, they were modelled as random intercepts to appropriately account for group-level variance.

For this analysis, we adopted a model comparison approach in which a full model (including all fixed and random effects) was compared to a null model (including all random effects, but excluding one fixed effect). To determine whether the inclusion of a specific fixed effect significantly improved the model fit, we performed an analysis of variance (ANOVA) to compare the two models, a method that is well supported by previous research in cognitive neuroscience (Riha et al., 2020; Scandola & Tidoni, 2021). Additionally, model fit was further evaluated using the Akaike Information Criterion (AIC) and Bayesian Information Criterion (BIC), with lower values indicating a superior fit (Burnham & Anderson, 2002) and, hence, better predictability. To control for multiple comparisons in the post-hoc analysis, we applied the False Discovery Rate (FDR) correction, reducing the likelihood of Type I errors and ensuring the robustness of p-values (Benjamini & Hochberg, 1995).

All statistical analyses were performed in R version 4.4.1 (https://www.r-project.org/) using the lme4 (Bates et al., 2015) and lmerTest (Kuznetsova et al., 2017) packages. The lme4 package was used to fit all LMEs, while the lmerTest package was employed to provide significance tests for the fixed effects by incorporating Satterthwaite’s approximation of degrees of freedom, ensuring more reliable p-values in the mixed-effects models. Graphical representations of the results were generated using the ggplot2 package (Wickham, 2016), version 3.5.1, and all the nodes and edges of the cLAN were visualized thro ugh schematic brain plots created using BrainNet viewer (Xia et al., 2013).

#### 2.12.1 Behaviour analysis

The behavioral scores for each of the 16 subdomains across the language comprehension (LC) and production (LP) tasks were first standardized using z-score normalization in R, ensuring comparability across tasks. Missing data were handled on a task-specific basis by removing rows with missing values for each respective behavioral task, while preserving data for other tasks. Following data preprocessing, a model comparison approach was implemented for each language behavior (LB) task. The full model, which included age as a fixed effect alongside random effects for gender and education (Equation 2), was compared with a null model that excluded age as a fixed effect (Equation 1). The comparison between these models allowed for the evaluation of age as a significant predictor of language behaviour. P-values from the analysis of variance (ANOVA) were FDR-corrected to statistically compare the estimate of age across each language behavior task.

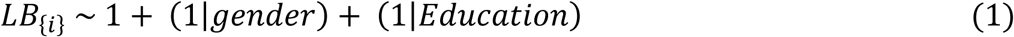

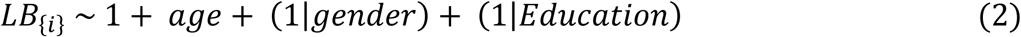

where, “𝑖” indexes each of the 16 language behavior (LB) measures.

Additionally, to test the model robustness, another set of analyses was conducted with gender specified as a fixed effect and compared the resulting model estimates and fit indices to those obtained when gender was modelled as a random effect (Equations 1 and 2).

#### 2.12.2 SC analysis

Prior to analysis, structural connectivity (SC) values from each of the edge within cLAN were standardized using z-score normalization to mitigate potential bias in parameter estimation and enhance model convergence. To investigate age-related differences in intra-hemispheric and inter-hemispheric SCs, we applied a linear mixed-effects (LME) model similar to the approach described earlier. A full model (Equation 4), incorporating age as a fixed effect while controlling for gender and education as random effects, was compared to a null model (Equation 3), which included only the random effects. The models were compared using ANOVA, with the resulting p-values subjected to FDR correction across the 45 LMEs fitted for each SC edge.

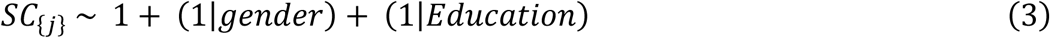

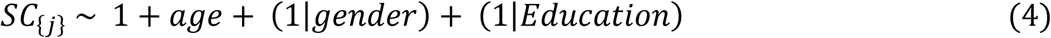

where, “j” indexes each of the 45 SC edges within the cLAN.

#### 2.12.3 FC analysis

Analogous to the SC analysis, normalized FC values for each of the 45 edges were employed to investigate age-related FC variations for all edges. The full model (Equation 6) was compared with the null model (Equation 5) to estimate the effect of age as a predictor of FC, with random effects for gender and education controlled in both models. Statistical comparisons between models were carried out using ANOVA, with FDR correction applied across the 45 LMEs fitted for each FC edge.

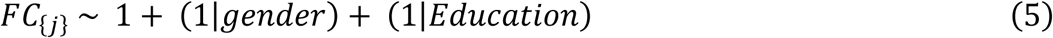

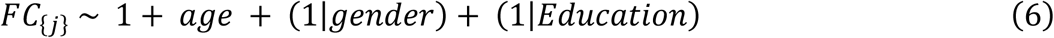

where, “j” indexes each of the 45 FC edges within the cLAN.

#### 2.12.4 Interaction analysis

To assess the effect of brain connectivity (45 SC/FC edges) on the variation in 16 language behaviour (LB) task scores across age, an initial analysis was conducted with the approach, using LME models. However, the model encountered convergence issues, likely due to the complexity of the data and the non-linear relationships between variables. A promising alternative for more flexible modeling is the Generalized Additive Mixed Model (GAMM; Lin & Zhang, 1999; Wood, 2017) because it does not require a priori knowledge of the functional form for modeling language behaviour specific trends with brain connectivity and its interaction with age. Specifically, GAMM substitute the fixed linear slopes with flexible ‘smooth’ functions to capture non-linear relationships and provide more robust estimates that can be applied to complex cross-sectional cognitive data (Roger et al., 2023; Sørensen, Brandmaier, et al., 2021).

In this study, GAMMs were implemented using *bam* function from *mgcv* package in the R library (version 1.9; Wood, 2017). Given that our primary objective was to explore how SC and FC influence language abilities/ behavior (LB) as a function of age, we adopted an interaction-based modeling approach to capture the dynamic interplay between connectivity and behavior across age. This approach utilized tensor-interaction smooth functions (𝑡𝑖()) which enables separation of main effects (i.e., smooth terms “𝑠()” for age and SC/FC individually) from their interaction term, allowing for proper hierarchical model comparisons (Wood, 2011, 2017).

For each of the 16 language behavior (LB) measures, we constructed GAMMs for all 45 SC edges and 45 FC edges individually. Smooth terms for random effects (RE) i.e., gender and education level were included in all models. The final models used in the SC × age analysis was as follows:

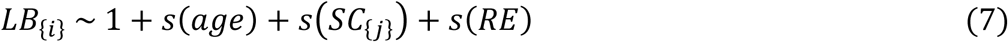

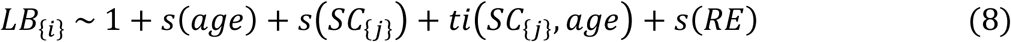

And following were the GAMMs equations used to analyse FC × age interactions:

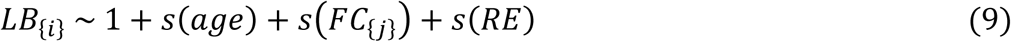

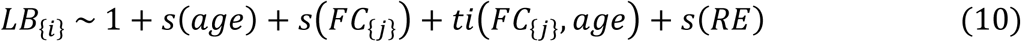

where, “𝑖” indexes each of the 16 language behavior measures, and “𝑗” indexes each of the 45 SC or FC edges analyzed, and RE represents the random effects such as gender, education. Thus, a total of 720 GAMMs (16 behaviors × 45 edges) were fitted for SC and FC analyses, respectively.

To test whether the inclusion of the interaction term significantly improved model fit, we compared the full interaction model to the main effects model using an ANOVA. This direct comparison enables isolation of the unique contribution of the interaction, in line with statistical best practices (Wood, 2017). Given the large number of comparisons (16 language behaviors × 45 SC edges and 16 language behaviors × 45 FC edges), we applied False Discovery Rate (FDR) correction to control for multiple testing. Specifically, for each behavior separately, we performed 45 independent model comparisons (i.e., one per SC or FC edge), and the resulting p-values from these ANOVAs were FDR-corrected across the 45 comparisons. Effects were considered significant at an FDR-adjusted p < 0.05.

## 2. Results

### 3.1 Variation of performance in language comprehension (LC) and production (LP) tasks across lifespan

To explore the impact of age on performance across the 16 subdomains of LC and LP tasks, the age estimates from full linear mixed effects (LME; Equation 2) were plotted with their respective error bars (Figure 1C). As anticipated, vocabulary (vocab) and proverb comprehension demonstrated the strongest positive associations with age, with significant estimates even after adjusting the p-values for multiple comparisons (Vocab: β = 0.017, p adj. < 0.0001; Proverb: β = 0.012, p adj. < 0.0001). However, LP tasks, including Verbal Fluency (VF), five subdomains of the picture priming (RT_low_phon, RT_low_sem, RT_high_phon, RT_high_sem, and High_sem), and the tip-of-the-tongue (ToT), demonstrated significant decline in performance with age (Figure 1C; VF: β = -0.012, p adj. < 0.001; RT_low_phon: β = -0.009, p adj. < 0.001; RT_low_sem: : β = -0.007, p adj. < 0.05; RT_high_phon: β = -0.006, p adj. < 0.05; RT_high_sem: β = -0.006, p adj. < 0.05; High_sem: β = - 0.008, p adj. < 0.01; ToT: β = -0.016, p adj. < 0.001). In contrast to the high semantic similarity condition (High_sem), no significant age-related changes were observed in the accuracy of the high phonological similarity condition in the picture priming task (High_phon: β = -0.003, p adj. = 0.16). Interestingly, performance among both accuracy and response time measures in the sentence comprehension tasks (Syn_comp, Sem_comp, RT_Syn_comp, and RT_Sem_comp) were not affected by age-related variation across the lifespan (Figure 1C; Syn_comp: β = 0.001, p adj. = 0.58; Sem_comp: β = -0.0006, p adj. = 0.82; RT_Syn_comp: β = -0.003, p adj. = 0.26; and RT_Sem_comp: β = -0.001, p adj. = 0.81).

**Figure 1.**
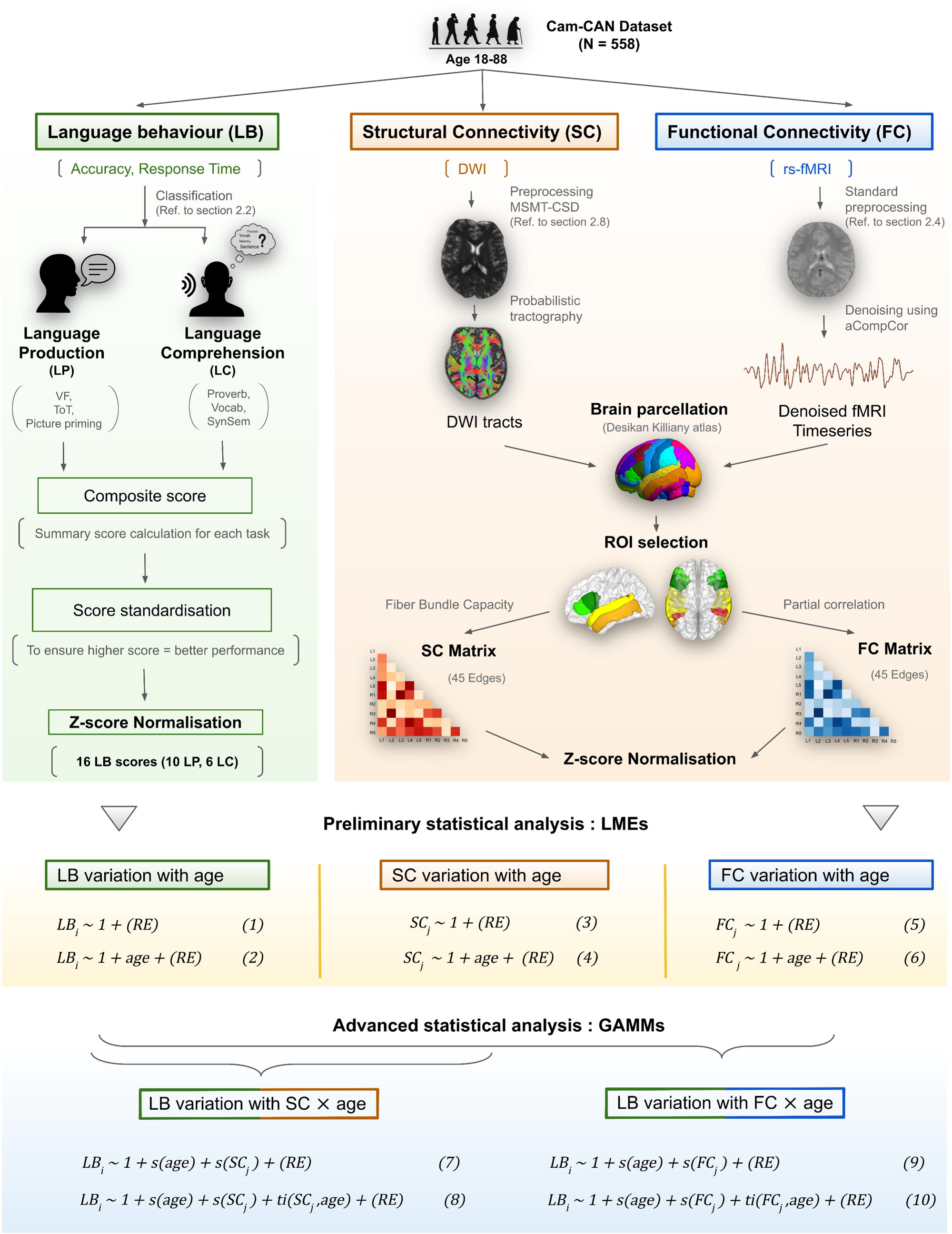
Multimodal framework for investigating age-related language trajectories using language behaviour (LB) and brain connectivity in the Cam-CAN cohort (N=558, ages 18–88 years). The workflow integrates multimodal assessments: (1) z-score normalized language behaviour (LB) scores for production and comprehension tasks; (2) structural connectivity (SC) matrices derived from diffusion MRI after advanced preprocessing and probabilistic tractography; and (3) functional connectivity (FC) matrices computed from denoised resting-state fMRI using partial correlations between core language network ROIs, with Fisher’s r-to-z normalization. Coloured regions in the ROI selection depict core language network regions (for details, see Figure 2). Preliminary analyses used linear mixed-effects models (LMEs) to test age effects on LB, SC, and FC scores, while advanced analyses employed generalized additive mixed models (GAMMs) to capture nonlinear and interaction effects, specifically how edge-specific connectivity modulates age-related language performance. Model equations use subscripts “𝑖” for each of the 16 LB measures, and “𝑗” indexes each of the 45 SC or FC edges analyzed, and RE represents the random effects such as gender, education.

To ensure that these findings were not dependent on model choice, e.g., treating gender as a random effect, we repeated the analyses treating gender as a fixed effect (Equations S1 and S2) and compared the resulting model estimates and performance metrics. As seen in the supplementary tables (Table S1 and S2), the model estimates for age and the overall model fit indices (AIC, log-likelihood, and FDR-adjusted p-values) show negligible differences between the two modeling strategies across all language tasks. These current finding attests to the stability and robustness of the model results, regardless of how gender is parameterized (Meteyard & Davies, 2020; Schielzeth et al., 2020). Hence, for the subsequent analyses gender is treated as a random effect.

### 3.2 Age-related variation in structure connectivity (SC) of core language network (cLAN)

The relationship between structural connectivity within the cLAN and age was visualized using schematic brain plots and horizontal plots (Figure 3) based on the age-related beta coefficients from full linear mixed effects (LME) model (Equation 4). Examination of intra-hemispheric structural connectivity patterns revealed significant age-related alterations in both cerebral hemispheres, encompassing both increases and decreases in connectivity. Interestingly, a greater number of left-hemispheric SCs showed significant negative association with age (Figure 3A; No. of connections = 6/10; p adj. < 0.05) as compared to the right-hemispheric SCs (Figure 3B; No. of connections = 3/10; p adj. < 0.05). The connection strength within the right Broca’s area (RParsO_RParsT) showed a significant increase with age (β = 0.006, p adj. < 0.01), whereas the SC between the left Pars-opercularis (LParsO) and left Pars-triangularis (LParsT) demonstrated a significant age-related decrease (β = -0.005, p adj. < 0.05). Additionally, most intra-hemispheric fronto-temporal connections displayed a general pattern of age-related decline, including connections between Broca’s area and the superior temporal sulcus (LSTS_LParsO: β = -0.012, p adj. < 0.001, LSTS_LParsT: β = -0.009, p adj. < 0.001, and RSTS_RParsT: β = -0.007, p adj. < 0.01), as well as the middle temporal gyrus (LMTG_LParsO: β = -0.021, p adj. < 0.001, LMTG_LParsT: β = -0.018, p adj. < 0.001, RMTG_RParsO: β = -0.01, p adj. < 0.001, and RMTG_RParsT: β = -0.016, p adj. < 0.001). Contrary to this overall trend of age-related decline, the structural strength of intra-hemispheric temporo-temporal connections showed significant positive association with age, involving the STS, MTG, and STG among both hemispheres (LSTS_LMTG: β = 0.019, p adj. < 0.001, LSTS_LSTG: β = 0.014, p adj. < 0.001, LMTG_LSTG: β = 0.014, p adj. < 0.001, RSTS_RMTG: β = 0.012, p adj. < 0.001, RSTS_RSTG: β = 0.016, p adj. < 0.001, RMTG_RSTG: β = 0.004, p adj. < 0.05).

**Figure 2.**
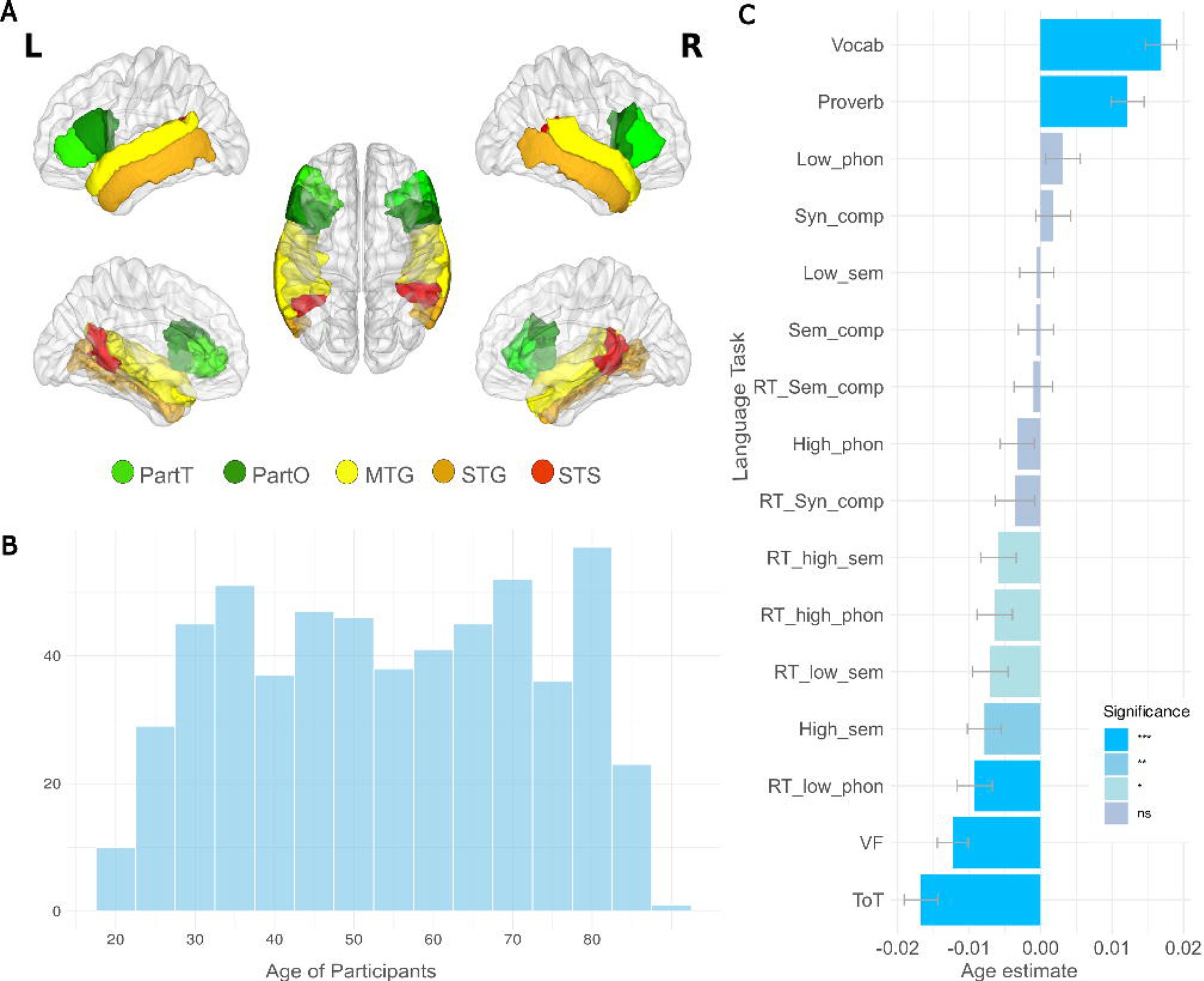
A, Three-dimensional representation of core language network (cLAN) ROIs (Regions of Interest) in both hemispheres, shown from medial and lateral views. B, Distribution of age (18-88) among 558 CamCAN participants. C, Age-related variation in language task performance depicted through horizontal bar plots. Each bar represents the beta coefficient for age across different language comprehension and production tasks derived from the Linear mixed model (LME; Equation 2), color-coded based on significance levels. Abbreviations: ParsO (Pars Opercularis), ParsT (Pars Triangularis), MTG (Middle Temporal Gyrus), STG (Superior Temporal Gyrus), STS (Superior Temporal Sulcus), Vocab (Vocabulary), ToT (Tip of the Tongue), VF (Verbal Fluency), High_phon (High Phonetic Similarity), High_sem (High Semantic Similarity), Low_phon (Low Phonetic Similarity), Low_sem (Low Semantic Similarity), RT (Response Time), Syn_comp (Syntactically Complex), and Sem_comp (Semantically Complex).

Regarding inter-hemispheric SCs, although a few connections remained stable across age (Figure 3C; No. of connections = 6/25: LMTG_RSTG: β = -0.0003, p adj. = 0.89, LMTG_RMTG: β = -0.0003, p adj. = 0.16, LSTG_RMTG: β = 0.0046, p adj. = 0.059, LSTS_RParsT: β = -0.0016, p adj. = 0.56, LParsT_RSTS: β = -0.0035, p adj. = 0.11, LParsT_RSTG: β = -0.0039, p adj. = 0.10), the majority showed significant age-related changes (Figure 3C; No. of connections = 19/25; p adj. < 0.05). Two primary patterns were readily observable among the notable age-related variations in the inter-hemispheric SCs. One of the trends consisted a significant decline in the SC of fronto-temporal (LParsO_RMTG: β = -0.014, p adj. < 0.001, LParsO_RSTG: β = -0.012, p adj. < 0.001, LParsO_RSTS: β = -0.008, p adj. < 0.001, LParsT_RMTG: β = -0.006, p adj. < 0.05, LMTG_RParsO: β = -0.014, p adj. < 0.001, LSTG_RParsO: β = -0.008, p adj. < 0.001, LMTG_RParsT: β = -0.007, p adj. < 0.01, LSTS_RParsO: β = -0.006, p adj. < 0.05, LSTG_RParsT: β = -0.005, p adj. < 0.05) and fronto-frontal connections (LParsO_RParsO: β = -0.015, p adj. < 0.001, LParsO_RParsT: β = -0.014, p adj. < 0.001, LParsT_RParsO: β = -0.011, p adj. < 0.001, LParsT_RParsT: β = -0.006, p adj. < 0.05). Conversely, the inter-hemispheric temporo-temporal showed significant strengthening with age, particularly between the left and right hemispheric regions of the STS, MTG, and STG (LSTS_RTS: β = 0.011, p adj. < 0.001, LSTS_RMTG: β = 0.008, p adj. < 0.001, LSTS_RSTG: β = 0.010, p adj. < 0.001, LMTG_RSTS: β = 0.005, p adj. < 0.05, LSTG_RSTG: β = 0.006, p adj. < 0.01, and LSTG_RSTS: β = 0.012, p adj. < 0.001).

**Figure 3.**
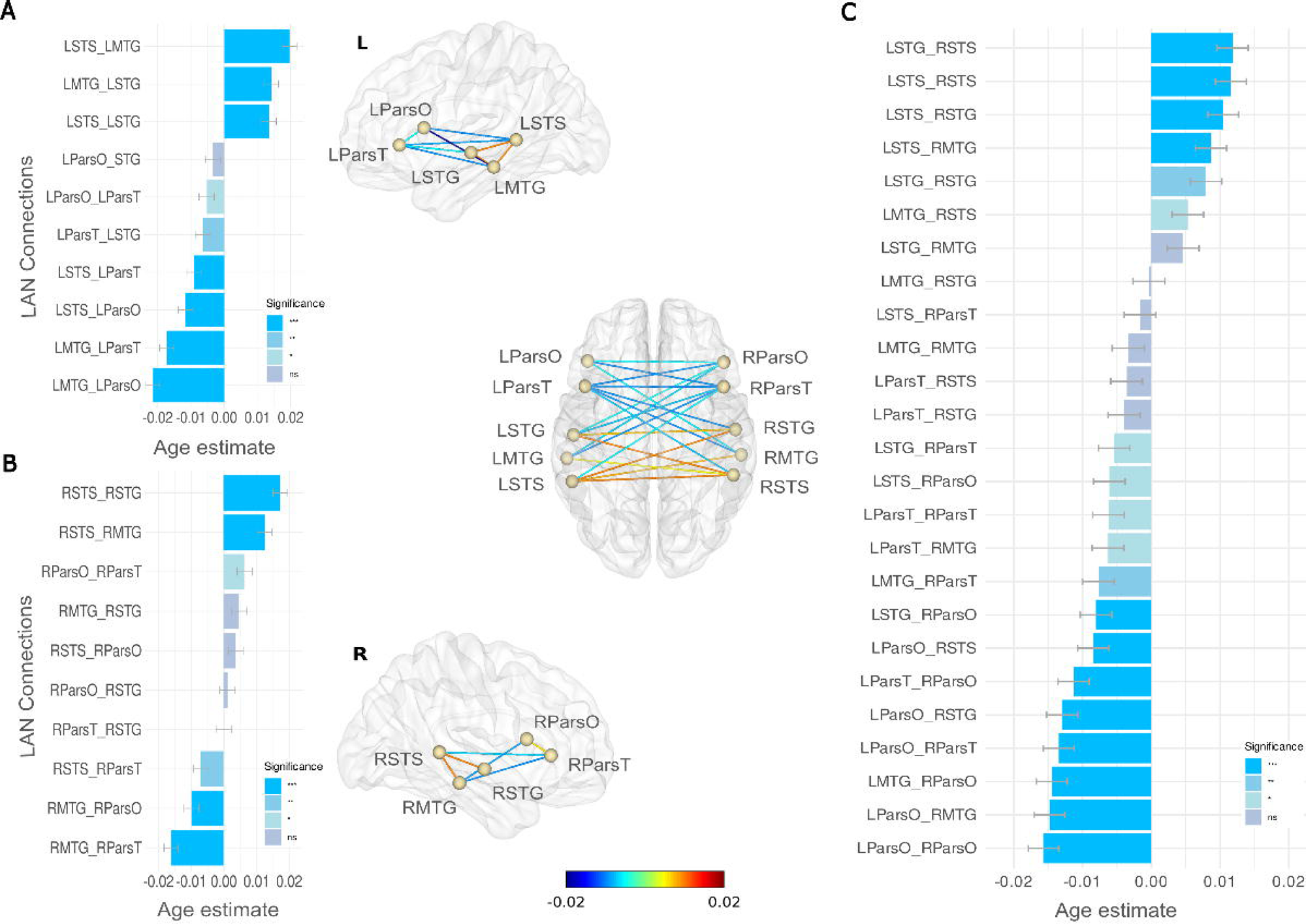
Age-related variation in structural connectivity within the language network across intra-and inter-hemispheric pathways. Beta estimates from linear mixed-effects models (LMEs; Equation 4) are visualized for 45 cLAN edges to quantify the effect of age on structural connectivity (SC). Panels A–C represent results for left-hemispheric (A), right-hemispheric (B), and inter-hemispheric (C) connections, respectively. Each horizontal bar corresponds to an individual edge, with the x-axis reflecting the fixed-effect estimates of age on SC strength. Significance levels are indicated by bar shading: dark blue for p < 0.001, medium blue for p < 0.01, light blue for p < 0.05, and grey for non-significant results (FDR-corrected). Central schematics represent the spatial distribution of connections across the cLAN, with edges coloured according to the signed beta coefficient, consistent with the colour bar scale. Labels “L” and “R” denote left and right hemispheres, respectively. All region abbreviations are consistent with those presented in Figure 2A.

**Figure 4.**
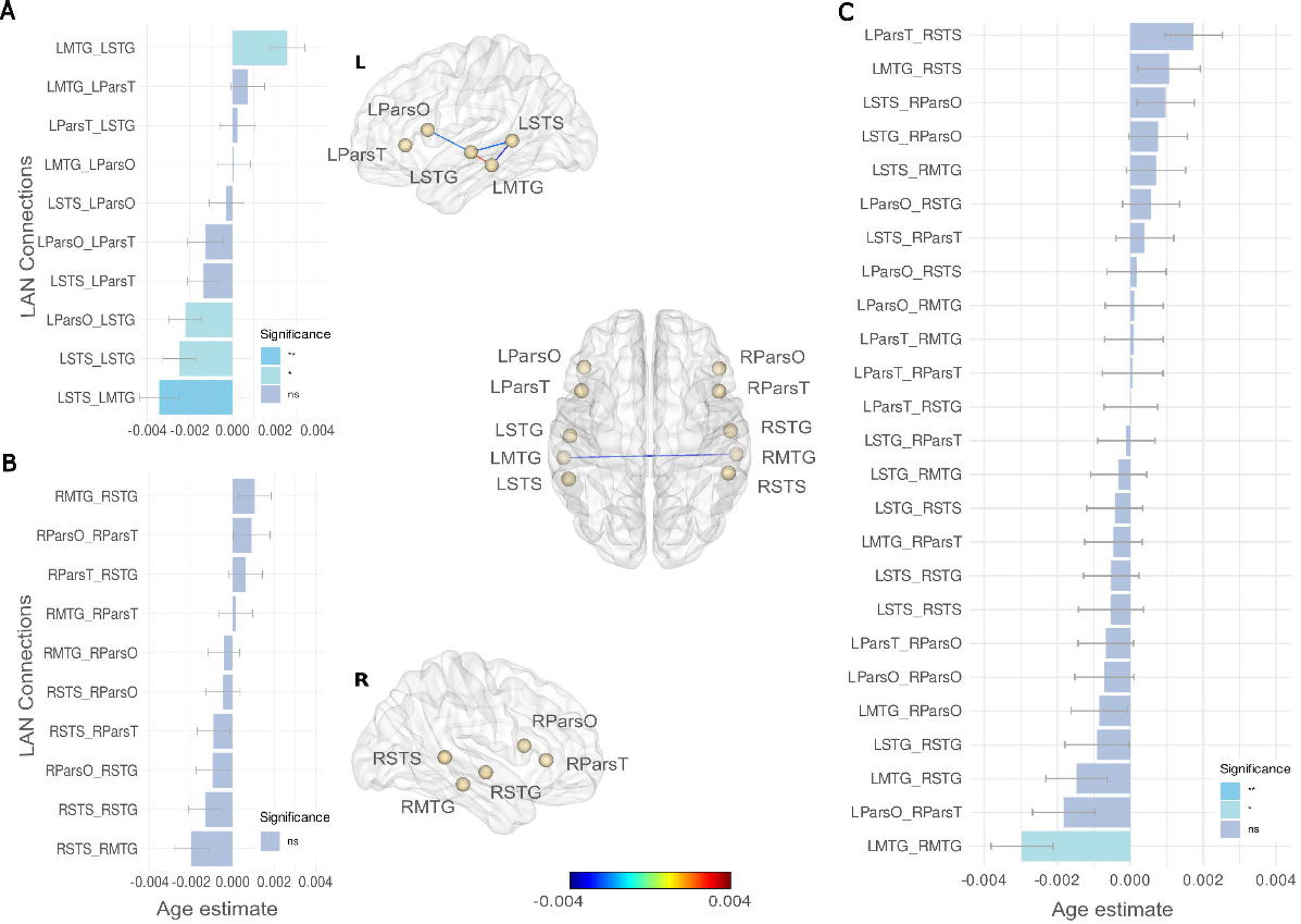
Age-related variation in functional connectivity within the language network across intra- and inter-hemispheric pathways. Panels A–C display fixed-effect estimates for age on functional connectivity (FC) within the cLAN, derived from linear mixed-effects modeling (LMEs; Equation 6). Panel A shows left-hemispheric connections, Panel B right-hemispheric connections, and Panel C inter-hemispheric connections. Bars represent the magnitude and direction of the age coefficient for each cLAN edge. Significance levels are color-coded following FDR correction: dark blue = p < 0.001, medium blue = p < 0.01, light blue = p < 0.05, grey = non-significant. Central schematic brain plots provide spatial context for each edge, with edges coloured by beta coefficient magnitude and direction as indicated by the continuous scale below. Labels “L” and “R” denote left and right hemispheres, respectively. All region abbreviations are consistent with those presented in Figure 2A.

### 3.3 Age-related variation in functional connectivity (FC) of cLAN

The age estimates derived from the full LME model for FC (Equation 6) were utilized to investigate the edge-specific variation in functional connectivities within the cLAN across the lifespan (Figure 3). Linear mixed effects (LME) modeling revealed distinct patterns of age-related changes in FC within the intra- and inter-hemispheric connections of the cLAN. In the case of intra-hemispheric FCs, connections within the left hemisphere demonstrated both significant negative (Figure 3A; No. of connections = 3/10: LSTS–LMTG, LSTS–LSTG, and LParsO_LSTG; p adj. < 0.05) and positive (Figure 3A; No. of connections = 1/10: LMTG_LSTG; p adj. < 0.05) associations with age, whereas no significant age-related associations were found for right-hemispheric functional connections (Figure 3B; No. of connections = 0/10; p adj. > 0.05). Within left-hemispheric FCs, among temporo-temporal connections, FC associated with STS region showed significant decline with age (LSTS–LMTG: β = - 0.0039, p adj. < 0.001; LSTS–LSTG: β = -0.0023, p adj. < 0.01), whereas FC between MTG and STG demonstrated a significant positive association with age (β = 0.0023, p adj. < 0.01). Considering inter-hemispheric FCs, most connections remained unaffected by age (Figure 3C; No. of connections = 24/25; p adj. > 0.05). However, a significant age-related decrease was observed in the FC between the left and right middle temporal gyri (MTG) (β = -0.003, p adj. < 0.05).

### 3.4 Non-linear relationship between edge-specific structural connectivity (SC) and age-related language variability

To examine the role of edge-specific SC in modulating the relationship between language behavior and age, a tensor interaction smooth term (𝑡𝑖()) was incorporated into the generalized additive mixed model (GAMM; Equation 8) independently for each of the 16 language behavior tasks and 45 SC edges, resulting in a total of 720 GAMMs. For each combination, the full interaction model (Equation 8) was compared with the main effects model (Equation 7) using ANOVA to assess whether including the SC × age interaction significantly improved model fit over an additive model (SC + age). For each language behavior task, the resulting p-values from the 45 edge-wise comparisons were FDR-corrected. Comprehensive model performance metrics, including effective degree of freedom (edf), adjusted R², AIC, F-statistic, and FDR-adjusted p-values were summarized in Table S3. The resulting F-values from the ANOVA model comparisons are displayed as a heatmap (Figure 5A), with each cell corresponding to a specific pair of language behavior task and SC edge. Grey cells indicate non-significant effects (p adj. ≥ 0.05), while blue cells mark significant SC × age interactions after FDR correction. The intensity of the blue colour reflects the magnitude of the F-value, with darker shades representing stronger interaction effects.

The heatmap revealed a sparse yet specific pattern of significant non-linear SC × age interactions, with 9 out of 720 comparisons reaching significance (p adj. > 0) across all language tasks. Majority of these significant effects emerged in language comprehension (LC) tasks (e.g., Syn_comp: 6/9), compared to language production (LP) tasks (e.g., VF: 1/9; RT_high_sem: 2/9). Within LC tasks, age-related modulation of performance in the comprehension of syntactically complex sentences was significantly associated with structural changes in few left-hemispheric and several inter-hemispheric connections (e.g., Syn_comp: LParsT_LSTG, F = 3.55, p adj. < 0.01, LParsO_RParsT, F = 3.90, p adj. < 0.1, LMTG_RMTG, F = 8.62, p adj. < 0.01, LMTG_RParsT, F = 2.78, p adj. < 0.01, LSTG_RMTG, F = 3.46, p adj. < 0.01, LSTG_RSTS, F = 3.19, p adj. < 0.01). In contrast, age-related variation in reaction time of picture priming task high semantic similarity condition (RT_high_sem) was significantly associated with structural strength of temporo-temporal connections, spanning both right-hemispheric (RSTS_RSTG, F = 3.25, p adj. < 0.1) and inter-hemispheric pathways (LSTS_RMTG, F = 4.46, p adj. < 0.01). Additionally, performance in verbal fluency task (VF) task was significantly modulated by structural strength of fronto-temporal inter-hemispheric connection (LParsT_RMTG, F = 7.24, p adj. < 0.01).

The significant interactions identified through ANOVA were further explored using GAMM trajectory plots (Figure 6B), where predicted behavioral trajectories were plotted across age for three SC quantiles (0.25, 0.50, and 0.75), highlighting differential effects of SC on language performance across the lifespan. Our findings revealed a differential role of SC in modulating age-related trajectories of LC and LP tasks, where SC influenced comprehension performance consistently across all ages, 18-88 years (Figure 6A), while its effect on production tasks (Figure 6B) became strongly evident in late adulthood, 60–80 years. Within LC tasks (Syn_comp), higher SC among all significant edges were associated with better performance particularly at older age (Figure 5A). Whereas at younger age, SC exhibited differential effect, some edges supported the syntactic comprehension (LMTG_RParsT, LParsT_LSTG) while other edges (LSTG_RSTS, LSTS_ RMTG, LParsO_RParsT) displayed negative association with performance. However, for production tasks (VF, RT_high_sem), SC effects emerged only in older adults, 70-88 years, with higher SC (LSTS_RMTG, LParsT_RMTG) associated with better performance (Figure 6B).

**Figure 5.**
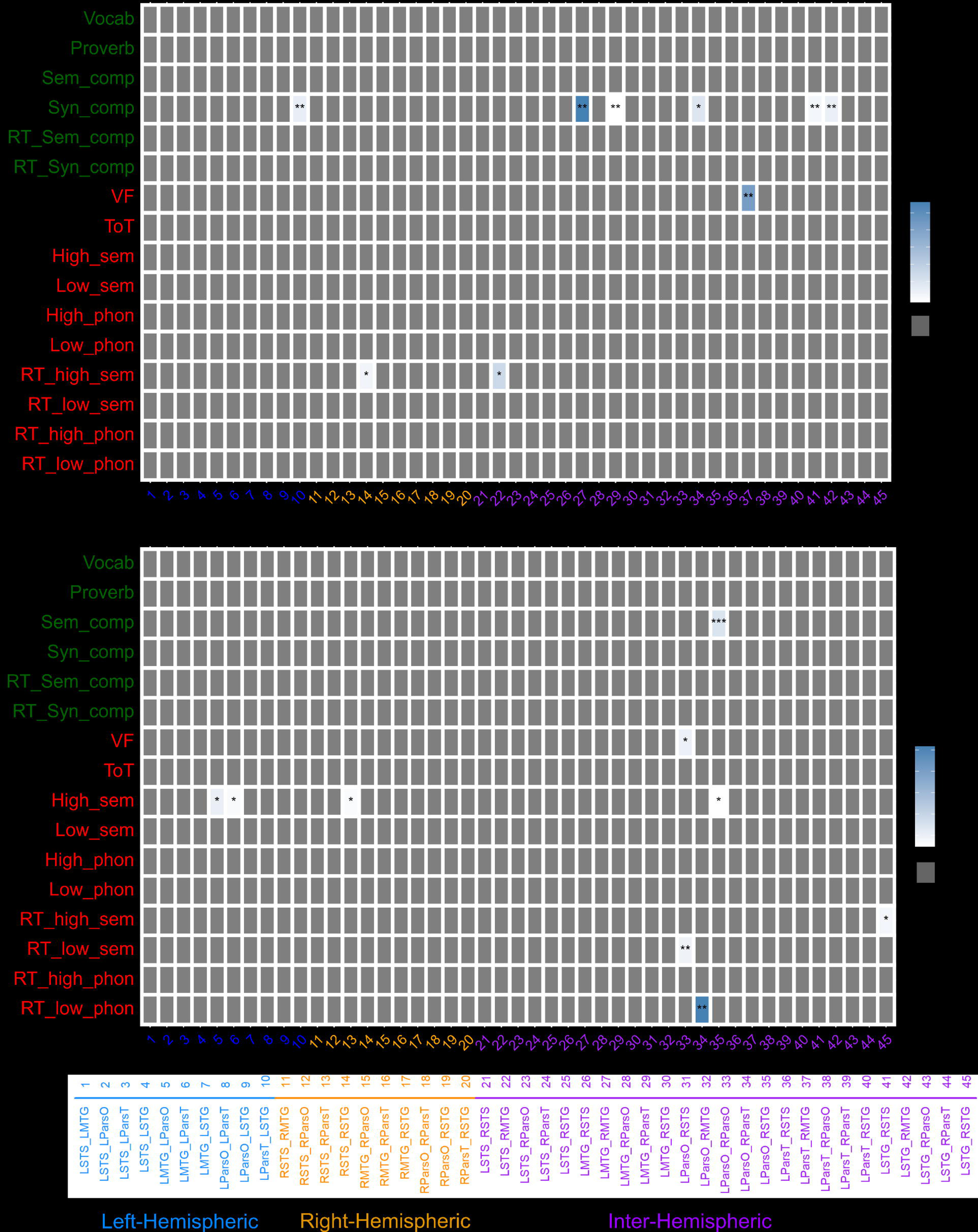
Edge-wise structural (SC) and functional connectivity (FC) × age interactions differentially modulate language comprehension (LC) and language production (LP) across the lifespan. Heatmaps display F-value statistics from Generalized Additive Mixed Models (GAMMs; equations 7-10) based ANOVA comparisons, capturing significant SC × age (A) and FC × age (B) interaction effects across 45 core language network (cLAN) edges and 16 language tasks. Blue squares indicate significant interaction effects (p adj. < 0.05), while grey squares represent non-significant interactions. Language tasks are grouped by domain: 6 LC tasks (green), 10 LP tasks (red); cLAN edges are grouped as left-hemispheric (blue), right-hemispheric (orange), or inter-hemispheric (purple, see legend). Edge abbreviations are provided in Figure 2A.

**Figure 6.**
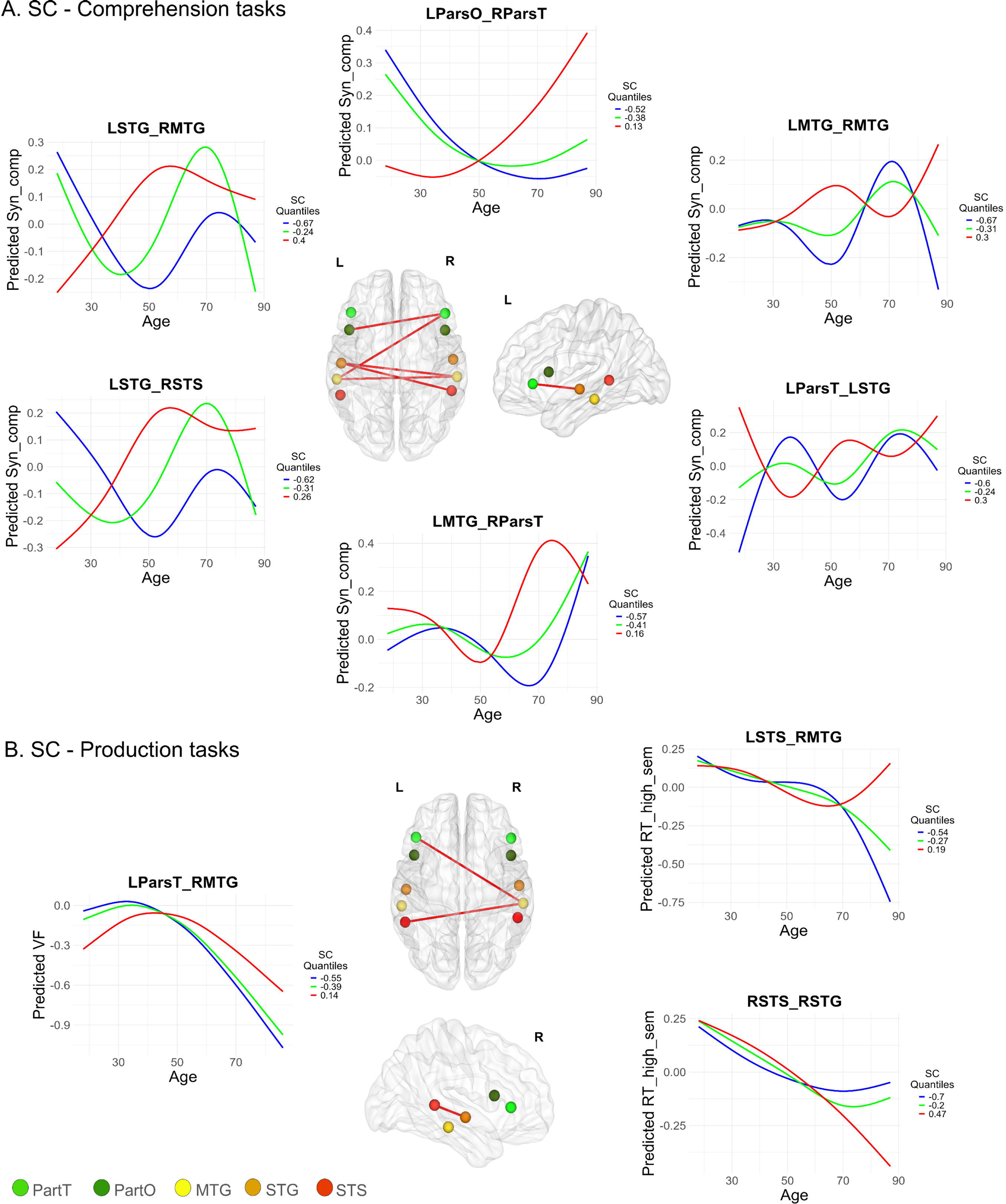
Differential lifespan modulation of language comprehension (LC) and production (LP) by edge-specific structural connectivity (SC) in the core language network (cLAN). This figure presents predicted behavioral trajectories as estimated from Generalized Additive Mixed Models (GAMMs) for representative SC × age interactions, as identified via FDR-corrected post-hoc ANOVA comparisons (see Figure 5). Panel A presents comprehension tasks (Syn_comp) and panel B production tasks (VF, RT_high_sem), demonstrating how language performance varies across the adult lifespan (18–88 years) at three quantiles of SC strength: low (0.25, red), medium (0.50, green), and high (0.75, blue). Notably, SC effects on LC tasks (Panel A) were observed consistently across all ages, whereas modulation of LP tasks (Panel B) by SC emerged predominantly in late adulthood (60–80 years). Accompanying brain schematics depict anatomical nodes and connections, color-coded by region as previously defined in Figure 2A.

### 3.5 Non-linear relationship between edge-specific functional connectivity (FC) and age-related language variability

The role of intra- and inter-hemispheric FCs in non-linear modulation of language performance across age was investigated with the help of an interaction term (FC × age) that was incorporated in the generalized additive mixed model (GAMM; Equation 10). To specifically assess the added explanatory value of the interaction term (FC × age), the full interaction model was systematically compared against the corresponding additive (FC + age) GAMM model (Equation 9), using ANOVA for each of the 16 language behavior (LB) tasks across all 45 core language network (cLAN) FC edges, resulting in a total of 720 model comparisons. To appropriately control for the risk of Type I error due to multiple comparisons, FDR correction was applied to the ANOVA-derived p-values for each set of 45 comparisons within each behavioral measure. Only interaction effects that survived FDR correction (p adj. < 0.05) were considered statistically significant. All GAMMs model performance metrics, including effective degree of freedom (edf), adjusted R², AIC, F-statistic, and FDR-adjusted p-values were summarized in Table S4. The F-value statistics from the ANOVA (FC × age interaction) comparisons are visualized in the heatmap (Figure 4B), where each cell represents a specific task-edge pair (total 16×45=720 comparisons). Significant interactions are depicted in a blue colour gradient with asterisks indicating the FDR-corrected significance level, whereas non-significant results are shaded in grey.

Notably, only a subset of the modelled interactions reached significance after FDR correction. Significant FC × age interactions were more prevalent among language production (LP) tasks (e.g., VF, High_sem, RT_high_sem, RT_low_sem, RT_low_phon) than among language comprehension (LC) tasks (e.g., Sem_comp). Within LP tasks, most significant FC × age interaction effects were identified for inter-hemispheric connections. For example, verbal fluency (VF) performance was significantly associated with age-related variability in the fronto-frontal inter-hemispheric FC between the left and right pars opercularis (LParsO_RParsO, F = 4.91, p adj. < 0.1), and response time measures on picture-priming tasks also showed significant FC × age interactions along inter-hemispheric edges only (RT_high_sem: LSTG_RSTG, F = 3.92, p adj. < 0.1; RT_low_sem: LParsO_RParsO, F = 4.48, p adj. < 0.01; RT_low_phon: LParsO_RParsT, F = 25.8, p adj. < 0.01). In contrast, accuracy in picture priming task with high semantic similarity was significantly associated with FC × age interactions encompasses both intra-hemispheric connections (High_sem: LMTG_RParsO, F = 5.20, p adj. < 0.1; LMTG_RParsT, F = 3.26, p adj. < 0.1; RSTS_RParsT, F = 3.09, p adj. < 0.1) and inter-hemispheric FCs (High_sem: LParsO_RSTG, F = 2.33, p adj. < 0.1). Whereas within LC tasks, only semantic comprehension (Sem_comp) task was significantly modulated by age-related variation within fronto-temporal inter-hemispheric connection (LParsO_RSTG, F = 7.24, p adj. < 0.001).

To characterize the nature of the significant FC × age interactions identified through ANOVA, we generated GAMM trajectory plots for each task-edge pair that survived FDR correction (Figure 7). In these models, predicted behavioral scores were plotted across the adult lifespan for three FC quantiles (0.25, 0.50, and 0.75), each represented by a distinct colour. These findings underscore the differential age-related modulation of LC (Figure 7A) and LP (Figure 7B) tasks by each specific FC edge. For comprehension tasks (Sem_comp), GAMM trajectory indicated that lower inter-hemispheric fronto-temporal FC (LParsO_RSTG) was associated with higher predicted comprehension scores in older adults, whereas higher FC (red line; 0.75 quantile) was associated with relatively stable scores across the lifespan. In the language production (LP) domain (Figure 7B), significant FC × age interaction effects included both intra- and inter-hemispheric connections, specifically for the picture-priming task with high-semantic similarity (High_sem: LMTG_LParsT, LMTG_LParsO, RSTS_RSTG, LParsO_RSTG). Both accuracy and response time in picture priming task (High_sem, RT_low_sem, RT_High_sem, RT_low_phon) displayed a consistent modulation pattern across all FC edges where the effects of higher and lower FC reversed near middle age (40-60 years). In several task-edge pairs (e.g., High_sem: LMTG_LParsO; RT_high_sem: LSTG_RSTG), higher FC was associated with better performance at younger ages (20–40 years), but this association declined or reversed in older adulthood, where lower FC became associated with better performance. Conversely, for other task-edge pairs (e.g., RT_low_phon: LParsO_RParsO; High_sem: LParsO_RSTG), higher FC predicted better performance predominantly in late adulthood (70–88 years) only. Additionally, performance in the verbal fluency task (VF) were associated with higher inter-hemispheric FC (LParsO_RParsO) exclusively in younger adults (18–40 years).

**Figure 7.**
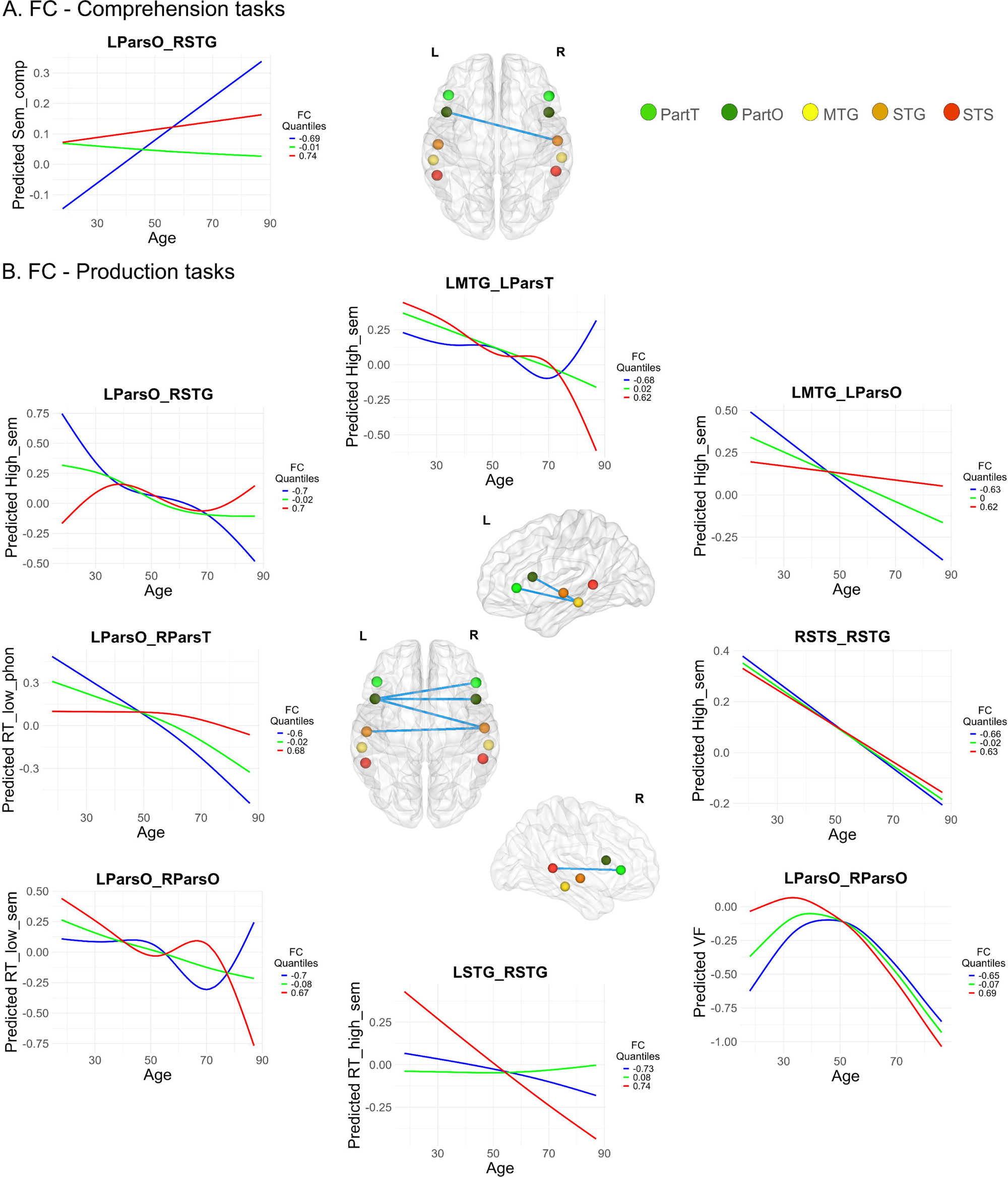
Differential lifespan modulation of language comprehension (LC) and production (LP) by edge-specific functional connectivity (FC) in the core language network (cLAN). Analogous to Figure 6, this figure illustrates predicted behavioral trajectories, estimated from Generalized Additive Mixed Models (GAMMs), for representative FC × age interactions identified via FDR-corrected post-hoc ANOVA (see Figure 5). Panel A presents comprehension tasks (Sem_comp), while Panel B shows production tasks (VF, High_sem, RT_high_sem, RT_low_sem, and RT_low_phon). Trajectories depict model-predicted, z-standardized behavioral outcomes across adulthood (18–88 years) at three FC quantiles: low (0.25, red), medium (0.50, green), and high (0.75, blue). Notably, age-dependent shifts and reversals in FC effects were observed: inter-hemispheric FC played a primary role in modulating LC performance, whereas both intra- and inter-hemispheric cLAN edges contributed to LP performance, revealing distinct FC dynamics across domains and ages. Brain schematics indicate relevant anatomical regions and connections, using consistent color-coding as in Figure 2A.

## 3. Discussion

Aligning with previous research, the present study reinforces language production (LP) often declines, relatively language comprehension (LC) is preserved or even improved (Baciu et al., 2021; Burke & Shafto, 2004; Diaz et al., 2016; Goral et al., 2011; Kemper et al., 2004; H. Zhang & Diaz, 2023a). Previous studies (Davis et al., 2009b; Ferré et al., 2019; Fletcher et al., 2018; Gertel et al., 2020), reported increase in resting-state functional connectivity (rsFC) in tandem with white matter loss and cortical thinning giving rise to conflicting theories of compensation via specialized networks that serve as cognitive reserve (Cabeza et al., 2018) and contrarily, through reduction in specialized processing, also referred to as de-differentiation (Zhang & Diaz, 2023a). Here, rsFC, structural connectivity (SC), and behavioral data across a large lifespan sample is integrated using non-linear Generalized additive mixed models (GAMM). Structural decline in fronto-temporal and within-frontal connections alongside strengthening within temporal lobe connections was observed in core language network (cLAN) regions (Figure 3). FC within cLAN was largely preserved, with limited left- and inter-hemispheric decline (Figure 4). GAMM analyses suggested that neurocognitive aging is not a uniform process but reflects strategic network adaptations with differentially modulating comprehension versus production abilities across lifespan (Figure 5, 6, 7). The implications of these findings are discussed in detail in following discussions.

### Heterogenous language aging

LP tasks, including VF, picture priming and Tip of tongue (ToT) demonstrated significantly reduced accuracy and prolonged response times with advancing age (Figure 1C), possibly driven by weakened connections between semantic and phonological representations, reduced inhibitory control, and slower processing speeds, which collectively impair efficient lexical retrieval (Baciu et al., 2021; Diaz et al., 2014, 2016; Shafto et al., 2007). While inhibitory control and other executive functions likely contribute to the observed decline in language production tasks, these tasks inherently involve multiple cognitive processes, including working memory and executive control. Future studies incorporating production tasks with varying executive demands and independent assessments of executive functions would be valuable to disentangle the specific contributions of language production mechanisms versus domain-general cognitive decline across aging (Martin et al., 2022).

LC performance got preserved (sentence comprehension) or enhanced with age (vocabulary and proverb comprehension). This also aligns with the cognitive reserve hypothesis, which suggests that accumulated knowledge and experience can compensate for age-related neural decline (Cabeza et al., 2018). Although LC tasks assessed here, including syntactically and semantically complex sentence comprehension, impose substantial executive and working memory demands, they also show resilience to age-related decline. These findings indicate that the differential vulnerability between LP and LC cannot be explained purely by differences in executive or working memory load or the crystallized versus fluid intelligence dichotomy (Diaz et al., 2016). Instead, such resilience may reflect domain-specific preservation supported by compensatory neural mechanisms (Shafto et al., 2020).

### Frontal vulnerability and temporal resilience

Fronto-temporal intrahemispheric connections exhibited bilateral SC reductions slightly more in left hemisphere compared to right possibly explaining increased right-hemispheric involvement during language comprehension (Geva et al., 2012; Peelle et al., 2010, 2013; Tyler et al., 2010) (Figure 3). Left-hemispheric fronto-frontal connections (LParsO-LParsT) also decline, aligning with “The frontal aging” hypothesis (Greenwood, 2000), suggesting frontal regions may be more susceptible to age-related degeneration as interetable from reduced fractional anisotropy (FA) and frontal atrophy in older adults (Jao et al., 2020; Madden et al., 2012; Pfefferbaum et al., 2005a). Conversely, SC within temporal regions were relatively preserved or strengthened, consistent with minimal FA changes and increased intra-temporal connectivity among elderly (Jao et al., 2020; Pfefferbaum et al., 2005b; Salat et al., 2005). Furthermore, the average strength of the right temporal module increased with age (Madden et al., 2020), supporting the anterior-posterior gradient of white matter decline (Sullivan et al., 2010), and the “last-in-first-out” model (Jao et al., 2020; Madden et al., 2012; Salat et al., 2005; Yang et al., 2014).

On the other hand, inter-hemispheric SCs declined in fronto-frontal and fronto-temporal connections, while temporal-temporal connections were preserved or enhanced. highlighting the vulnerability of frontal regions Such reorganization aligns with widely recognized pattern of anterior-to-posterior degeneration of corpus callosum (Bastin et al., 2010; Hou & Pakkenberg, 2012; Kochunov et al., 2007; Salat et al., 2005; Sullivan et al., 2010), given that the anterior corpus callosum supports fronto-frontal and fronto-temporal connectivity, and the posterior corpus callosum connects bilateral temporal lobes (De Benedictis et al., 2016; Goldstein et al., 2025; Huang et al., 2005; Wang et al., 2020).

In contrast to SC, FC demonstrated a relative age-related resilience among cLAN regions (Figure 4) accompanied by selective declines between LSTS-LSTG, LSTS-LMTG, and LParsO-LSTG. The increased FC between LMTG and LSTG may be supported by enhanced SC between left temporal regions observed in this study. FC aging mimiced the asymmetrical trajectory of SC, with significant reorganization in the left hemisphere and minimal changes in the right, suggesting right-hemispheric resilience, consistent with Rafiq et al., (2022). The inter-hemispheric FC demonstrated age-related stability, with few observed age-related reorganizations (Figure 4C). Specifically, preservation of fronto-temporal FC despite SC decline may reflect compensatory mechanisms maintaining functional integrity across lifespan (e.g., Pistono et al., (2021)) .

### Selective connectivity reorganization explains divergent language aging

GAMM analysis (Sørensen et al., 2021) revealed only a minority of possible SC × age and FC × age interactions significantly modulated language behaviour (LB) outcomes, highlighting that compensatory mechanisms are highly selective (Billot et al., 2024; Scheller et al., 2014; Zhang et al., 2014). Essentially, specific pathways are adaptively leveraged to preserve function (Fitzhugh et al., 2019; Rafiq et al., 2022; Rizio & Diaz, 2016). This aligns with the targeted compensation framework, in which network flexibility relies on functionally relevant structural highways (Park & Reuter-Lorenz, 2009; Reuter-Lorenz & Cappell, 2008).

Left-hemispheric dominace of fronto-temporal pathways in SC X age interactions for LC (Figure 5A, 6A) is consistent with the critical role of left fronto-temporal regions in syntax (Den Ouden et al., 2019; Price, 2010; Wilson et al., 2011) and with evidence for bilateral reorganization in aging (Tyler et al., 2010; Friederici, 2011; Friederici et al., 2007). Complementing these observations, Antonenko et al., (2013) demonstrated that integrity of the uncinate fasciculus and forceps minor support syntactic comprehension with advanced age. Further, the trajectory analyses revealed that stronger left-hemispheric and inter-hemispheric fronto-frontal/ fronto-temporal connections predicted better syntactic comprehension in elderly. Whereas, in young, effects were mixed, some fronto–temporal pathways enhanced comprehension, while others showed negative associations (Figure 6A), suggesting that when the system operates near peak efficiency, added reliance on alternative routes may introduce interference (Naeser et al., 2011). The same pathways may become supportive in elderly, consistent with scaffolding and CRUNCH models (Park & Reuter-Lorenz, 2009; Reuter-Lorenz & Cappell, 2008), highlighting a dynamic shift from inefficiency to compensation across the lifespan.

In LP tasks (VF, RT_High_sem), SC × age interactions were confined to a subset of temporo-temporal and fronto-temporal connections across right- and inter-hemispheric regions (Figure 5A, 6B). In verbal fluency, higher SC within the inter-hemispheric fronto–temporal pathway (LParsT_RMTG) predicted better performance in elderly (70-88 years), consistent with compensatory recruitment of right–hemisphere homologues during age–related production decline (Cabeza, 2002; Reuter-Lorenz & Cappell, 2008; Wierenga et al., 2008; Yeske et al., 2021). In picture–priming, enhanced inter-hemispheric temporo–temporal SC (LSTS_RMTG) facilitated response times in elderly (65-88 years).

These temporally specific effects suggest that structural compensation in production follows a delayed, threshold–dependent mechanism (Reuter-Lorenz & Cappell, 2008). This contrasts with comprehension-related compensation, which shows more gradual engagement across age (18-88 years), indicating that production and comprehension may rely on fundamentally different compensatory architectures (Baciu et al., 2021; Diaz et al., 2016; H. Zhang & Diaz, 2023a).

Significant FC × age interactions during LC tasks were limited to semantically complex sentence processing (Sem_comp), where lower fronto–temporal FC predicted better performance among elderly (60-88 years), possibly reflecting a shift from efficient compensation to de-differentiation or over-recruitment (Baciu et al., 2021; H. Zhang & Diaz, 2023a). Whereas for LP tasks, particularly VF performance followed an inverted-U shaped trajectory associated with inter-hemispheric fronto-frontal FC (LParsO_RParsO), aligning with cognitive flexibility patterns (Uddin, 2021). For picture priming task, varying semantic and phonological priming conditions were associated with distinct FC edges, highlighting the differential neural substrates underlying these processes (Hoffman & Morcom, 2018; Jarret et al., 2022; Kircher et al., 2009). For specific edges, trajectory plots showed that higher FC is associated with better performance in younger adults (18-40 years), but lower FC can become advantageous with advanced age (70-88 years), suggesting a dynamic shift in compensatory mechanisms (Cabeza et al., 2002; Reuter-Lorenz & Cappell, 2008).

In summary, “frontal vulnerability” and “temporal resilience” are identified as key neuroanatomical signatures, supporting anterior-to-posterior and last-in-first-out models of white matter aging (Bastin et al., 2010; Madden et al., 2020; Salat et al., 2005; Sullivan et al., 2010) and CRUNCH model of neurocompesnsation (Reuter-Lorenz & Cappell, 2008) for language aging. An important limitation exists that focusing on cLAN inherently excludes broader language and domain-general networks that contribute to language function and aging. Future studies expanding beyond the cLAN and integrating executive function measures are needed to fully capture the complexity of language aging.

### Data and Code availability

The neuroimaging data of cognitively healthy participants were obtained from CamCAN repository (available at https://www.mrc-cbu.cam.ac.uk/datasets/camcan/; Shafto et al., 2014; Taylor et al., 2017). Custom shell script developed for DWI preprocessing and main analysis scripts can be downloaded from GitHub (https://github.com/PartikaJain/Language_aging)

## Acknowledgements

Authors acknowledge the generous support of Computing and Network Facility of NBRC.

