## Supplementary Info for "Frontal Vulnerability vs. Temporal Resilience within the core Language Network: Neuro-compensatory mechanisms underline differential language aging trajectories"

**Supplementary information for paper title:** Frontal Vulnerability vs. Temporal Resilience within the core Language Network: Neuro-compensatory mechanisms underline differential language aging trajectories.

**Abbreviated title**: Frontal-Temporal Reorganization in Language Aging

**Authors:** Partika Jain[1], Azman Akhter[1], Arpan Banerjee[1]

**Affiliation**: [1] National Brain Research Centre, NH 8, Manesar, Gurgaon 122052, India

Here, supplementary tables 1 and 2 represents the results of linear mixed-effects models assessing age-related changes across 16 language behavior tasks. Table S1 models gender as a random effect, whereas Table S2 models gender as a fixed effect, with education treated as a random effect in both. For generating Table S2, equations 1 and 2 are rewritten as

$$LB_{\{i\}} \sim 1 + gender + (1|Education) \qquad (Null\ model) \qquad S1$$

$$LB_{\{i\}} \sim 1 + age + gender + (1|Education) \qquad (Full\ model) \qquad S2$$

The model estimates for age and the overall model fit indices (AIC, log-likelihood, and FDR-adjusted p-values) show negligible differences between the two modeling strategies (Eqn 1-2, S1-S2) across all language tasks. This empirical finding attests to the stability and robustness of the model results, regardless of how gender is parameterized.

And the supplementary tables 3 and 4 provide a comprehensive depiction of the results derived from the generalized additive mixed models (GAMMs) that systematically explore the non-linear relationships between age-related brain connectivity metrics and language behavior across the adult lifespan. These tables collectively facilitate the identification of domain- and edge-specific connectivity patterns that significantly modulate language performance across the lifespan, offering an explicit quantitative basis for the manuscript's inferences regarding selective neuro-compensatory mechanisms in healthy aging.

**Supplementary table 1**: Linear Mixed Models (LMMs) results representing the age-related variation in each of the 16 language behaviour (LB) tasks, when age was treated as fixed effect, and gender and education as random effects. Each row represents one particular LB task and columns reports the model estimates: beta coefficient for age; and performance matrices: AIC, log-likelihood and FDR corrected p-values.

| Language task | Estimate age | AIC value | Log-likelihood | FDR p-value |
|---|---|---|---|---|
| **VF** | -1.23E-02 | 1.49E+03 | -7.38E+02 | 1.33E-07 |
| **High_phon** | -3.26E-03 | 1.52E+03 | -7.55E+02 | 2.66E-01 |
| **High_sem** | -7.87E-03 | 1.51E+03 | -7.50E+02 | 2.87E-03 |
| **Low_phon** | 3.14E-03 | 1.52E+03 | -7.55E+02 | 2.66E-01 |
| **Low_sem** | -5.26E-04 | 1.52E+03 | -7.57E+02 | 8.26E-01 |
| **RT_high_phon** | -6.42E-03 | 1.36E+03 | -6.76E+02 | 2.01E-02 |
| **RT_high_sem** | -5.90E-03 | 1.36E+03 | -6.76E+02 | 3.20E-02 |
| **RT_low_phon** | -9.23E-03 | 1.35E+03 | -6.72E+02 | 6.44E-04 |
| **RT_low_sem** | -7.05E-03 | 1.36E+03 | -6.75E+02 | 1.07E-02 |
| **ToT** | -1.68E-02 | 1.46E+03 | -7.24E+02 | 3.02E-11 |
| **Vocab** | 1.69E-02 | 1.49E+03 | -7.40E+02 | 7.50E-13 |
| **Proverb** | 1.22E-02 | 1.47E+03 | -7.30E+02 | 7.64E-07 |
| **Syn_comp** | 1.77E-03 | 1.44E+03 | -7.14E+02 | 5.80E-01 |
| **Sem_comp** | -6.50E-04 | 1.44E+03 | -7.13E+02 | 8.26E-01 |
| **RT_Syn_comp** | -3.56E-03 | 1.11E+03 | -5.51E+02 | 2.66E-01 |
| **RT_Sem_comp** | -1.01E-03 | 1.13E+03 | -5.60E+02 | 8.10E-01 |

**Supplementary table 2**: Linear Mixed Models (LMMs) results representing the age-related variation in each of the 16 language behaviour (LB) tasks, when age and gender were treated as fixed effects, and education as random effect. Each row represents one particular LB task and columns reports the model estimates: beta coefficient for age; and performance matrices: AIC, log-likelihood and FDR corrected p-values.

| **Language task** | **Estimate age** | **AIC value** | **Log-likelihood** | **FDR p-value** |
|---|---|---|---|---|
| **VF** | -1.22E-02 | 1.48E+03 | -7.37E+02 | 1.50E-07 |
| **High_phon** | -3.20E-03 | 1.52E+03 | -7.53E+02 | 2.60E-01 |
| **High_sem** | -7.99E-03 | 1.51E+03 | -7.48E+02 | 1.87E-03 |
| **Low_phon** | 3.24E-03 | 1.52E+03 | -7.54E+02 | 2.60E-01 |
| **Low_sem** | -4.49E-04 | 1.52E+03 | -7.57E+02 | 8.51E-01 |
| **RT_high_phon** | -6.43E-03 | 1.36E+03 | -6.76E+02 | 1.98E-02 |
| **RT_high_sem** | -5.97E-03 | 1.36E+03 | -6.76E+02 | 2.95E-02 |
| **RT_low_phon** | -9.17E-03 | 1.35E+03 | -6.71E+02 | 6.99E-04 |
| **RT_low_sem** | -7.03E-03 | 1.36E+03 | -6.75E+02 | 1.09E-02 |
| **ToT** | -1.68E-02 | 1.46E+03 | -7.24E+02 | 2.93E-11 |
| **Vocab** | 1.68E-02 | 1.49E+03 | -7.38E+02 | 8.21E-13 |
| **Proverb** | 1.21E-02 | 1.47E+03 | -7.28E+02 | 8.86E-07 |
| **Syn_comp** | 1.77E-03 | 1.44E+03 | -7.14E+02 | 5.79E-01 |
| **Sem_comp** | -5.84E-04 | 1.44E+03 | -7.13E+02 | 8.51E-01 |
| **RT_Syn_comp** | -3.59E-03 | 1.11E+03 | -5.50E+02 | 2.60E-01 |
| **RT_Sem_comp** | -1.11E-03 | 1.13E+03 | -5.60E+02 | 7.80E-01 |

**Supplementary Table 3**. Generalized Additive Mixed Model (GAMM) results for SC × age interactions across 45 language network edges and 16 language behaviours. Each row represents a specific language behavior for a given pair of SC edges. Reported metrics include the estimated degrees of freedom (EDF) for the interaction term, adjusted $R^2$ of the full model (Adj. $R^2$), F statistics value (F value), false discovery rate-corrected p-value (FDR P-value), and Akaike Information Criterion (AIC). FDR correction was applied across all SC edges within each language behavior. Higher edf values indicate more complex, nonlinear relationships, while significant FDR-corrected p-values denote reliable age-by-SC interactions contributing to individual differences in language performance.

| SC Edge | Language Behaviour (LB) | Edf | Adj. R2 | F value | FDR P-value | AIC | SC Edge | Edf | Adj. R2 | F value | FDR P-value | AIC |
|---|---|---|---|---|---|---|---|---|---|---|---|---|
| LSTS_LMTG | VF | 1 | 0.23 | 1 | 0.59 | 1460.33 | LSTS_LParsO | 1 | 0.23 | 1 | 0.32 | 1459.57 |
| | Vocab | 3.37 | 0.24 | 1 | 0.04 | 1462.31 | | 1.46 | 0.22 | 1 | 0.35 | 1470.46 |
| | High_phon | 1.86 | 0.05 | 1 | 0.05 | 1502.04 | | 1 | 0.01 | 0.81 | 0.67 | 1517.61 |
| | High_sem | 4.66 | 0.07 | 1 | 0.09 | 1499.66 | | 2.27 | 0.05 | 1 | 0.08 | 1507.37 |
| | Low_phon | 2.28 | 0.07 | 1 | 0.03 | 1497.29 | | 1.14 | 0.04 | 1 | 0.35 | 1512.04 |
| | Low_sem | 2.02 | 0.04 | 1 | 0.01 | 1513.86 | | 2.65 | 0.02 | 1 | 0.09 | 1522.03 |
| | RT_high_phon | 1 | 0.02 | 1 | 0.62 | 1358.1 | | 1 | 0.01 | 0.45 | 0.84 | 1361.92 |
| | RT_high_sem | 2.64 | 0.02 | 1 | 0.57 | 1358.17 | | 1 | 0.01 | 1 | 0.61 | 1362.73 |
| | RT_low_phon | 1.16 | 0.03 | 0.84 | 0.73 | 1353.55 | | 1 | 0.03 | 1 | 0.15 | 1353.21 |
| | RT_low_sem | 1.09 | 0.02 | 1 | 0.25 | 1357.05 | | 1.84 | 0.02 | 1 | 0.49 | 1358.87 |
| | Proverb | 1 | 0.17 | 0.85 | 0.55 | 1458.37 | | 1.42 | 0.17 | 0.66 | 0.84 | 1461.05 |
| | Syn_comp | 1 | 0 | 0.67 | 0.87 | 1436.91 | | 1 | 0 | 1 | 0.26 | 1437.29 |
| | Sem_comp | 1 | 0 | 0.34 | 0.7 | 1435.97 | | 1 | 0 | 0.55 | 0.58 | 1436.05 |
| | RT_Syn_comp | 4.17 | 0.02 | 1 | 0.33 | 1108.13 | | 1.12 | 0.01 | 0.39 | 0.85 | 1110.74 |
| | RT_Sem_comp | 1 | 0 | 1 | 0.32 | 1128.81 | | 3.33 | 0.03 | 1 | 0.31 | 1120.06 |
| | ToT | 1.18 | 0.15 | 0.18 | 0.95 | 1443.8 | | 1 | 0.14 | 0.13 | 0.88 | 1448.81 |
| LSTS_LMTG | VF | 1.56 | 0.23 | 1 | 0.49 | 1459.34 | LSTS_LSTG | 1 | 0.23 | 1 | 0.66 | 1461.13 |
| | Vocab | 1 | 0.21 | 1 | 0.25 | 1472.99 | | 1 | 0.21 | 1 | 0.58 | 1474.19 |
| | High_phon | 1 | 0.01 | 0 | 0.99 | 1517.72 | | 2.3 | 0.03 | 1 | 0.03 | 1509.82 |
| | High_sem | 1 | 0.04 | 0.11 | 0.93 | 1510.41 | | 2.21 | 0.04 | 1 | 0.13 | 1506.21 |
| | Low_phon | 1 | 0.05 | 1 | 0.01 | 1503.5 | | 3.32 | 0.06 | 1 | 0.62 | 1511.57 |
| | Low_sem | 2.87 | 0.02 | 1 | 0.03 | 1519.2 | | 1.69 | 0.01 | 1 | 0.53 | 1522.49 |
| | RT_high_phon | 1.75 | 0.02 | 1 | 0.27 | 1361.54 | | 2.1 | 0.02 | 1 | 0.2 | 1358.76 |
| | RT_high_sem | 1 | 0.01 | 1 | 0.51 | 1362.4 | | 2.18 | 0.02 | 1 | 0.17 | 1356.9 |
| | RT_low_phon | 1.37 | 0.03 | 1 | 0.66 | 1355.04 | | 1 | 0.06 | 1 | 0.41 | 1342.47 |
| | RT_low_sem | 1 | 0.02 | 1 | 0.14 | 1357.14 | | 1 | 0.01 | 1 | 0.58 | 1358.94 |
| | Proverb | 1 | 0.16 | 0.66 | 0.66 | 1459.05 | | 1 | 0.16 | 0.7 | 0.59 | 1456.97 |
| | Syn_comp | 2.34 | 0 | 1 | 0.72 | 1438.21 | | 2.46 | 0.01 | 1 | 0.38 | 1438.53 |
| | Sem_comp | 1 | 0 | 0.14 | 0.87 | 1433.67 | | 1 | 0 | 1 | 0.14 | 1434.82 |
| | RT_Syn_comp | 2.41 | 0.01 | 1 | 0.43 | 1110.85 | | 2.01 | 0.01 | 1 | 0.43 | 1109.76 |
| | RT_Sem_comp | 1.57 | 0 | 1 | 0.93 | 1128.67 | | 2.83 | 0.02 | 1 | 0.12 | 1124.91 |

| | | | | | | | | | | | | |
|---|---|---|---|---|---|---|---|---|---|---|---|---|
| | ToT | 1 | 0.14 | 0.18 | 0.83 | 1445.07 | | 1 | 0.15 | 1 | 0.06 | 1438.34 |
| **LMTG_LParsO** | VF | 1 | 0.23 | 1 | 0.29 | 1459.42 | **LMTG_LParsT** | 2.06 | 0.24 | 1 | 0.24 | 1459.19 |
| | Vocab | 1.63 | 0.22 | 1 | 0.26 | 1471.59 | | 1 | 0.21 | 1 | 0.63 | 1474.06 |
| | High_phon | 1 | 0.02 | 1 | 0.44 | 1516.27 | | 1 | 0.01 | 1 | 0.23 | 1516.47 |
| | High_sem | 1 | 0.04 | 1 | 0.13 | 1509.17 | | 1 | 0.04 | 1 | 0.17 | 1509.61 |
| | Low_phon | 1 | 0.04 | 1 | 0.36 | 1512.97 | | 1.19 | 0.04 | 1 | 0.44 | 1513.44 |
| | Low_sem | 1 | 0 | 0.15 | 0.91 | 1525.49 | | 1 | 0.01 | 1 | 0.24 | 1522.83 |
| | RT_high_phon | 1 | 0.01 | 1 | 0.33 | 1360.77 | | 1 | 0.01 | 0.45 | 0.87 | 1361.99 |
| | RT_high_sem | 1 | 0.01 | 1 | 0.45 | 1362.08 | | 1 | 0.01 | 1 | 0.42 | 1362.3 |
| | RT_low_phon | 1 | 0.03 | 1 | 0.41 | 1353.11 | | 3.04 | 0.04 | 1 | 0.23 | 1352.48 |
| | RT_low_sem | 1 | 0.01 | 0.82 | 0.76 | 1359.45 | | 1 | 0.01 | 0 | 0.95 | 1360.24 |
| | Proverb | 1.54 | 0.16 | 0.92 | 0.64 | 1461.25 | | 1.6 | 0.16 | 1 | 0.36 | 1457.55 |
| | Syn_comp | 1.79 | 0.01 | 1 | 0.08 | 1434.65 | | 3.99 | 0.01 | 1 | 0.21 | 1435.55 |
| | Sem_comp | 1 | 0 | 0.07 | 0.93 | 1435.86 | | 1 | 0 | 0.07 | 0.92 | 1434.01 |
| | RT_Syn_comp | 1 | 0.01 | 0.22 | 0.86 | 1110.5 | | 2.09 | 0.01 | 1 | 0.22 | 1109.35 |
| | RT_Sem_comp | 1 | 0.02 | 1 | 0.55 | 1119.14 | | 1.68 | 0.02 | 1 | 0.62 | 1123.06 |
| | ToT | 1.84 | 0.14 | 1 | 0.39 | 1445.58 | | 1 | 0.14 | 0.13 | 0.91 | 1445.66 |
| **LMTG_LSTG** | VF | 2.32 | 0.23 | 1 | 0.69 | 1462.25 | **LParsO_LParsT** | 1.21 | 0.23 | 1 | 0.5 | 1463.24 |
| | Vocab | 1 | 0.21 | 1 | 0.81 | 1473.97 | | 1.97 | 0.21 | 1 | 0.28 | 1473.63 |
| | High_phon | 1 | 0.02 | 1 | 0.07 | 1514.09 | | 1.93 | 0.01 | 0.93 | 0.67 | 1517.97 |
| | High_sem | 1 | 0.04 | 1 | 0.14 | 1510.21 | | 2.32 | 0.04 | 1 | 0.26 | 1508.68 |
| | Low_phon | 1 | 0.04 | 0.51 | 0.79 | 1514.61 | | 3.15 | 0.04 | 1 | 0.34 | 1512.59 |
| | Low_sem | 1 | 0.01 | 0.34 | 0.86 | 1524.47 | | 2.97 | 0.02 | 1 | 0.06 | 1520.15 |
| | RT_high_phon | 1 | 0.01 | 1 | 0.59 | 1359.42 | | 2.03 | 0.02 | 1 | 0.15 | 1359.11 |
| | RT_high_sem | 2.16 | 0.02 | 1 | 0.28 | 1359.51 | | 1.04 | 0.01 | 1 | 0.21 | 1361.34 |
| | RT_low_phon | 1 | 0.03 | 1 | 0.62 | 1353.84 | | 1 | 0.03 | 0.75 | 0.86 | 1355.52 |
| | RT_low_sem | 1.93 | 0.03 | 1 | 0.02 | 1352.83 | | 1.51 | 0.02 | 1 | 0.45 | 1359 |
| | Proverb | 1 | 0.17 | 0.66 | 0.68 | 1455.46 | | 1 | 0.16 | 1 | 0.32 | 1455.85 |
| | Syn_comp | 2.47 | 0 | 1 | 0.79 | 1437.85 | | 1 | 0 | 0 | 0.95 | 1438.64 |
| | Sem_comp | 3.1 | 0.01 | 0.83 | 0.62 | 1435.33 | | 1.89 | 0 | 0.71 | 0.63 | 1436.77 |
| | RT_Syn_comp | 3.15 | 0.02 | 1 | 0.53 | 1110.54 | | 1.1 | 0.01 | 0.67 | 0.84 | 1108.39 |
| | RT_Sem_comp | 1.81 | 0 | 1 | 0.43 | 1129.41 | | 1 | -0.01 | 1 | 0.56 | 1129.24 |
| | ToT | 1 | 0.14 | 0.13 | 0.94 | 1444.64 | | 1 | 0.14 | 1 | 0.39 | 1446.31 |
| **LParsO_LSTG** | VF | 2.88 | 0.24 | 1 | 0.47 | 1461.56 | **LParsT_LSTG** | 3.34 | 0.25 | 1 | 0.04 | 1453.76 |
| | Vocab | 2.4 | 0.22 | 1 | 0.11 | 1469.04 | | 2.01 | 0.22 | 1 | 0.19 | 1471.89 |
| | High_phon | 3.81 | 0.04 | 1 | 0.06 | 1510.33 | | 1 | 0.01 | 0.52 | 0.79 | 1517.74 |
| | High_sem | 2.24 | 0.04 | 1 | 0.63 | 1512.61 | | 1.74 | 0.04 | 1 | 0.34 | 1510.86 |
| | Low_phon | 4.13 | 0.06 | 1 | 0.05 | 1505.71 | | 3.81 | 0.06 | 1 | 0.02 | 1504.1 |
| | Low_sem | 3.59 | 0.01 | 1 | 0.34 | 1524.15 | | 2.02 | 0.01 | 1 | 0.25 | 1523.37 |
| | RT_high_phon | 1 | 0.01 | 1 | 0.7 | 1361.39 | | 1.59 | 0.01 | 1 | 0.91 | 1362.71 |
| | RT_high_sem | 1 | 0.02 | 1 | 0.16 | 1355.06 | | 1.9 | 0.02 | 1 | 0.41 | 1360.69 |
| | RT_low_phon | 1 | 0.03 | 1 | 0.42 | 1354.17 | | 1 | 0.03 | 0.75 | 0.82 | 1354.86 |
| | RT_low_sem | 1.25 | 0.02 | 0.82 | 0.91 | 1360.23 | | 2 | 0.02 | 1 | 0.17 | 1357.65 |
| | Proverb | 1 | 0.16 | 0.9 | 0.52 | 1455.88 | | 1.66 | 0.16 | 1 | 0.24 | 1459.63 |
| | Syn_comp | 1.96 | 0 | 1 | 0.73 | 1439.68 | | 6.62 | 0.07 | 1 | 0 | 1411.59 |
| | Sem_comp | 4.63 | 0.02 | 1 | 0.09 | 1428.33 | | 1 | 0 | 0.19 | 0.81 | 1434.87 |
| | RT_Syn_comp | 1 | 0.01 | 0.73 | 0.62 | 1109.49 | | 1.66 | 0.01 | 1 | 0.46 | 1110.4 |

| | | | | | | | | | | | |
|---|---|---|---|---|---|---|---|---|---|---|---|
| | RT_Sem_comp | 1.64 | 0 | 1 | 0.5 | 1128.77 | | 1 | 0 | 1 | 0.26 | 1127.91 |
| | ToT | 1.12 | 0.14 | 0.17 | 0.96 | 1449.41 | | 2.16 | 0.14 | 1 | 0.53 | 1446.12 |
| **RSTS_RMTG** | VF | 1 | 0.23 | 1 | 0.29 | 1460.23 | **RSTS_RParsO** | 2.77 | 0.24 | 1 | 0.23 | 1459.87 |
| | Vocab | 1 | 0.21 | 1 | 0.31 | 1472.4 | | 2.04 | 0.21 | 1 | 0.42 | 1475.83 |
| | High_phon | 1.24 | 0.01 | 0.82 | 0.85 | 1516.69 | | 2.63 | 0.04 | 1 | 0.01 | 1504.5 |
| | High_sem | 2.32 | 0.05 | 1 | 0.09 | 1504.24 | | 1.54 | 0.04 | 1 | 0.59 | 1509.2 |
| | Low_phon | 4.17 | 0.05 | 1 | 0.41 | 1510.45 | | 1 | 0.04 | 1 | 0.47 | 1509.23 |
| | Low_sem | 1 | 0 | 0.39 | 0.82 | 1525.68 | | 2.13 | 0.02 | 1 | 0.22 | 1521.43 |
| | RT_high_phon | 1 | 0.02 | 1 | 0.49 | 1357.23 | | 2.05 | 0.03 | 1 | 0.03 | 1355.73 |
| | RT_high_sem | 1 | 0.01 | 0.54 | 0.81 | 1362.6 | | 1 | 0.02 | 1 | 0.03 | 1358.93 |
| | RT_low_phon | 1.47 | 0.03 | 1 | 0.66 | 1354.64 | | 2.15 | 0.05 | 1 | 0.06 | 1348.2 |
| | RT_low_sem | 1 | 0.02 | 1 | 0.13 | 1357.24 | | 1.94 | 0.03 | 1 | 0.08 | 1355.19 |
| | Proverb | 2.24 | 0.17 | 1 | 0.07 | 1449.8 | | 1 | 0.17 | 1 | 0.1 | 1453.12 |
| | Syn_comp | 1 | 0 | 1 | 0.28 | 1437.16 | | 3.44 | 0 | 1 | 0.64 | 1439.95 |
| | Sem_comp | 1 | 0 | 0.25 | 0.75 | 1436.29 | | 1 | 0 | 0.26 | 0.74 | 1434.43 |
| | RT_Syn_comp | 1.01 | 0 | 1 | 0.37 | 1110.28 | | 1 | 0.01 | 0.72 | 0.69 | 1110.78 |
| | RT_Sem_comp | 3.74 | 0.03 | 1 | 0.08 | 1123.47 | | 1.61 | 0 | 1 | 0.43 | 1129.04 |
| | ToT | 1 | 0.14 | 1 | 0.46 | 1445.02 | | 1 | 0.14 | 0.45 | 0.72 | 1443.92 |
| **RSTS_RParsT** | VF | 1 | 0.23 | 1 | 0.15 | 1458.86 | **RSTS_RSTG** | 1.83 | 0.24 | 1 | 0.29 | 1460.48 |
| | Vocab | 1.7 | 0.21 | 1 | 0.38 | 1475.4 | | 1 | 0.21 | 1 | 0.54 | 1474.39 |
| | High_phon | 3.87 | 0.03 | 1 | 0.09 | 1512.31 | | 1 | 0.01 | 1 | 0.4 | 1515.74 |
| | High_sem | 1 | 0.03 | 0.51 | 0.77 | 1511.07 | | 1 | 0.04 | 1 | 0.31 | 1511.25 |
| | Low_phon | 1 | 0.04 | 1 | 0.46 | 1512.08 | | 1 | 0.03 | 0 | 0.98 | 1514.05 |
| | Low_sem | 1.38 | 0 | 0.71 | 0.69 | 1525.8 | | 1 | 0 | 0.15 | 0.93 | 1525.72 |
| | RT_high_phon | 2.61 | 0.03 | 1 | 0.04 | 1356.16 | | 2.24 | 0.03 | 1 | 0.03 | 1354.58 |
| | RT_high_sem | 1.23 | 0.01 | 1 | 0.35 | 1361.74 | | 4.13 | 0.07 | 1 | 0 | 1341.82 |
| | RT_low_phon | 1 | 0.03 | 1 | 0.32 | 1354.04 | | 1.27 | 0.03 | 1 | 0.14 | 1352.35 |
| | RT_low_sem | 1 | 0.02 | 1 | 0.5 | 1357.73 | | 5.64 | 0.06 | 1 | 0 | 1342.13 |
| | Proverb | 2.06 | 0.17 | 1 | 0.19 | 1457.1 | | 1 | 0.17 | 1 | 0.41 | 1452.21 |
| | Syn_comp | 3.17 | 0.01 | 1 | 0.1 | 1433.07 | | 1 | 0 | 1 | 0.63 | 1438.3 |
| | Sem_comp | 1 | 0 | 0.13 | 0.87 | 1434.69 | | 2.89 | 0 | 0.83 | 0.64 | 1437.78 |
| | RT_Syn_comp | 2.09 | 0.02 | 1 | 0.07 | 1106.06 | | 1 | 0.01 | 0.72 | 0.69 | 1110.46 |
| | RT_Sem_comp | 1 | 0.01 | 1 | 0.16 | 1125.48 | | 2.64 | 0.01 | 1 | 0.47 | 1128.19 |
| | ToT | 1 | 0.14 | 0.13 | 0.89 | 1444.36 | | 1 | 0.15 | 0.13 | 0.93 | 1441.7 |
| **RMTG_RParsO** | VF | 1.26 | 0.23 | 1 | 0.56 | 1461.42 | **RMTG_RParsT** | 1 | 0.23 | 1 | 0.1 | 1458.69 |
| | Vocab | 2.2 | 0.22 | 1 | 0.14 | 1470.57 | | 1.67 | 0.21 | 1 | 0.44 | 1476.78 |
| | High_phon | 4.27 | 0.04 | 1 | 0.14 | 1511.54 | | 2.49 | 0.02 | 1 | 0.54 | 1515.38 |
| | High_sem | 1 | 0.03 | 0.27 | 0.85 | 1510.03 | | 1 | 0.04 | 0.38 | 0.82 | 1511.4 |
| | Low_phon | 3.54 | 0.06 | 1 | 0.08 | 1507.37 | | 1 | 0.04 | 1 | 0.21 | 1510.38 |
| | Low_sem | 2.09 | 0.02 | 1 | 0.44 | 1521.35 | | 1 | 0 | 1 | 0.27 | 1525.09 |
| | RT_high_phon | 2.86 | 0.03 | 1 | 0.11 | 1357.53 | | 1.54 | 0.02 | 1 | 0.32 | 1360.84 |
| | RT_high_sem | 2.79 | 0.02 | 1 | 0.1 | 1359.08 | | 1 | 0.01 | 1 | 0.25 | 1361.45 |
| | RT_low_phon | 3.2 | 0.04 | 1 | 0.22 | 1353.06 | | 1 | 0.03 | 1 | 0.24 | 1352.77 |
| | RT_low_sem | 4.14 | 0.04 | 1 | 0.04 | 1354.51 | | 1 | 0.01 | 1 | 0.39 | 1359.2 |
| | Proverb | 1 | 0.17 | 1 | 0.1 | 1453.96 | | 1.57 | 0.19 | 0.88 | 0.7 | 1446.26 |
| | Syn_comp | 1 | 0 | 1 | 0.7 | 1438.74 | | 1.77 | 0 | 1 | 0.42 | 1436.51 |
| | Sem_comp | 1 | 0 | 0.37 | 0.67 | 1434.33 | | 1 | 0 | 0 | 0.96 | 1435.38 |

| | | | | | | | | | | | | |
|---|---|---|---|---|---|---|---|---|---|---|---|---|
| | RT_Syn_comp | 1.73 | 0.01 | 0.73 | 0.88 | 1111.5 | | 2.09 | 0.02 | 1 | 0.15 | 1107.16 |
| | RT_Sem_comp | 1 | 0 | 1 | 0.17 | 1127.48 | | 1 | 0.01 | 1 | 0.23 | 1124.31 |
| | ToT | 1 | 0.14 | 1 | 0.24 | 1445.22 | | 1.73 | 0.14 | 1 | 0.5 | 1444.96 |
| **RMTG_RSTG** | VF | 1 | 0.23 | 1 | 0.57 | 1460.04 | **RParsO_RParsT** | 1 | 0.23 | 1 | 0.79 | 1462.15 |
| | Vocab | 2.78 | 0.22 | 1 | 0.29 | 1472.09 | | 2.49 | 0.22 | 1 | 0.03 | 1468.63 |
| | High_phon | 1 | 0.01 | 0.52 | 0.8 | 1516.86 | | 1.33 | 0.02 | 1 | 0.52 | 1514.96 |
| | High_sem | 1 | 0.04 | 0 | 0.95 | 1509.54 | | 1 | 0.03 | 0.75 | 0.7 | 1506.19 |
| | Low_phon | 1 | 0.04 | 1 | 0.46 | 1512.86 | | 1 | 0.04 | 1 | 0.42 | 1512.82 |
| | Low_sem | 1 | 0 | 0.39 | 0.8 | 1525.65 | | 1 | 0 | 0.15 | 0.94 | 1523.94 |
| | RT_high_phon | 1 | 0.01 | 0.22 | 0.94 | 1360.8 | | 1 | 0.01 | 1 | 0.63 | 1361.56 |
| | RT_high_sem | 1 | 0.02 | 1 | 0.03 | 1358.14 | | 1 | 0.01 | 1 | 0.26 | 1361.66 |
| | RT_low_phon | 1.89 | 0.03 | 1 | 0.7 | 1355.15 | | 4.19 | 0.04 | 1 | 0.28 | 1353.67 |
| | RT_low_sem | 3.67 | 0.03 | 1 | 0.41 | 1358.44 | | 1 | 0.01 | 1 | 0.61 | 1360.23 |
| | Proverb | 1 | 0.17 | 1 | 0.43 | 1449.25 | | 1 | 0.18 | 0.7 | 0.64 | 1448.91 |
| | Syn_comp | 1 | 0 | 1 | 0.08 | 1435.66 | | 5.31 | 0.03 | 1 | 0.06 | 1429.94 |
| | Sem_comp | 1 | 0 | 0.55 | 0.56 | 1436.35 | | 2.42 | 0 | 1 | 0.45 | 1435.89 |
| | RT_Syn_comp | 1 | 0.01 | 0.73 | 0.63 | 1110.34 | | 1.53 | 0.01 | 1 | 0.63 | 1109.91 |
| | RT_Sem_comp | 2.09 | 0.01 | 1 | 0.72 | 1126.06 | | 2.43 | 0.01 | 1 | 0.07 | 1124.88 |
| | ToT | 1 | 0.14 | 1 | 0.52 | 1445.22 | | 1 | 0.14 | 0.13 | 0.88 | 1445.27 |
| **RMTG_RSTG** | VF | 1 | 0.23 | 1 | 0.71 | 1460.17 | **RParsT_RSTG** | 3.07 | 0.24 | 1 | 0.15 | 1458.6 |
| | Vocab | 1 | 0.21 | 1 | 0.73 | 1472.77 | | 2.46 | 0.22 | 1 | 0.09 | 1471.15 |
| | High_phon | 1 | 0.01 | 1 | 0.51 | 1516.54 | | 2.83 | 0.02 | 1 | 0.46 | 1515.03 |
| | High_sem | 1.49 | 0.04 | 0.86 | 0.84 | 1507.47 | | 1.92 | 0.04 | 1 | 0.1 | 1507 |
| | Low_phon | 1 | 0.04 | 0 | 0.96 | 1513.57 | | 1 | 0.04 | 1 | 0.24 | 1512.81 |
| | Low_sem | 1.2 | 0.01 | 1 | 0.23 | 1523.99 | | 1.98 | 0 | 1 | 0.69 | 1525.81 |
| | RT_high_phon | 1 | 0.02 | 1 | 0.08 | 1355.6 | | 1.59 | 0.03 | 1 | 0.31 | 1355.17 |
| | RT_high_sem | 1 | 0.01 | 0.67 | 0.76 | 1362.88 | | 1 | 0.01 | 1 | 0.35 | 1362.09 |
| | RT_low_phon | 2.64 | 0.04 | 1 | 0.24 | 1352.86 | | 1.83 | 0.04 | 1 | 0.07 | 1350.84 |
| | RT_low_sem | 1 | 0.01 | 1 | 0.38 | 1358.86 | | 1 | 0.01 | 1 | 0.49 | 1358.96 |
| | Proverb | 1 | 0.17 | 0 | 0.95 | 1452.18 | | 1.54 | 0.17 | 1 | 0.27 | 1450.92 |
| | Syn_comp | 2.84 | 0.01 | 1 | 0.59 | 1435.5 | | 1 | 0 | 1 | 0.29 | 1437.66 |
| | Sem_comp | 1.19 | 0 | 0.49 | 0.65 | 1435.89 | | 1 | 0 | 0.24 | 0.78 | 1436.16 |
| | RT_Syn_comp | 1.03 | 0.01 | 0.15 | 0.99 | 1110.71 | | 1.93 | 0.01 | 0.73 | 0.67 | 1110.84 |
| | RT_Sem_comp | 1 | 0 | 1 | 0.51 | 1127.41 | | 1 | 0 | 1 | 0.66 | 1127.93 |
| | ToT | 1 | 0.14 | 0.8 | 0.63 | 1444.23 | | 1 | 0.14 | 0.13 | 0.9 | 1444.66 |
| **LSTS_RSTS** | VF | 1.24 | 0.23 | 1 | 0.89 | 1462.07 | **LSTS_RMTG** | 1 | 0.23 | 1 | 0.37 | 1459.54 |
| | Vocab | 1 | 0.21 | 1 | 0.67 | 1473.54 | | 1 | 0.22 | 1 | 0.07 | 1470.1 |
| | High_phon | 1 | 0.01 | 1 | 0.53 | 1516.76 | | 1 | 0.01 | 1 | 0.36 | 1517.06 |
| | High_sem | 1.7 | 0.04 | 1 | 0.32 | 1509.99 | | 1 | 0.04 | 0 | 0.96 | 1511.78 |
| | Low_phon | 1.26 | 0.04 | 0.51 | 0.84 | 1514.22 | | 1 | 0.04 | 1 | 0.58 | 1513.82 |
| | Low_sem | 1 | 0 | 0.39 | 0.8 | 1525.2 | | 1 | 0 | 0.63 | 0.71 | 1525.52 |
| | RT_high_phon | 3.11 | 0.03 | 1 | 0.19 | 1356.64 | | 6.4 | 0.04 | 1 | 0.07 | 1353.52 |
| | RT_high_sem | 2.58 | 0.06 | 1 | 0 | 1342.02 | | 4.54 | 0.06 | 1 | 0 | 1344.62 |
| | RT_low_phon | 1.99 | 0.05 | 1 | 0.01 | 1345 | | 2.86 | 0.04 | 1 | 0.17 | 1352.06 |
| | RT_low_sem | 2.04 | 0.03 | 1 | 0.01 | 1350.16 | | 1.65 | 0.02 | 1 | 0.33 | 1358.5 |
| | Proverb | 1 | 0.16 | 0 | 1 | 1459.44 | | 1 | 0.16 | 1 | 0.39 | 1453.46 |
| | Syn_comp | 3.87 | 0.04 | 1 | 0.02 | 1422.58 | | 3.07 | 0.13 | 1 | 0.28 | 1377.13 |

| | | | | | | | | | | | | |
|---|---|---|---|---|---|---|---|---|---|---|---|---|
| | Sem_comp | 1.04 | 0 | 0.55 | 0.61 | 1435.71 | | 2.88 | 0.01 | 1 | 0.39 | 1433.14 |
| | RT_Syn_comp | 1 | 0.01 | 1 | 0.41 | 1109.68 | | 2.21 | 0.02 | 1 | 0.36 | 1108.06 |
| | RT_Sem_comp | 1 | 0 | 1 | 0.35 | 1128.09 | | 1.48 | 0 | 1 | 0.78 | 1130.64 |
| | ToT | 2.6 | 0.16 | 1 | 0.01 | 1436.38 | | 3.48 | 0.15 | 1 | 0.07 | 1440.36 |
| **LSTS_RParsO** | VF | 1 | 0.23 | 1 | 0.43 | 1460.55 | **LSTS_RParsT** | 2.24 | 0.24 | 1 | 0.02 | 1454.62 |
| | Vocab | 1.9 | 0.21 | 1 | 0.37 | 1473.06 | | 1 | 0.21 | 1 | 0.19 | 1471.36 |
| | High_phon | 1.52 | 0.01 | 1 | 0.64 | 1517.75 | | 1.62 | 0.02 | 1 | 0.09 | 1513.76 |
| | High_sem | 1.53 | 0.04 | 1 | 0.61 | 1511.36 | | 1.79 | 0.04 | 1 | 0.15 | 1508.96 |
| | Low_phon | 1 | 0.04 | 0.9 | 0.62 | 1513.06 | | 1 | 0.04 | 1 | 0.13 | 1510.86 |
| | Low_sem | 1.77 | 0.01 | 1 | 0.45 | 1525.49 | | 1.59 | 0 | 1 | 0.49 | 1525.42 |
| | RT_high_phon | 1 | 0.02 | 1 | 0.22 | 1355.26 | | 3.26 | 0.03 | 1 | 0.17 | 1358.77 |
| | RT_high_sem | 1.5 | 0.01 | 1 | 0.31 | 1361.28 | | 3.65 | 0.02 | 1 | 0.23 | 1360.6 |
| | RT_low_phon | 1.3 | 0.03 | 0.81 | 0.86 | 1355.09 | | 2.8 | 0.03 | 1 | 0.49 | 1355.19 |
| | RT_low_sem | 2.01 | 0.02 | 1 | 0.26 | 1358.69 | | 2.61 | 0.02 | 1 | 0.5 | 1360.2 |
| | Proverb | 1 | 0.17 | 0.45 | 0.79 | 1452 | | 1 | 0.16 | 0.11 | 0.93 | 1459.02 |
| | Syn_comp | 3 | 0.01 | 1 | 0.18 | 1435.48 | | 1 | 0.24 | 1 | 0.01 | 1306.55 |
| | Sem_comp | 1 | -0.01 | 0.25 | 0.75 | 1436.49 | | 1 | 0 | 0.72 | 0.49 | 1433.96 |
| | RT_Syn_comp | 1 | 0.01 | 1 | 0.5 | 1109.22 | | 1 | 0.01 | 1 | 0.46 | 1109.8 |
| | RT_Sem_comp | 1.32 | 0 | 1 | 0.1 | 1126.19 | | 2.44 | 0.01 | 1 | 0.26 | 1127.14 |
| | ToT | 2.04 | 0.15 | 1 | 0.63 | 1443.64 | | 1.56 | 0.14 | 1 | 0.54 | 1447.31 |
| **LSTS_RSTG** | VF | 1.25 | 0.23 | 1 | 0.86 | 1460.88 | **LMTG_RSTS** | 1.64 | 0.23 | 1 | 0.45 | 1460.54 |
| | Vocab | 1 | 0.21 | 1 | 0.46 | 1473.04 | | 2.65 | 0.23 | 1 | 0 | 1459.6 |
| | High_phon | 1 | 0.01 | 0.75 | 0.7 | 1517.65 | | 1.56 | 0.01 | 1 | 0.36 | 1516.52 |
| | High_sem | 1 | 0.04 | 0.51 | 0.76 | 1510.51 | | 1.7 | 0.04 | 1 | 0.37 | 1511.21 |
| | Low_phon | 1 | 0.04 | 0.65 | 0.71 | 1513.6 | | 2 | 0.04 | 1 | 0.07 | 1506.84 |
| | Low_sem | 1 | 0 | 0.93 | 0.55 | 1525.44 | | 1.11 | 0.01 | 0.9 | 0.71 | 1523.52 |
| | RT_high_phon | 3.14 | 0.02 | 1 | 0.18 | 1358.89 | | 2.08 | 0.02 | 1 | 0.68 | 1361.72 |
| | RT_high_sem | 4.15 | 0.04 | 1 | 0.02 | 1352.62 | | 5.31 | 0.05 | 1 | 0.01 | 1351.08 |
| | RT_low_phon | 2.38 | 0.04 | 1 | 0.29 | 1353.37 | | 1.51 | 0.04 | 1 | 0.08 | 1350.26 |
| | RT_low_sem | 1.11 | 0.02 | 1 | 0.41 | 1356.89 | | 1 | 0.02 | 1 | 0.16 | 1359.01 |
| | Proverb | 1 | 0.16 | 0.7 | 0.62 | 1453.97 | | 3.07 | 0.17 | 1 | 0.3 | 1455.23 |
| | Syn_comp | 3.5 | 0.07 | 1 | 0.17 | 1410.21 | | 3.43 | 0.04 | 1 | 0.01 | 1420.88 |
| | Sem_comp | 2.72 | 0.01 | 1 | 0.35 | 1435.43 | | 2.08 | 0 | 1 | 0.53 | 1435.84 |
| | RT_Syn_comp | 1.91 | 0.02 | 1 | 0.29 | 1108.33 | | 1 | 0.01 | 0.73 | 0.66 | 1108.79 |
| | RT_Sem_comp | 3.04 | 0.02 | 1 | 0.13 | 1126.38 | | 1 | 0.01 | 1 | 0.42 | 1124.68 |
| | ToT | 3.81 | 0.15 | 1 | 0.07 | 1440.17 | | 4.41 | 0.16 | 1 | 0.01 | 1436.03 |
| **LMTG_RMTG** | VF | 1 | 0.23 | 1 | 0.14 | 1459.33 | **LMTG_RParsO** | 1.21 | 0.23 | 1 | 0.51 | 1464.99 |
| | Vocab | 1.68 | 0.23 | 1 | 0.16 | 1465.73 | | 1.08 | 0.21 | 1 | 0.34 | 1477.47 |
| | High_phon | 1.6 | 0.01 | 1 | 0.55 | 1517.46 | | 1 | 0.01 | 1 | 0.5 | 1517.22 |
| | High_sem | 1.87 | 0.04 | 1 | 0.23 | 1509.98 | | 1 | 0.04 | 1 | 0.28 | 1511.44 |
| | Low_phon | 1.85 | 0.04 | 1 | 0.16 | 1511.64 | | 1.87 | 0.05 | 1 | 0.34 | 1507.53 |
| | Low_sem | 1 | 0 | 0.63 | 0.71 | 1525.14 | | 1 | 0 | 1 | 0.18 | 1524.78 |
| | RT_high_phon | 2.38 | 0.02 | 1 | 0.14 | 1358.12 | | 1 | 0.02 | 1 | 0.14 | 1358.76 |
| | RT_high_sem | 1 | 0.01 | 1 | 0.14 | 1360.71 | | 1 | 0.01 | 0.54 | 0.82 | 1362.84 |
| | RT_low_phon | 2.82 | 0.04 | 1 | 0.33 | 1353.77 | | 1 | 0.03 | 0.75 | 0.85 | 1354.24 |
| | RT_low_sem | 1 | 0.02 | 1 | 0.11 | 1357.4 | | 1.17 | 0.01 | 0.82 | 0.8 | 1360.37 |
| | Proverb | 2.23 | 0.16 | 1 | 0.55 | 1459.01 | | 1 | 0.19 | 0.22 | 0.87 | 1442.84 |

| | | | | | | | | | | | | |
|---|---|---|---|---|---|---|---|---|---|---|---|---|
| | Syn_comp | 3.68 | 0.04 | 1 | 0 | 1419.75 | | 1 | 0 | 1 | 0.21 | 1437.07 |
| | Sem_comp | 3.35 | 0.01 | 1 | 0.43 | 1435.02 | | 1 | 0 | 0.19 | 0.81 | 1436.19 |
| | RT_Syn_comp | 1 | 0.01 | 0.15 | 0.93 | 1108.79 | | 1 | 0.01 | 1 | 0.25 | 1108.44 |
| | RT_Sem_comp | 1.92 | 0 | 1 | 0.89 | 1129.76 | | 1 | 0 | 1 | 0.33 | 1128.85 |
| | ToT | 2.85 | 0.15 | 1 | 0.38 | 1444.74 | | 1.76 | 0.14 | 1 | 0.47 | 1445.7 |
| **LMTG_RParsT** | VF | 2.95 | 0.24 | 1 | 0.07 | 1454.23 | **LMTG_RSTG** | 1 | 0.23 | 1 | 0.17 | 1458.86 |
| | Vocab | 1 | 0.21 | 1 | 0.41 | 1473.65 | | 1 | 0.21 | 1 | 0.79 | 1477.32 |
| | High_phon | 1 | 0.02 | 1 | 0.02 | 1511.78 | | 2.3 | 0.03 | 1 | 0.03 | 1510.22 |
| | High_sem | 2.36 | 0.05 | 1 | 0.07 | 1506.17 | | 3 | 0.09 | 1 | 0.02 | 1490.4 |
| | Low_phon | 1.43 | 0.04 | 1 | 0.23 | 1512.85 | | 2.23 | 0.05 | 1 | 0.04 | 1509.16 |
| | Low_sem | 1.36 | 0.01 | 1 | 0.18 | 1522.44 | | 1.79 | 0.01 | 1 | 0.51 | 1524.49 |
| | RT_high_phon | 1.26 | 0.01 | 1 | 0.53 | 1361.76 | | 3.43 | 0.03 | 1 | 0.16 | 1355.97 |
| | RT_high_sem | 1.52 | 0.01 | 1 | 0.56 | 1363.14 | | 1 | 0.01 | 1 | 0.63 | 1363.26 |
| | RT_low_phon | 2.43 | 0.03 | 1 | 0.46 | 1355.01 | | 2.33 | 0.04 | 1 | 0.46 | 1353.96 |
| | RT_low_sem | 1.97 | 0.02 | 1 | 0.63 | 1359.67 | | 1 | 0.02 | 1 | 0.3 | 1357.43 |
| | Proverb | 1.41 | 0.17 | 1 | 0.11 | 1455.02 | | 2.44 | 0.17 | 1 | 0.34 | 1463.63 |
| | Syn_comp | 7.15 | 0.28 | 1 | 0 | 1297.86 | | 2.79 | 0.04 | 1 | 0.12 | 1423.07 |
| | Sem_comp | 1 | 0 | 0.55 | 0.57 | 1436.28 | | 3.1 | 0.01 | 1 | 0.52 | 1434.63 |
| | RT_Syn_comp | 1 | 0.01 | 1 | 0.31 | 1109.28 | | 4.68 | 0.03 | 1 | 0.43 | 1108.6 |
| | RT_Sem_comp | 1.13 | 0 | 1 | 0.24 | 1126.67 | | 3.18 | 0.01 | 1 | 0.35 | 1126.65 |
| | ToT | 1 | 0.14 | 0.25 | 0.81 | 1443.49 | | 4.58 | 0.15 | 1 | 0.13 | 1440.6 |
| **LParsO_RSTS** | VF | 3.11 | 0.24 | 1 | 0.31 | 1464.18 | **LParsO_RMTG** | 1 | 0.23 | 1 | 0.62 | 1461.49 |
| | Vocab | 1 | 0.21 | 1 | 0.6 | 1473.64 | | 1 | 0.22 | 1 | 0.04 | 1471.11 |
| | High_phon | 1 | 0.02 | 1 | 0.58 | 1517.37 | | 1.07 | 0.01 | 0.11 | 0.96 | 1517.75 |
| | High_sem | 1.74 | 0.04 | 1 | 0.84 | 1510.29 | | 2.26 | 0.05 | 1 | 0.54 | 1509.37 |
| | Low_phon | 1 | 0.04 | 0.11 | 0.92 | 1513.34 | | 1 | 0.04 | 0.51 | 0.78 | 1513.2 |
| | Low_sem | 2.13 | 0.01 | 1 | 0.5 | 1525.27 | | 1 | 0 | 1 | 0.28 | 1524.62 |
| | RT_high_phon | 1.01 | 0.02 | 1 | 0.07 | 1357.38 | | 1 | 0.01 | 1 | 0.24 | 1360.76 |
| | RT_high_sem | 1 | 0.01 | 0.79 | 0.7 | 1362.81 | | 1 | 0.01 | 1 | 0.59 | 1362.82 |
| | RT_low_phon | 1 | 0.03 | 1 | 0.56 | 1353.49 | | 1 | 0.03 | 0.84 | 0.72 | 1354.18 |
| | RT_low_sem | 1 | 0.01 | 1 | 0.41 | 1359.04 | | 1 | 0.01 | 1 | 0.64 | 1359.56 |
| | Proverb | 5.83 | 0.18 | 1 | 0.22 | 1453.58 | | 1.04 | 0.16 | 0.6 | 0.75 | 1457.91 |
| | Syn_comp | 1.1 | 0.01 | 1 | 0.12 | 1433.99 | | 1.85 | 0.01 | 1 | 0.05 | 1433.44 |
| | Sem_comp | 1 | 0 | 0.25 | 0.77 | 1436.06 | | 1.15 | 0 | 0.07 | 0.97 | 1436.67 |
| | RT_Syn_comp | 1.01 | 0.01 | 0.22 | 0.9 | 1109.91 | | 1 | 0.01 | 0.22 | 0.87 | 1110.14 |
| | RT_Sem_comp | 2.05 | 0.01 | 1 | 0.06 | 1124.41 | | 1 | 0.01 | 1 | 0.14 | 1125.28 |
| | ToT | 3.18 | 0.15 | 1 | 0.21 | 1446.71 | | 1.87 | 0.14 | 1 | 0.2 | 1443.99 |
| **LParsO_RParsO** | VF | 2.73 | 0.24 | 1 | 0.15 | 1459.91 | **LParsO_RParsT** | 1 | 0.23 | 1 | 0.59 | 1463.72 |
| | Vocab | 1.38 | 0.21 | 1 | 0.73 | 1475.86 | | 1 | 0.21 | 0.45 | 0.9 | 1477.19 |
| | High_phon | 1 | 0.01 | 0.56 | 0.75 | 1517.7 | | 1 | 0.02 | 1 | 0.2 | 1515.95 |
| | High_sem | 1.03 | 0.04 | 1 | 0.22 | 1508.61 | | 1 | 0.05 | 1 | 0.08 | 1506.4 |
| | Low_phon | 1.66 | 0.05 | 0.65 | 0.93 | 1512.46 | | 2.7 | 0.04 | 1 | 0.45 | 1513.14 |
| | Low_sem | 1 | 0 | 0.9 | 0.58 | 1524.74 | | 1 | 0 | 1 | 0.26 | 1524.01 |
| | RT_high_phon | 1.92 | 0.01 | 1 | 0.62 | 1362.34 | | 1.19 | 0.01 | 0.54 | 0.84 | 1360.89 |
| | RT_high_sem | 1 | 0.01 | 0.54 | 0.83 | 1362.6 | | 1.52 | 0.01 | 1 | 0.7 | 1363 |
| | RT_low_phon | 1 | 0.03 | 1 | 0.66 | 1353.09 | | 2.62 | 0.04 | 1 | 0.17 | 1350.13 |
| | RT_low_sem | 1 | 0.01 | 1 | 0.45 | 1360.06 | | 1 | 0.01 | 0.22 | 0.94 | 1360.23 |

| | | | | | | | | | | | |
|---|---|---|---|---|---|---|---|---|---|---|---|
| | Proverb | 1 | 0.17 | 0.7 | 0.6 | 1455.11 | | 1 | 0.16 | 0 | 0.99 | 1455.54 |
| | Syn_comp | 2.13 | 0.02 | 1 | 0.01 | 1430.2 | | 1.96 | 0.02 | 1 | 0 | 1427.24 |
| | Sem_comp | 1 | 0 | 0.19 | 0.81 | 1436.56 | | 1 | 0 | 0.07 | 0.93 | 1436.38 |
| | RT_Syn_comp | 1 | 0.01 | 1 | 0.16 | 1108.44 | | 1.32 | 0.01 | 1 | 0.7 | 1110.58 |
| | RT_Sem_comp | 1 | 0.01 | 1 | 0.39 | 1127.12 | | 1 | 0 | 1 | 0.25 | 1128.48 |
| | ToT | 1 | 0.14 | 1 | 0.17 | 1443.77 | | 1 | 0.14 | 1 | 0.28 | 1443.83 |
| **LParsO_RSTG** | VF | 1 | 0.23 | 1 | 0.08 | 1458.89 | **LParsT_RSTS** | 1 | 0.23 | 1 | 0.02 | 1456.41 |
| | Vocab | 1 | 0.21 | 1 | 0.54 | 1477.02 | | 2.23 | 0.22 | 1 | 0.24 | 1473.4 |
| | High_phon | 1 | 0.01 | 0.56 | 0.77 | 1517.41 | | 1 | 0.02 | 0 | 0.99 | 1515.73 |
| | High_sem | 1 | 0.04 | 1 | 0.51 | 1509.43 | | 2.45 | 0.04 | 1 | 0.74 | 1507.28 |
| | Low_phon | 4.45 | 0.05 | 1 | 0.21 | 1511.51 | | 1 | 0.04 | 1 | 0.42 | 1513.33 |
| | Low_sem | 1 | 0 | 0.39 | 0.79 | 1525.61 | | 1 | 0.01 | 0.71 | 0.64 | 1521.83 |
| | RT_high_phon | 1 | 0.03 | 1 | 0.02 | 1354.83 | | 1.22 | 0.01 | 1 | 0.23 | 1360.77 |
| | RT_high_sem | 1 | 0.01 | 1 | 0.09 | 1359.79 | | 1 | 0.01 | 0 | 0.96 | 1362.38 |
| | RT_low_phon | 1 | 0.03 | 1 | 0.25 | 1352.91 | | 2.35 | 0.03 | 1 | 0.5 | 1353.96 |
| | RT_low_sem | 2.32 | 0.03 | 1 | 0.05 | 1352.46 | | 1 | 0.01 | 0.82 | 0.77 | 1360.07 |
| | Proverb | 1.59 | 0.16 | 0.92 | 0.66 | 1461.69 | | 1 | 0.17 | 1 | 0.12 | 1451.21 |
| | Syn_comp | 1 | 0 | 1 | 0.06 | 1435.07 | | 1.38 | 0 | 1 | 0.79 | 1438.14 |
| | Sem_comp | 1 | -0.01 | 0.14 | 0.87 | 1436.77 | | 1 | 0 | 1 | 0.37 | 1435.6 |
| | RT_Syn_comp | 1 | 0.01 | 0.15 | 0.94 | 1110.53 | | 1 | 0.01 | 1 | 0.13 | 1108.3 |
| | RT_Sem_comp | 1 | 0 | 1 | 0.32 | 1125.3 | | 1.14 | 0 | 1 | 0.95 | 1129.7 |
| | ToT | 2.44 | 0.14 | 1 | 0.45 | 1445.28 | | 3.57 | 0.16 | 1 | 0.02 | 1437.83 |
| **LParsT_RMTG** | VF | 2.34 | 0.26 | 1 | 0 | 1445.95 | **LParsT_RParsO** | 2.56 | 0.24 | 1 | 0.2 | 1462.84 |
| | Vocab | 3.74 | 0.23 | 1 | 0.08 | 1467.32 | | 2.05 | 0.21 | 1 | 0.59 | 1475.23 |
| | High_phon | 1.45 | 0.01 | 1 | 0.58 | 1517.73 | | 1 | 0.01 | 1 | 0.55 | 1517.07 |
| | High_sem | 1 | 0.04 | 1 | 0.13 | 1508.07 | | 1 | 0.04 | 1 | 0.12 | 1505.96 |
| | Low_phon | 1 | 0.03 | 0.11 | 0.92 | 1513.97 | | 1 | 0.04 | 0.65 | 0.69 | 1513.11 |
| | Low_sem | 1 | 0.01 | 1 | 0.12 | 1523.65 | | 1 | 0.01 | 1 | 0.24 | 1521.06 |
| | RT_high_phon | 1 | 0.01 | 0.22 | 0.91 | 1361.96 | | 1.81 | 0.02 | 1 | 0.09 | 1357.44 |
| | RT_high_sem | 1.37 | 0.01 | 0.79 | 0.81 | 1362.91 | | 1.41 | 0.01 | 1 | 0.41 | 1360.48 |
| | RT_low_phon | 1 | 0.03 | 1 | 0.51 | 1353.81 | | 1.79 | 0.04 | 1 | 0.14 | 1348.73 |
| | RT_low_sem | 1 | 0.01 | 1 | 0.44 | 1359.36 | | 2.6 | 0.03 | 1 | 0.1 | 1356.03 |
| | Proverb | 2.39 | 0.17 | 1 | 0.32 | 1455.28 | | 1.43 | 0.17 | 0.7 | 0.67 | 1457.98 |
| | Syn_comp | 1.47 | 0 | 1 | 0.68 | 1438.58 | | 1.6 | 0 | 1 | 0.42 | 1436.67 |
| | Sem_comp | 1 | 0 | 1 | 0.38 | 1436.24 | | 1 | -0.01 | 0.14 | 0.85 | 1436.58 |
| | RT_Syn_comp | 1 | 0.01 | 1 | 0.51 | 1109.65 | | 1 | 0.01 | 1 | 0.29 | 1107.41 |
| | RT_Sem_comp | 1.3 | 0.01 | 1 | 0.09 | 1124.9 | | 1 | 0 | 1 | 0.29 | 1128.4 |
| | ToT | 2.9 | 0.15 | 1 | 0.37 | 1443.89 | | 1 | 0.15 | 1 | 0.04 | 1440.76 |
| **LParsT_RParsT** | VF | 2.66 | 0.24 | 1 | 0.03 | 1459.85 | **LParsT_RSTG** | 1.38 | 0.24 | 1 | 0.54 | 1458.64 |
| | Vocab | 1.45 | 0.21 | 1 | 0.68 | 1475.62 | | 2.54 | 0.23 | 1 | 0.01 | 1467.32 |
| | High_phon | 1 | 0.02 | 1 | 0.06 | 1512.36 | | 1 | 0.01 | 0.93 | 0.6 | 1517.02 |
| | High_sem | 1 | 0.05 | 1 | 0.15 | 1500.08 | | 2.31 | 0.06 | 1 | 0.02 | 1497.73 |
| | Low_phon | 1.71 | 0.04 | 1 | 0.35 | 1511.23 | | 1.53 | 0.04 | 1 | 0.61 | 1511.84 |
| | Low_sem | 1 | 0.02 | 1 | 0.05 | 1516.69 | | 1.36 | 0.01 | 0.71 | 0.72 | 1524.59 |
| | RT_high_phon | 1.75 | 0.02 | 1 | 0.1 | 1356.49 | | 1 | 0.02 | 1 | 0.04 | 1357.31 |
| | RT_high_sem | 1 | 0.01 | 1 | 0.42 | 1361.14 | | 2.04 | 0.02 | 1 | 0.36 | 1361.33 |

| | | | | | | | | | | | |
|---|---|---|---|---|---|---|---|---|---|---|---|
| | RT_low_phon | 1.53 | 0.04 | 1 | 0.55 | 1351.98 | | 1.88 | 0.03 | 1 | 0.56 | 1354.06 |
| | RT_low_sem | 1.91 | 0.02 | 1 | 0.14 | 1358.19 | | 1 | 0.01 | 1 | 0.33 | 1358.45 |
| | Proverb | 1.35 | 0.17 | 1 | 0.26 | 1459.52 | | 4.45 | 0.18 | 1 | 0.06 | 1450.32 |
| | Syn_comp | 1.28 | 0 | 1 | 0.45 | 1437.32 | | 1 | 0 | 1 | 0.61 | 1438.44 |
| | Sem_comp | 1 | 0 | 0.16 | 0.84 | 1436.19 | | 1 | 0 | 0 | 0.97 | 1436.58 |
| | RT_Syn_comp | 1.4 | 0.02 | 1 | 0.16 | 1107.48 | | 1 | 0.01 | 1 | 0.11 | 1107.75 |
| | RT_Sem_comp | 1 | 0 | 1 | 0.14 | 1127.37 | | 2.56 | 0 | 1 | 0.33 | 1128.93 |
| | ToT | 2.69 | 0.15 | 1 | 0.14 | 1441.61 | | 3.42 | 0.16 | 1 | 0.07 | 1440.71 |
| **LSTG_RSTS** | VF | 1 | 0.23 | 1 | 0.66 | 1459.36 | **LSTG_RMTG** | 4.2 | 0.24 | 1 | 0.24 | 1465.37 |
| | Vocab | 2.97 | 0.22 | 1 | 0.38 | 1471.95 | | 1.58 | 0.23 | 1 | 0.24 | 1467.85 |
| | High_phon | 1.63 | 0.01 | 1 | 0.35 | 1516.62 | | 1.97 | 0.02 | 1 | 0.24 | 1515.64 |
| | High_sem | 1.21 | 0.04 | 0.76 | 0.7 | 1511.16 | | 2 | 0.05 | 1 | 0.13 | 1507.5 |
| | Low_phon | 2.02 | 0.05 | 1 | 0.07 | 1506.09 | | 3.93 | 0.05 | 1 | 0.25 | 1511.97 |
| | Low_sem | 1 | 0 | 0.71 | 0.64 | 1524.98 | | 1.21 | 0.01 | 1 | 0.32 | 1522.59 |
| | RT_high_phon | 1.66 | 0.02 | 1 | 0.29 | 1359.27 | | 1.81 | 0.03 | 1 | 0.42 | 1356.68 |
| | RT_high_sem | 1 | 0.03 | 1 | 0.02 | 1354.08 | | 1 | 0.03 | 1 | 0.08 | 1356.43 |
| | RT_low_phon | 1.61 | 0.04 | 1 | 0.1 | 1348.73 | | 1 | 0.03 | 1 | 0.34 | 1354.29 |
| | RT_low_sem | 2.02 | 0.03 | 1 | 0.03 | 1353.68 | | 1.64 | 0.02 | 1 | 0.37 | 1358.75 |
| | Proverb | 1 | 0.16 | 0.66 | 0.67 | 1456.98 | | 1 | 0.16 | 0.45 | 0.78 | 1456.7 |
| | Syn_comp | 6.35 | 0.05 | 1 | 0 | 1420.22 | | 6.73 | 0.06 | 1 | 0 | 1416.1 |
| | Sem_comp | 2.31 | 0.01 | 1 | 0.3 | 1434.48 | | 2.93 | 0.02 | 1 | 0.46 | 1431.22 |
| | RT_Syn_comp | 1 | 0.01 | 0.73 | 0.61 | 1109.88 | | 1 | 0.01 | 1 | 0.29 | 1107.97 |
| | RT_Sem_comp | 2.16 | 0.01 | 1 | 0.67 | 1127.59 | | 2.62 | 0 | 1 | 0.42 | 1129.3 |
| | ToT | 3.79 | 0.17 | 1 | 0.01 | 1436.68 | | 3.74 | 0.16 | 1 | 0.05 | 1439.14 |
| **LSTG_RParsO** | VF | 1.05 | 0.23 | 1 | 0.73 | 1465.1 | **LSTG_RParsT** | 3.76 | 0.25 | 1 | 0.02 | 1448.93 |
| | Vocab | 1.71 | 0.21 | 1 | 0.3 | 1472.97 | | 1.82 | 0.22 | 1 | 0.08 | 1469.85 |
| | High_phon | 1 | 0.01 | 0.11 | 0.94 | 1516.57 | | 1.67 | 0.02 | 1 | 0.11 | 1514.1 |
| | High_sem | 1 | 0.04 | 0 | 1 | 1511.93 | | 1.68 | 0.04 | 1 | 0.23 | 1509.43 |
| | Low_phon | 1.14 | 0.04 | 0.65 | 0.87 | 1514.2 | | 2.81 | 0.04 | 1 | 0.47 | 1512.73 |
| | Low_sem | 1.98 | 0.01 | 1 | 0.52 | 1525.21 | | 1 | 0 | 1 | 0.28 | 1524.31 |
| | RT_high_phon | 1 | 0.03 | 1 | 0.15 | 1354.96 | | 1 | 0.02 | 1 | 0.09 | 1358.06 |
| | RT_high_sem | 1.22 | 0.01 | 0.54 | 0.89 | 1363.09 | | 2.65 | 0.01 | 1 | 0.46 | 1363.49 |
| | RT_low_phon | 1.34 | 0.03 | 1 | 0.38 | 1354.11 | | 1.04 | 0.03 | 1 | 0.72 | 1354.63 |
| | RT_low_sem | 1.96 | 0.02 | 1 | 0.13 | 1356.94 | | 2.32 | 0.02 | 1 | 0.61 | 1361.07 |
| | Proverb | 1 | 0.16 | 1 | 0.46 | 1455.77 | | 1.77 | 0.17 | 1 | 0.22 | 1456.3 |
| | Syn_comp | 3.1 | 0.01 | 1 | 0.07 | 1432.83 | | 3.77 | 0.17 | 1 | 0 | 1357.18 |
| | Sem_comp | 1 | 0 | 0.36 | 0.68 | 1436.47 | | 1 | 0 | 0.45 | 0.63 | 1435.79 |
| | RT_Syn_comp | 1 | 0.01 | 0.79 | 0.58 | 1110.17 | | 1 | 0.02 | 1 | 0.02 | 1104.97 |
| | RT_Sem_comp | 1 | -0.01 | 1 | 0.46 | 1129.27 | | 1.5 | 0.01 | 1 | 0.36 | 1126.39 |
| | ToT | 2.43 | 0.14 | 1 | 0.39 | 1445.06 | | 2.83 | 0.15 | 1 | 0.53 | 1442.95 |
| **LSTG_RSTG** | VF | 1.14 | 0.23 | 1 | 0.59 | 1462.88 | | | | | | |
| | Vocab | 1 | 0.22 | 1 | 0.44 | 1471.88 | | | | | | |
| | High_phon | 2.44 | 0.03 | 1 | 0.04 | 1511.11 | | | | | | |
| | High_sem | 2.29 | 0.05 | 1 | 0.11 | 1506.62 | | | | | | |
| | Low_phon | 2.06 | 0.05 | 1 | 0.18 | 1512.42 | | | | | | |
| | Low_sem | 1 | 0.01 | 1 | 0.37 | 1525.72 | | | | | | |
| | RT_high_phon | 1.61 | 0.03 | 1 | 0.54 | 1355.02 | | | | | | |

| | | | | | | |
|---|---|---|---|---|---|---|
| | RT_high_sem | 2.43 | 0.02 | 1 | 0.76 | 1362.16 |
| | RT_low_phon | 1 | 0.03 | 0.81 | 0.76 | 1355.06 |
| | RT_low_sem | 2.23 | 0.03 | 1 | 0.06 | 1352.81 |
| | Proverb | 1 | 0.16 | 0.22 | 0.87 | 1456.79 |
| | Syn_comp | 3.08 | 0.02 | 1 | 0.05 | 1431.16 |
| | Sem_comp | 3.2 | 0.02 | 1 | 0.16 | 1429.62 |
| | RT_Syn_comp | 1 | 0.01 | 1 | 0.42 | 1108.77 |
| | RT_Sem_comp | 3.77 | 0.03 | 1 | 0.04 | 1122.23 |
| | ToT | 3.41 | 0.15 | 1 | 0.25 | 1445.31 |

**Supplementary Table 4.** Generalized Additive Mixed Model (GAMM) results for FC × age interactions across 45 language network edges and 16 language behaviours. Each row represents a specific language behavior for a given pair of SC edges. Reported metrics include the estimated degrees of freedom (EDF) for the interaction term, adjusted R² of the full model (Adj. R²), F statistics value (F value), false discovery rate-corrected p-value (FDR P-value), and Akaike Information Criterion (AIC). FDR correction was applied across all SC edges within each language behavior. Higher edf values indicate more complex, nonlinear relationships, while significant FDR-corrected p-values denote reliable age-by-FC interactions contributing to individual differences in language performance.

| FC edge | Language behavior | EDF | Adj. R2 | F value | FDR P-value | AIC | | FC edge | EDF | Adj. R2 | F value | FDR P-value | AIC |
|---|---|---|---|---|---|---|---|---|---|---|---|---|---|
| LSTS_LMTG | VF | 1 | 0.23 | 0.45 | 0.73 | 1460.15 | | LSTS_LParsO | 1 | 0.23 | 1 | 0.46 | 1460.28 |
| | Vocab | 1 | 0.21 | 0.9 | 0.57 | 1474.34 | | | 1.58 | 0.21 | 0.87 | 0.88 | 1478.17 |
| | High_phon | 1 | 0.01 | 1 | 0.24 | 1516.19 | | | 6.2 | 0.04 | 1 | 0.07 | 1511.55 |
| | High_sem | 1 | 0.04 | 1 | 0.46 | 1508.9 | | | 2.5 | 0.04 | 1 | 0.3 | 1509.69 |
| | Low_phon | 1 | 0.05 | 1 | 0.04 | 1508.87 | | | 1 | 0.04 | 0 | 0.96 | 1513.81 |
| | Low_sem | 1 | 0.01 | 1 | 0.34 | 1523.73 | | | 1 | 0 | 0 | 0.95 | 1525.66 |
| | RT_high_phon | 2.1 | 0.02 | 1 | 0.2 | 1358.9 | | | 2.09 | 0.02 | 1 | 0.26 | 1360.15 |
| | RT_high_sem | 1.17 | 0.02 | 1 | 0.65 | 1357.96 | | | 2.38 | 0.02 | 1 | 0.09 | 1357.24 |
| | RT_low_phon | 2.1 | 0.03 | 1 | 0.5 | 1354.64 | | | 2.95 | 0.04 | 1 | 0.26 | 1353.35 |
| | RT_low_sem | 1 | 0.02 | 1 | 0.24 | 1354.82 | | | 2.04 | 0.02 | 1 | 0.3 | 1358.98 |
| | Proverb | 1 | 0.17 | 1 | 0.09 | 1458.02 | | | 2.28 | 0.17 | 1 | 0.1 | 1450.76 |
| | Syn_comp | 1 | 0.01 | 1 | 0.09 | 1435.81 | | | 1 | 0 | 1 | 0.29 | 1437.63 |
| | Sem_comp | 1.95 | 0 | 0.71 | 0.79 | 1437.49 | | | 1 | 0 | 0.95 | 0.45 | 1436.12 |
| | RT_Syn_comp | 1.11 | 0.01 | 1 | 0.68 | 1109.02 | | | 1 | 0.01 | 1 | 0.56 | 1109.96 |
| | RT_Sem_comp | 1.1 | 0 | 1 | 0.2 | 1128.2 | | | 2.17 | 0 | 1 | 0.35 | 1129.37 |
| | ToT | 2.19 | 0.14 | 1 | 0.69 | 1446.01 | | | 1 | 0.14 | 0.15 | 0.93 | 1445.62 |
| LSTS_LParsT | VF | 6.31 | 0.25 | 1 | 0.08 | 1454.8 | | LSTS_LSTG | 1.88 | 0.23 | 1 | 0.4 | 1461.72 |
| | Vocab | 1.32 | 0.22 | 1 | 0.06 | 1472.21 | | | 2.07 | 0.21 | 1 | 0.42 | 1480.38 |
| | High_phon | 1 | 0.01 | 1 | 0.67 | 1516.49 | | | 4.49 | 0.03 | 1 | 0.23 | 1515.75 |
| | High_sem | 1 | 0.04 | 1 | 0.31 | 1510.37 | | | 2.13 | 0.04 | 1 | 0.21 | 1509.6 |
| | Low_phon | 1.77 | 0.04 | 1 | 0.61 | 1514.29 | | | 1.6 | 0.04 | 1 | 0.64 | 1514.23 |
| | Low_sem | 1 | 0 | 0.98 | 0.59 | 1525.68 | | | 1.77 | 0.01 | 1 | 0.3 | 1523.29 |
| | RT_high_phon | 1.7 | 0.02 | 1 | 0.32 | 1360.14 | | | 3.58 | 0.02 | 1 | 0.49 | 1359.3 |
| | RT_high_sem | 1 | 0.03 | 1 | 0.56 | 1353.11 | | | 1.71 | 0.02 | 1 | 0.21 | 1357.05 |
| | RT_low_phon | 2.57 | 0.05 | 1 | 0.01 | 1344.75 | | | 2.19 | 0.04 | 1 | 0.2 | 1351.08 |
| | RT_low_sem | 1.61 | 0.03 | 1 | 0.02 | 1351.79 | | | 1.98 | 0.02 | 1 | 0.24 | 1358.42 |
| | Proverb | 1.86 | 0.17 | 1 | 0.12 | 1452.69 | | | 1 | 0.16 | 1 | 0.17 | 1455.1 |
| | Syn_comp | 1 | 0 | 0.56 | 0.75 | 1438.29 | | | 2.81 | 0 | 1 | 0.45 | 1438.16 |
| | Sem_comp | 1 | 0 | 0.9 | 0.48 | 1435.31 | | | 1 | 0 | 1 | 0.33 | 1435.65 |
| | RT_Syn_comp | 1.6 | 0.01 | 1 | 0.65 | 1110.93 | | | 1 | 0.02 | 1 | 0.04 | 1106.27 |
| | RT_Sem_comp | 1 | 0 | 0.5 | 0.78 | 1128.86 | | | 5.28 | 0.04 | 1 | 0.2 | 1119.31 |

| | | | | | | | | | | | | |
|---|---|---|---|---|---|---|---|---|---|---|---|---|
| | ToT | 1 | 0.14 | 1 | 0.23 | 1444.16 | | 1 | 0.15 | 1 | 0.05 | 1445.27 |
| **LMTG_LParsO** | VF | 1 | 0.24 | 1 | 0.27 | 1458 | **LMTG_LParsT** | 1.85 | 0.24 | 1 | 0.16 | 1457.57 |
| | Vocab | 1.6 | 0.22 | 1 | 0.59 | 1474.11 | | 1 | 0.21 | 1 | 0.04 | 1470.97 |
| | High_phon | 1 | 0.02 | 1 | 0.04 | 1513.29 | | 3.54 | 0.03 | 1 | 0.45 | 1514.05 |
| | High_sem | 1.13 | 0.07 | 1 | 0 | 1495.08 | | 3.43 | 0.07 | 1 | 0 | 1496.21 |
| | Low_phon | 1 | 0.04 | 1 | 0.31 | 1512 | | 1 | 0.04 | 1 | 0.44 | 1513.01 |
| | Low_sem | 1 | 0 | 1 | 0.27 | 1524.42 | | 1 | 0 | 0.36 | 0.85 | 1525.2 |
| | RT_high_phon | 1 | 0.03 | 1 | 0.3 | 1356.55 | | 1 | 0.01 | 1 | 0.7 | 1359.87 |
| | RT_high_sem | 1 | 0.01 | 1 | 0.37 | 1361.76 | | 1 | 0.01 | 1 | 0.65 | 1362.47 |
| | RT_low_phon | 4.86 | 0.06 | 1 | 0.39 | 1349.16 | | 3.76 | 0.04 | 1 | 0.4 | 1353.12 |
| | RT_low_sem | 5.38 | 0.04 | 1 | 0.11 | 1353.55 | | 3.37 | 0.03 | 1 | 0.26 | 1357.73 |
| | Proverb | 1.96 | 0.16 | 1 | 0.43 | 1455.79 | | 1.6 | 0.16 | 1 | 0.38 | 1457.52 |
| | Syn_comp | 1.43 | 0 | 0.67 | 0.87 | 1438.89 | | 1 | 0 | 0.09 | 0.93 | 1438.65 |
| | Sem_comp | 1 | 0 | 0.25 | 0.82 | 1436.33 | | 1 | 0 | 0.71 | 0.57 | 1436.24 |
| | RT_Syn_comp | 1 | 0.02 | 1 | 0.23 | 1107.24 | | 1.71 | 0.02 | 1 | 0.53 | 1108.65 |
| | RT_Sem_comp | 1 | 0.01 | 0.93 | 0.58 | 1127.11 | | 1.94 | 0 | 1 | 0.35 | 1128.84 |
| | ToT | 2.84 | 0.15 | 1 | 0.16 | 1442.42 | | 2.22 | 0.15 | 1 | 0.17 | 1444.87 |
| **LMTG_LSTG** | VF | 1 | 0.23 | 1 | 0.27 | 1459.66 | **LParsO_LParsT** | 1 | 0.23 | 1 | 0.44 | 1460.93 |
| | Vocab | 1.39 | 0.22 | 1 | 0.45 | 1474.97 | | 3.85 | 0.22 | 1 | 0.17 | 1478.28 |
| | High_phon | 1.59 | 0.02 | 1 | 0.49 | 1516.1 | | 3.14 | 0.02 | 1 | 0.43 | 1516.36 |
| | High_sem | 1.6 | 0.04 | 1 | 0.68 | 1508.63 | | 1.44 | 0.04 | 1 | 0.19 | 1509.45 |
| | Low_phon | 2.48 | 0.04 | 1 | 0.67 | 1514.71 | | 4.84 | 0.07 | 1 | 0.15 | 1506.86 |
| | Low_sem | 2.57 | 0.01 | 0.98 | 0.83 | 1527.32 | | 1 | 0 | 1 | 0.52 | 1524.18 |
| | RT_high_phon | 1 | 0.01 | 1 | 0.75 | 1361.66 | | 1.64 | 0.02 | 1 | 0.25 | 1360.47 |
| | RT_high_sem | 1 | 0.01 | 0.9 | 0.78 | 1363.2 | | 1 | 0.01 | 0.45 | 0.93 | 1362.92 |
| | RT_low_phon | 1.41 | 0.03 | 1 | 0.4 | 1353.6 | | 1 | 0.03 | 1 | 0.42 | 1353.79 |
| | RT_low_sem | 1 | 0.01 | 0.45 | 0.83 | 1360.08 | | 1 | 0.02 | 0 | 0.99 | 1359.36 |
| | Proverb | 1 | 0.17 | 1 | 0 | 1447.21 | | 1 | 0.18 | 1 | 0.01 | 1448.5 |
| | Syn_comp | 1 | 0 | 0 | 0.97 | 1438.9 | | 1 | 0 | 1 | 0.34 | 1437.66 |
| | Sem_comp | 1 | 0 | 0.19 | 0.87 | 1436.31 | | 1 | 0 | 0.11 | 0.9 | 1435.83 |
| | RT_Syn_comp | 1 | 0.01 | 1 | 0.47 | 1110.2 | | 1.88 | 0.02 | 1 | 0.4 | 1108.16 |
| | RT_Sem_comp | 2.08 | 0.01 | 1 | 0.31 | 1124.91 | | 1 | 0 | 1 | 0.35 | 1127.52 |
| | ToT | 1 | 0.14 | 1 | 0.6 | 1444.65 | | 1 | 0.15 | 1 | 0.01 | 1438.05 |
| **LParsO_LSTG** | VF | 1 | 0.23 | 1 | 0.33 | 1460.13 | **LParsT_LSTG** | 1 | 0.23 | 0.16 | 0.85 | 1461.58 |
| | Vocab | 1.08 | 0.21 | 0.45 | 0.98 | 1478.04 | | 1.96 | 0.22 | 1 | 0.28 | 1472.98 |
| | High_phon | 1 | 0.02 | 1 | 0.34 | 1515.96 | | 1 | 0.02 | 0 | 0.96 | 1517.07 |
| | High_sem | 1 | 0.05 | 1 | 0.02 | 1505.15 | | 1 | 0.05 | 1 | 0 | 1502.57 |
| | Low_phon | 1 | 0.05 | 1 | 0.56 | 1510.84 | | 1 | 0.04 | 1 | 0.61 | 1512.54 |
| | Low_sem | 1 | 0 | 0 | 0.97 | 1525.47 | | 1.44 | 0.03 | 1 | 0.2 | 1518.46 |
| | RT_high_phon | 1.91 | 0.01 | 1 | 0.43 | 1361.68 | | 2.2 | 0.02 | 1 | 0.16 | 1359.05 |
| | RT_high_sem | 1.42 | 0.01 | 1 | 0.7 | 1363.15 | | 3.34 | 0.03 | 1 | 0.08 | 1356.44 |
| | RT_low_phon | 1.21 | 0.03 | 0.6 | 0.9 | 1355.28 | | 3.32 | 0.04 | 1 | 0.45 | 1353.54 |
| | RT_low_sem | 2.33 | 0.02 | 1 | 0.14 | 1356.82 | | 4.28 | 0.03 | 1 | 0.37 | 1359.74 |
| | Proverb | 1 | 0.16 | 1 | 0.35 | 1456.01 | | 2.93 | 0.17 | 1 | 0.12 | 1452.33 |
| | Syn_comp | 1 | 0 | 1 | 0.47 | 1438.15 | | 1.95 | 0.02 | 1 | 0.24 | 1432.51 |
| | Sem_comp | 1 | 0 | 1 | 0.05 | 1431.57 | | 1 | 0 | 1 | 0.22 | 1434.46 |

| | | | | | | | | | | | | |
|---|---|---|---|---|---|---|---|---|---|---|---|---|
| | RT_Syn_comp | 1.9 | 0.02 | 1 | 0.67 | 1108.82 | | 1.4 | 0.01 | 1 | 0.61 | 1110.46 |
| | RT_Sem_comp | 1.03 | 0 | 1 | 0.3 | 1129.13 | | 2.4 | 0.01 | 1 | 0.31 | 1127.99 |
| | ToT | 2.37 | 0.15 | 1 | 0.51 | 1446.56 | | 3.01 | 0.16 | 1 | 0.14 | 1442.45 |
| **RSTS_RMTG** | VF | 2.04 | 0.23 | 0.16 | 0.99 | 1465.58 | **RSTS_RParsO** | 1 | 0.23 | 0.16 | 0.84 | 1460.02 |
| | Vocab | 1 | 0.21 | 1 | 0.22 | 1472.99 | | 1.4 | 0.21 | 0.87 | 0.79 | 1476.72 |
| | High_phon | 1 | 0.03 | 1 | 0.01 | 1508.75 | | 3.08 | 0.03 | 1 | 0.05 | 1511.54 |
| | High_sem | 1.07 | 0.04 | 1 | 0 | 1498.76 | | 3.22 | 0.04 | 1 | 0.43 | 1510.94 |
| | Low_phon | 1 | 0.04 | 1 | 0.02 | 1507.78 | | 1 | 0.04 | 0.22 | 0.91 | 1512.73 |
| | Low_sem | 1.01 | 0.01 | 1 | 0.01 | 1517.56 | | 1.41 | 0 | 0.98 | 0.79 | 1526 |
| | RT_high_phon | 1.3 | 0.01 | 1 | 0.8 | 1361.41 | | 1 | 0.02 | 1 | 0.26 | 1359.16 |
| | RT_high_sem | 1.89 | 0.01 | 1 | 0.28 | 1362.05 | | 1.48 | 0.01 | 1 | 0.23 | 1361.12 |
| | RT_low_phon | 1.56 | 0.03 | 1 | 0.54 | 1354.75 | | 1.1 | 0.03 | 0.99 | 0.89 | 1355.04 |
| | RT_low_sem | 2.74 | 0.03 | 1 | 0.07 | 1354.14 | | 1.89 | 0.02 | 1 | 0.32 | 1359.52 |
| | Proverb | 1 | 0.16 | 1 | 0.48 | 1454.85 | | 1.76 | 0.17 | 1 | 0.49 | 1454.47 |
| | Syn_comp | 1 | 0 | 1 | 0.38 | 1434.32 | | 3.28 | 0.03 | 1 | 0.02 | 1427.77 |
| | Sem_comp | 1.51 | 0 | 0.71 | 0.68 | 1436.08 | | 1 | -0.01 | 0.25 | 0.83 | 1436.58 |
| | RT_Syn_comp | 2.65 | 0.03 | 1 | 0.18 | 1105.11 | | 1.74 | 0.03 | 1 | 0.51 | 1103.1 |
| | RT_Sem_comp | 3.37 | 0.02 | 1 | 0.09 | 1123.46 | | 1 | -0.01 | 0.5 | 0.75 | 1129.8 |
| | ToT | 2.61 | 0.15 | 1 | 0.04 | 1440 | | 1.67 | 0.14 | 1 | 0.51 | 1445.61 |
| **RSTS_RParsT** | VF | 1 | 0.23 | 1 | 0.17 | 1459.08 | **RSTS_RSTG** | 1 | 0.23 | 1 | 0.39 | 1456.4 |
| | Vocab | 1.59 | 0.21 | 1 | 0.62 | 1476.07 | | 1 | 0.21 | 0.87 | 0.66 | 1474.24 |
| | High_phon | 1 | 0.01 | 1 | 0.82 | 1515.88 | | 2.25 | 0.02 | 1 | 0.8 | 1518.18 |
| | High_sem | 3.48 | 0.07 | 1 | 0.01 | 1497.58 | | 1 | 0.03 | 1 | 0.63 | 1511.61 |
| | Low_phon | 3.87 | 0.05 | 1 | 0.22 | 1512.24 | | 1.59 | 0.04 | 1 | 0.32 | 1513.27 |
| | Low_sem | 2.43 | 0.02 | 1 | 0.18 | 1517.05 | | 1.41 | 0 | 0.98 | 0.76 | 1525.5 |
| | RT_high_phon | 1.86 | 0.02 | 1 | 0.21 | 1358.48 | | 2.37 | 0.02 | 1 | 0.17 | 1358.59 |
| | RT_high_sem | 1 | 0.01 | 0.9 | 0.81 | 1361.99 | | 1.01 | 0.01 | 1 | 0.58 | 1362.67 |
| | RT_low_phon | 2.32 | 0.05 | 1 | 0.34 | 1348.39 | | 1 | 0.03 | 1 | 0.17 | 1352.85 |
| | RT_low_sem | 2.7 | 0.02 | 1 | 0.76 | 1361.43 | | 6.75 | 0.05 | 1 | 0.11 | 1353.38 |
| | Proverb | 1.86 | 0.16 | 1 | 0.7 | 1463.74 | | 1 | 0.16 | 0 | 0.96 | 1456.01 |
| | Syn_comp | 5.31 | 0.03 | 1 | 0.02 | 1428 | | 1 | 0 | 0.09 | 0.94 | 1438.31 |
| | Sem_comp | 1 | 0 | 0.34 | 0.77 | 1437.02 | | 1.52 | 0 | 1 | 0.4 | 1435.44 |
| | RT_Syn_comp | 1 | 0.01 | 1 | 0.68 | 1109.97 | | 1.46 | 0.01 | 1 | 0.66 | 1110.87 |
| | RT_Sem_comp | 1.28 | 0 | 1 | 0.3 | 1127.75 | | 1 | 0 | 1 | 0.05 | 1125.72 |
| | ToT | 1 | 0.14 | 0 | 0.97 | 1445.48 | | 1 | 0.15 | 1 | 0.02 | 1439.73 |
| **RMTG_RParsO** | VF | 1 | 0.23 | 0.51 | 0.68 | 1461.02 | **RMTG_RParsT** | 1 | 0.23 | 0.16 | 0.87 | 1461.25 |
| | Vocab | 1 | 0.21 | 0.56 | 0.81 | 1474.09 | | 1.97 | 0.22 | 1 | 0.12 | 1471.03 |
| | High_phon | 2.51 | 0.02 | 1 | 0.13 | 1514.22 | | 2.88 | 0.03 | 1 | 0.03 | 1509.84 |
| | High_sem | 3.27 | 0.05 | 1 | 0.62 | 1510.63 | | 4.24 | 0.05 | 1 | 0.3 | 1508.54 |
| | Low_phon | 1.83 | 0.04 | 1 | 0.41 | 1512.97 | | 3.52 | 0.06 | 1 | 0.02 | 1505.66 |
| | Low_sem | 1.7 | 0.01 | 1 | 0.48 | 1524.81 | | 1.58 | 0.01 | 1 | 0.37 | 1524.99 |
| | RT_high_phon | 3.43 | 0.03 | 1 | 0.07 | 1357.19 | | 2.55 | 0.02 | 1 | 0.69 | 1360.96 |
| | RT_high_sem | 4.2 | 0.03 | 1 | 0.26 | 1359.98 | | 5.75 | 0.05 | 1 | 0.02 | 1352.48 |
| | RT_low_phon | 2.6 | 0.04 | 1 | 0.03 | 1348.32 | | 1.39 | 0.03 | 1 | 0.57 | 1354.88 |
| | RT_low_sem | 3.53 | 0.03 | 1 | 0.09 | 1355.48 | | 3.87 | 0.03 | 1 | 0.19 | 1357.01 |
| | Proverb | 2.08 | 0.18 | 1 | 0.23 | 1450.18 | | 3.18 | 0.18 | 1 | 0.01 | 1444.79 |
| | Syn_comp | 2.14 | 0.05 | 1 | 0.21 | 1417.18 | | 1.32 | 0 | 1 | 0.17 | 1435.42 |

| | | | | | | | | | | | | | |
|---|---|---|---|---|---|---|---|---|---|---|---|---|---|
| | Sem_comp | 4.13 | 0.01 | 1 | 0.37 | 1433.9 | | | 1 | 0 | 0.15 | 0.9 | 1436.17 |
| | RT_Syn_comp | 3.34 | 0.02 | 1 | 0.39 | 1108.83 | | | 1 | 0.01 | 1 | 0.58 | 1110.17 |
| | RT_Sem_comp | 1 | 0 | 0.63 | 0.71 | 1127.65 | | | 1 | -0.01 | 0.74 | 0.63 | 1129.27 |
| | ToT | 1 | 0.14 | 1 | 0.67 | 1444.42 | | | 1.91 | 0.15 | 1 | 0.57 | 1444.56 |
| **RMTG_RSTG** | VF | 1 | 0.24 | 0.16 | 0.87 | 1455.29 | | **RParsO_RParsT** | 1 | 0.23 | 1 | 0.34 | 1462.72 |
| | Vocab | 3.91 | 0.22 | 1 | 0.61 | 1476.31 | | | 1.48 | 0.23 | 0.87 | 0.72 | 1471.35 |
| | High_phon | 3.17 | 0.03 | 1 | 0.37 | 1514.28 | | | 1.25 | 0.01 | 1 | 0.7 | 1517.65 |
| | High_sem | 2.5 | 0.04 | 1 | 0.65 | 1512.38 | | | 2.37 | 0.04 | 1 | 0.26 | 1509.02 |
| | Low_phon | 4.99 | 0.05 | 1 | 0.19 | 1510.55 | | | 2.96 | 0.04 | 1 | 0.59 | 1514.78 |
| | Low_sem | 1 | 0 | 1 | 0.44 | 1524.2 | | | 4.32 | 0.02 | 1 | 0.31 | 1523.4 |
| | RT_high_phon | 3.48 | 0.03 | 1 | 0.09 | 1356.79 | | | 3.84 | 0.04 | 1 | 0.02 | 1353.09 |
| | RT_high_sem | 1.4 | 0.01 | 1 | 0.69 | 1362.13 | | | 7.26 | 0.07 | 1 | 0 | 1346.9 |
| | RT_low_phon | 5.19 | 0.06 | 1 | 0.14 | 1345.32 | | | 2.3 | 0.04 | 1 | 0.16 | 1351.45 |
| | RT_low_sem | 1 | 0.06 | 1 | 0.42 | 1341.86 | | | 3.99 | 0.03 | 1 | 0.33 | 1358.19 |
| | Proverb | 1 | 0.16 | 1 | 0.54 | 1455.52 | | | 1.91 | 0.16 | 1 | 0.4 | 1456.53 |
| | Syn_comp | 3.19 | 0.03 | 1 | 0.03 | 1427.09 | | | 1 | 0.01 | 0.15 | 0.88 | 1436.16 |
| | Sem_comp | 1 | 0 | 0.88 | 0.49 | 1436.21 | | | 1.53 | 0 | 0.61 | 0.83 | 1436.99 |
| | RT_Syn_comp | 3.27 | 0.02 | 1 | 0.61 | 1112.36 | | | 1.54 | 0.01 | 1 | 0.64 | 1109.56 |
| | RT_Sem_comp | 1.13 | 0 | 1 | 0.15 | 1127.57 | | | 1 | 0.02 | 1 | 0.21 | 1124.03 |
| | ToT | 1.56 | 0.15 | 1 | 0.08 | 1442.98 | | | 1 | 0.14 | 1 | 0.22 | 1443.21 |
| **RParsO_RSTG** | VF | 1 | 0.23 | 0.16 | 0.84 | 1461.22 | | **RParsT_RSTG** | 1 | 0.23 | 0.16 | 0.84 | 1460.95 |
| | Vocab | 1 | 0.21 | 0.56 | 0.77 | 1474.38 | | | 1 | 0.21 | 0.56 | 0.87 | 1474.61 |
| | High_phon | 6.77 | 0.05 | 1 | 0.03 | 1508.32 | | | 1.25 | 0.02 | 1 | 0.17 | 1515.2 |
| | High_sem | 1 | 0.05 | 1 | 0.05 | 1506.37 | | | 1 | 0.03 | 1 | 0.76 | 1510.64 |
| | Low_phon | 5.79 | 0.06 | 1 | 0.18 | 1508.68 | | | 1 | 0.04 | 1 | 0.33 | 1513.29 |
| | Low_sem | 1.81 | 0 | 1 | 0.7 | 1525.94 | | | 1 | 0.01 | 0.98 | 0.61 | 1524.3 |
| | RT_high_phon | 2.27 | 0.02 | 1 | 0.27 | 1358.95 | | | 1.63 | 0.02 | 1 | 0.33 | 1361.05 |
| | RT_high_sem | 1.37 | 0.02 | 1 | 0.11 | 1359.71 | | | 1.47 | 0.02 | 1 | 0.19 | 1360.95 |
| | RT_low_phon | 1.71 | 0.03 | 1 | 0.44 | 1354.55 | | | 4.13 | 0.04 | 1 | 0.47 | 1355.4 |
| | RT_low_sem | 1.27 | 0.01 | 1 | 0.56 | 1360.01 | | | 1 | 0.01 | 0.45 | 0.83 | 1360.4 |
| | Proverb | 3.7 | 0.18 | 1 | 0.43 | 1450.53 | | | 1 | 0.16 | 1 | 0.54 | 1456.16 |
| | Syn_comp | 1 | 0 | 1 | 0.33 | 1437.75 | | | 1.58 | 0 | 1 | 0.56 | 1438.69 |
| | Sem_comp | 1 | 0 | 0.86 | 0.51 | 1436.1 | | | 1 | 0 | 0.35 | 0.75 | 1436.33 |
| | RT_Syn_comp | 2.57 | 0.02 | 1 | 0.65 | 1110.74 | | | 1 | 0.01 | 1 | 0.14 | 1108.03 |
| | RT_Sem_comp | 2.99 | 0.01 | 1 | 0.23 | 1127.31 | | | 1 | -0.01 | 1 | 0.49 | 1129.46 |
| | ToT | 1.37 | 0.14 | 1 | 0.78 | 1446.93 | | | 2.6 | 0.15 | 1 | 0.21 | 1444.3 |
| **LSTS_RSTS** | VF | 2.21 | 0.23 | 1 | 0.2 | 1459.92 | | **LSTS_RMTG** | 1.61 | 0.23 | 1 | 0.57 | 1466.55 |
| | Vocab | 1 | 0.21 | 1 | 0.32 | 1470.96 | | | 1 | 0.21 | 0.56 | 0.75 | 1476.86 |
| | High_phon | 5.13 | 0.03 | 1 | 0.13 | 1513.53 | | | 1.09 | 0.02 | 1 | 0.15 | 1515.23 |
| | High_sem | 1 | 0.04 | 1 | 0.04 | 1506.82 | | | 1.54 | 0.04 | 1 | 0.65 | 1512.48 |
| | Low_phon | 1 | 0.04 | 1 | 0.04 | 1510.15 | | | 1 | 0.04 | 1 | 0.08 | 1510.39 |
| | Low_sem | 1 | 0 | 0.98 | 0.56 | 1525.36 | | | 1 | 0 | 0.22 | 0.9 | 1525.59 |
| | RT_high_phon | 1.56 | 0.01 | 1 | 0.71 | 1361.79 | | | 2.81 | 0.03 | 1 | 0.31 | 1359.18 |
| | RT_high_sem | 2.07 | 0.03 | 1 | 0.16 | 1357.47 | | | 3.71 | 0.03 | 1 | 0.2 | 1357.62 |
| | RT_low_phon | 1 | 0.06 | 1 | 0.23 | 1347.09 | | | 2.1 | 0.03 | 1 | 0.51 | 1354.65 |
| | RT_low_sem | 1 | 0.01 | 0.22 | 0.92 | 1360.2 | | | 1 | 0.03 | 1 | 0.04 | 1354.66 |

| | | | | | | | | | | | | |
|---|---|---|---|---|---|---|---|---|---|---|---|---|
| | Proverb | 5.76 | 0.19 | 1 | 0.41 | 1448.73 | | 2.07 | 0.18 | 1 | 0.42 | 1449.06 |
| | Syn_comp | 1 | 0 | 0 | 0.98 | 1436.01 | | 1.76 | 0.01 | 1 | 0.27 | 1434.72 |
| | Sem_comp | 1 | 0 | 0.69 | 0.61 | 1434.5 | | 1 | 0 | 1 | 0.17 | 1434.89 |
| | RT_Syn_comp | 2.32 | 0.03 | 1 | 0.31 | 1101.53 | | 3.37 | 0.02 | 1 | 0.7 | 1110.22 |
| | RT_Sem_comp | 1 | 0 | 0.5 | 0.76 | 1128.27 | | 1.08 | 0 | 0.22 | 0.98 | 1127.82 |
| | ToT | 3.31 | 0.15 | 1 | 0.25 | 1444.51 | | 1.47 | 0.14 | 1 | 0.26 | 1445.1 |
| **LSTS_RParsO** | VF | 1 | 0.24 | 1 | 0.05 | 1455.24 | **LSTS_RParsT** | 3.11 | 0.25 | 1 | 0.05 | 1451.7 |
| | Vocab | 3.94 | 0.23 | 1 | 0.57 | 1476.49 | | 1 | 0.21 | 1 | 0.47 | 1473.74 |
| | High_phon | 3.27 | 0.02 | 1 | 0.63 | 1518.37 | | 1 | 0.01 | 1 | 0.64 | 1516.44 |
| | High_sem | 2.04 | 0.04 | 1 | 0.1 | 1506.55 | | 1.82 | 0.04 | 1 | 0.62 | 1510.94 |
| | Low_phon | 1.98 | 0.04 | 1 | 0.42 | 1511.86 | | 2.17 | 0.05 | 1 | 0.14 | 1510.93 |
| | Low_sem | 1.92 | 0.01 | 1 | 0.45 | 1525.18 | | 1 | 0.01 | 0.15 | 0.93 | 1524.12 |
| | RT_high_phon | 1 | 0.02 | 1 | 0.04 | 1355.48 | | 1.09 | 0.03 | 1 | 0.01 | 1354.55 |
| | RT_high_sem | 2.49 | 0.03 | 1 | 0.08 | 1355.32 | | 3.58 | 0.04 | 1 | 0.01 | 1351.02 |
| | RT_low_phon | 1 | 0.03 | 0.6 | 0.85 | 1352.67 | | 4.52 | 0.05 | 1 | 0.19 | 1351.35 |
| | RT_low_sem | 4.92 | 0.03 | 1 | 0.44 | 1356.22 | | 5.34 | 0.03 | 1 | 0.39 | 1359.26 |
| | Proverb | 3.46 | 0.17 | 1 | 0.58 | 1456.56 | | 1 | 0.16 | 1 | 0.39 | 1455.9 |
| | Syn_comp | 1.22 | 0.01 | 1 | 0.35 | 1434.27 | | 1 | 0 | 1 | 0.42 | 1438 |
| | Sem_comp | 1 | 0 | 0.69 | 0.63 | 1436.12 | | 1 | 0 | 0.71 | 0.59 | 1436.16 |
| | RT_Syn_comp | 1 | 0.01 | 0.34 | 0.86 | 1110.6 | | 1 | 0.01 | 1 | 0.55 | 1110.13 |
| | RT_Sem_comp | 1.88 | 0 | 1 | 0.3 | 1128.58 | | 2.26 | 0 | 1 | 0.54 | 1130.33 |
| | ToT | 1.1 | 0.14 | 1 | 0.79 | 1448.17 | | 1 | 0.14 | 1 | 0.33 | 1445.1 |
| **LSTS_RSTG** | VF | 2.26 | 0.23 | 1 | 0.6 | 1463.2 | **LMTG_RSTS** | 1 | 0.23 | 1 | 0.12 | 1461.52 |
| | Vocab | 1.52 | 0.21 | 1 | 0.21 | 1473.95 | | 1 | 0.21 | 0.87 | 0.59 | 1473.88 |
| | High_phon | 3.5 | 0.03 | 1 | 0.06 | 1508.8 | | 2.54 | 0.02 | 1 | 0.13 | 1514.36 |
| | High_sem | 3.76 | 0.06 | 1 | 0.01 | 1503.6 | | 2.51 | 0.05 | 1 | 0.04 | 1505.53 |
| | Low_phon | 2.05 | 0.04 | 1 | 0.76 | 1513.07 | | 2.61 | 0.06 | 1 | 0 | 1504.31 |
| | Low_sem | 1.22 | 0.01 | 1 | 0.17 | 1518.79 | | 2.54 | 0.01 | 1 | 0.39 | 1525.37 |
| | RT_high_phon | 1.47 | 0.02 | 1 | 0.3 | 1359.73 | | 3.14 | 0.02 | 1 | 0.58 | 1362.86 |
| | RT_high_sem | 1.4 | 0.01 | 1 | 0.84 | 1363.32 | | 1.31 | 0.01 | 1 | 0.5 | 1362.5 |
| | RT_low_phon | 1.04 | 0.03 | 1 | 0.27 | 1353.8 | | 1.76 | 0.05 | 1 | 0.3 | 1347.04 |
| | RT_low_sem | 1 | 0.02 | 1 | 0.09 | 1357.64 | | 1 | 0.01 | 0.75 | 0.7 | 1358.96 |
| | Proverb | 1 | 0.17 | 1 | 0.74 | 1452.8 | | 1 | 0.16 | 0.45 | 0.88 | 1456.32 |
| | Syn_comp | 1 | 0.01 | 1 | 0.12 | 1432.42 | | 1 | 0 | 0.82 | 0.66 | 1438.2 |
| | Sem_comp | 1 | 0 | 1 | 0.41 | 1436 | | 1 | 0 | 1 | 0.22 | 1435.06 |
| | RT_Syn_comp | 1.95 | 0.01 | 1 | 0.48 | 1109.64 | | 2.31 | 0.04 | 1 | 0.35 | 1107.42 |
| | RT_Sem_comp | 1.34 | -0.01 | 0.73 | 0.82 | 1130.26 | | 1 | 0 | 1 | 0.52 | 1129.58 |
| | ToT | 1 | 0.14 | 1 | 0.38 | 1445.52 | | 1.07 | 0.14 | 1 | 0.59 | 1446.82 |
| **LMTG_RMTG** | VF | 4.36 | 0.24 | 1 | 0.42 | 1455.29 | **LMTG_RParsO** | 1 | 0.24 | 1 | 0.08 | 1455.02 |
| | Vocab | 1.15 | 0.21 | 1 | 0.38 | 1476.77 | | 1.21 | 0.22 | 1 | 0.35 | 1474.9 |
| | High_phon | 1 | 0.02 | 1 | 0.2 | 1512.82 | | 3.04 | 0.03 | 1 | 0.03 | 1510.28 |
| | High_sem | 2.04 | 0.04 | 1 | 0.81 | 1510.84 | | 2 | 0.04 | 1 | 0.39 | 1509.67 |
| | Low_phon | 1.56 | 0.04 | 1 | 0.91 | 1514.47 | | 1 | 0.04 | 1 | 0.35 | 1509.97 |
| | Low_sem | 3.15 | 0.02 | 1 | 0.09 | 1520.47 | | 1 | 0 | 0.98 | 0.54 | 1524.56 |
| | RT_high_phon | 2.2 | 0.03 | 1 | 0.24 | 1354.09 | | 1.96 | 0.01 | 1 | 0.69 | 1362.8 |
| | RT_high_sem | 2.16 | 0.03 | 1 | 0.2 | 1352.43 | | 1.57 | 0.01 | 1 | 0.62 | 1363.2 |
| | RT_low_phon | 1 | 0.03 | 1 | 0.44 | 1353.36 | | 2.34 | 0.06 | 1 | 0.18 | 1345.34 |

| | | | | | | | | | | | | |
|---|---|---|---|---|---|---|---|---|---|---|---|---|
| | RT_low_sem | 1.41 | 0.03 | 1 | 0.49 | 1353.01 | | 4.4 | 0.04 | 1 | 0.05 | 1353.08 |
| | Proverb | 1 | 0.17 | 1 | 0.07 | 1452.71 | | 6.25 | 0.19 | 1 | 0.26 | 1443.84 |
| | Syn_comp | 1 | 0 | 1 | 0.29 | 1437.61 | | 1.33 | 0 | 0.65 | 0.9 | 1438.63 |
| | Sem_comp | 1 | 0 | 0.82 | 0.53 | 1436.25 | | 1 | 0 | 1 | 0.35 | 1434.15 |
| | RT_Syn_comp | 6.61 | 0.05 | 1 | 0.09 | 1104.93 | | 1 | 0.01 | 1 | 0.51 | 1109.98 |
| | RT_Sem_comp | 1 | 0.01 | 0.22 | 0.88 | 1127.65 | | 2.01 | 0.01 | 1 | 0.1 | 1124.39 |
| | ToT | 1 | 0.15 | 1 | 0.02 | 1439.93 | | 1.67 | 0.15 | 1 | 0.05 | 1441.65 |
| **LMTG_RParsT** | VF | 1.13 | 0.25 | 0.16 | 0.96 | 1449 | **LMTG_RSTG** | 1.04 | 0.24 | 0.98 | 0.57 | 1463.56 |
| | Vocab | 1.82 | 0.22 | 1 | 0.15 | 1470.35 | | 2.37 | 0.21 | 0.87 | 0.87 | 1476.11 |
| | High_phon | 3.04 | 0.02 | 1 | 0.16 | 1513.53 | | 1.24 | 0.02 | 1 | 0.35 | 1514.35 |
| | High_sem | 2.76 | 0.05 | 1 | 0.04 | 1504.56 | | 1 | 0.05 | 1 | 0.22 | 1504.34 |
| | Low_phon | 1.66 | 0.04 | 1 | 0.18 | 1511.83 | | 1 | 0.04 | 0.9 | 0.81 | 1513.23 |
| | Low_sem | 1.97 | 0.01 | 1 | 0.69 | 1523.9 | | 1.53 | 0 | 1 | 0.48 | 1525.09 |
| | RT_high_phon | 1 | 0.03 | 1 | 0.18 | 1351.98 | | 1 | 0.02 | 1 | 0.22 | 1359.83 |
| | RT_high_sem | 1 | 0.01 | 1 | 0.53 | 1360.89 | | 1.62 | 0.01 | 1 | 0.26 | 1361.54 |
| | RT_low_phon | 1 | 0.03 | 0.45 | 0.91 | 1353.7 | | 1.87 | 0.07 | 1 | 0.45 | 1343.21 |
| | RT_low_sem | 1.65 | 0.01 | 1 | 0.56 | 1359.93 | | 1.49 | 0.01 | 1 | 0.69 | 1360.04 |
| | Proverb | 1.12 | 0.16 | 1 | 0.44 | 1459.05 | | 4.67 | 0.17 | 1 | 0.23 | 1454.21 |
| | Syn_comp | 4.27 | 0.01 | 1 | 0.52 | 1439.58 | | 1 | 0 | 0.9 | 0.62 | 1436.36 |
| | Sem_comp | 1 | 0 | 0.69 | 0.61 | 1435.99 | | 1.57 | 0 | 1 | 0.49 | 1435.75 |
| | RT_Syn_comp | 1 | 0.02 | 1 | 0.47 | 1109.11 | | 1 | 0.02 | 1 | 0.14 | 1106.8 |
| | RT_Sem_comp | 1.74 | 0.01 | 1 | 0.22 | 1127.11 | | 1 | 0 | 1 | 0.35 | 1125.52 |
| | ToT | 2.42 | 0.15 | 1 | 0.09 | 1442.43 | | 1 | 0.15 | 1 | 0.34 | 1438.46 |
| **LParsO_RSTS** | VF | 1 | 0.23 | 1 | 0.1 | 1456.72 | **LParsO_RMTG** | 1 | 0.23 | 0.16 | 0.86 | 1461.16 |
| | Vocab | 6.18 | 0.23 | 1 | 0.25 | 1472.34 | | 1.26 | 0.21 | 0.56 | 0.81 | 1478.03 |
| | High_phon | 4.34 | 0.04 | 1 | 0.01 | 1506.53 | | 1 | 0.02 | 1 | 0.28 | 1515.29 |
| | High_sem | 3.63 | 0.06 | 1 | 0.01 | 1500.39 | | 1.11 | 0.04 | 1 | 0.18 | 1510.52 |
| | Low_phon | 5.72 | 0.06 | 1 | 0.11 | 1505.18 | | 5.74 | 0.05 | 1 | 0.21 | 1508.73 |
| | Low_sem | 1.71 | 0.01 | 0.52 | 0.92 | 1525.15 | | 1 | 0.01 | 1 | 0.32 | 1524.62 |
| | RT_high_phon | 1 | 0.01 | 1 | 0.75 | 1360.37 | | 1 | 0.01 | 1 | 0.54 | 1360.43 |
| | RT_high_sem | 2.83 | 0.02 | 1 | 0.73 | 1363.07 | | 1 | 0.01 | 0.9 | 0.82 | 1362.63 |
| | RT_low_phon | 1.78 | 0.04 | 1 | 0.68 | 1352.77 | | 2.16 | 0.04 | 1 | 0.47 | 1353.94 |
| | RT_low_sem | 1 | 0.02 | 1 | 0.11 | 1354.08 | | 1.92 | 0.02 | 1 | 0.47 | 1358.67 |
| | Proverb | 3.11 | 0.17 | 1 | 0.24 | 1454.17 | | 1.83 | 0.16 | 1 | 0.4 | 1456.82 |
| | Syn_comp | 2.61 | 0.02 | 1 | 0.08 | 1430.08 | | 1.02 | 0 | 1 | 0.46 | 1438.45 |
| | Sem_comp | 1 | 0 | 0.34 | 0.76 | 1436.32 | | 1 | 0 | 1 | 0.06 | 1432.74 |
| | RT_Syn_comp | 3.06 | 0.02 | 1 | 0.72 | 1110.61 | | 2.44 | 0.03 | 1 | 0.28 | 1103.08 |
| | RT_Sem_comp | 1 | -0.01 | 0.22 | 0.89 | 1129.81 | | 1 | -0.01 | 0.38 | 0.82 | 1129.31 |
| | ToT | 1 | 0.14 | 1 | 0.62 | 1445.43 | | 1 | 0.14 | 0.15 | 0.91 | 1443.49 |
| **LParsO_RParsO** | VF | 1.91 | 0.25 | 1 | 0 | 1447.52 | **LParsO_RParsT** | 1 | 0.23 | 0.53 | 0.66 | 1464.68 |
| | Vocab | 1 | 0.21 | 1 | 0.21 | 1470.88 | | 2 | 0.22 | 1 | 0.27 | 1471.44 |
| | High_phon | 1.65 | 0.02 | 1 | 0.48 | 1516.7 | | 1.3 | 0.02 | 1 | 0.54 | 1516.17 |
| | High_sem | 5.08 | 0.05 | 1 | 0.34 | 1508.38 | | 1.41 | 0.04 | 1 | 0.57 | 1511.41 |
| | Low_phon | 1 | 0.04 | 1 | 0.56 | 1512.33 | | 2.07 | 0.05 | 1 | 0.35 | 1511.54 |
| | Low_sem | 5.86 | 0.03 | 1 | 0.08 | 1517.39 | | 1.55 | 0.01 | 1 | 0.57 | 1523.64 |
| | RT_high_phon | 2.4 | 0.02 | 1 | 0.23 | 1360.36 | | 1.76 | 0.02 | 1 | 0.11 | 1358.78 |
| | RT_high_sem | 4.36 | 0.03 | 1 | 0.08 | 1356.9 | | 1 | 0.01 | 1 | 0.08 | 1359.26 |

| | | | | | | | | | | | | |
|---|---|---|---|---|---|---|---|---|---|---|---|---|
| | RT_low_phon | 2.25 | 0.04 | 1 | 0.05 | 1349.08 | | 1 | 0.06 | 1 | 0 | 1339.5 |
| | RT_low_sem | 4.38 | 0.06 | 1 | 0 | 1341.03 | | 1.73 | 0.02 | 1 | 0.14 | 1357.38 |
| | Proverb | 1 | 0.17 | 1 | 0.03 | 1451.36 | | 2.07 | 0.17 | 1 | 0.22 | 1456.93 |
| | Syn_comp | 2.92 | 0.01 | 1 | 0.21 | 1435.05 | | 2 | 0 | 1 | 0.44 | 1438.19 |
| | Sem_comp | 1 | 0 | 1 | 0.28 | 1434.9 | | 1 | 0 | 0.27 | 0.8 | 1436.5 |
| | RT_Syn_comp | 1 | 0.02 | 1 | 0.28 | 1107.89 | | 1 | 0.02 | 1 | 0.1 | 1106.4 |
| | RT_Sem_comp | 3.44 | 0.02 | 1 | 0.07 | 1122.84 | | 2.79 | 0.02 | 1 | 0.06 | 1124.45 |
| | ToT | 2.35 | 0.15 | 1 | 0.33 | 1445.48 | | 4.03 | 0.14 | 1 | 0.69 | 1450.07 |
| **LParsO_RSTG** | VF | 1 | 0.23 | 0.48 | 0.7 | 1458.21 | **LParsT_RSTS** | 1 | 0.24 | 1 | 0.24 | 1457.89 |
| | Vocab | 2.57 | 0.22 | 1 | 0.55 | 1473.69 | | 1 | 0.21 | 1 | 0.44 | 1473.61 |
| | High_phon | 1.22 | 0.02 | 1 | 0.62 | 1515.92 | | 1 | 0.01 | 1 | 0.54 | 1516.81 |
| | High_sem | 5.17 | 0.08 | 1 | 0 | 1498.72 | | 1.38 | 0.04 | 1 | 0.3 | 1510.09 |
| | Low_phon | 1 | 0.04 | 1 | 0.13 | 1511.57 | | 1 | 0.04 | 1 | 0.47 | 1513.32 |
| | Low_sem | 1 | 0.01 | 1 | 0.2 | 1524.32 | | 1.3 | 0 | 0.56 | 0.88 | 1526.19 |
| | RT_high_phon | 1 | 0.02 | 1 | 0.72 | 1358.76 | | 1 | 0.01 | 1 | 0.27 | 1360.13 |
| | RT_high_sem | 1 | 0.01 | 1 | 0.46 | 1359.02 | | 1 | 0.01 | 1 | 0.23 | 1359.92 |
| | RT_low_phon | 2.85 | 0.05 | 1 | 0.25 | 1347.93 | | 2.14 | 0.04 | 1 | 0.11 | 1350.7 |
| | RT_low_sem | 1.8 | 0.01 | 1 | 0.45 | 1359.88 | | 3.6 | 0.04 | 1 | 0.16 | 1355.36 |
| | Proverb | 2.95 | 0.16 | 1 | 0.67 | 1457.95 | | 2.04 | 0.16 | 1 | 0.38 | 1456.66 |
| | Syn_comp | 1 | 0 | 1 | 0.35 | 1437.38 | | 1 | 0 | 1 | 0.18 | 1435.9 |
| | Sem_comp | 3.66 | 0.12 | 1 | 0 | 1379.61 | | 1 | 0 | 1 | 0.25 | 1435.17 |
| | RT_Syn_comp | 1 | 0.01 | 1 | 0.49 | 1110.15 | | 3.05 | 0.04 | 1 | 0.01 | 1101.09 |
| | RT_Sem_comp | 3.71 | 0.02 | 1 | 0.44 | 1126.51 | | 1 | 0 | 1 | 0.34 | 1129.39 |
| | ToT | 2.42 | 0.14 | 1 | 0.74 | 1450.43 | | 1 | 0.14 | 1 | 0.3 | 1444.58 |
| **LParsT_RMTG** | VF | 1 | 0.23 | 0.16 | 0.89 | 1461.44 | **LParsT_RParsO** | 2.12 | 0.24 | 1 | 0.17 | 1459.98 |
| | Vocab | 1.96 | 0.22 | 1 | 0.22 | 1472.97 | | 2.1 | 0.22 | 1 | 0.44 | 1471.94 |
| | High_phon | 1.77 | 0.02 | 1 | 0.73 | 1516.11 | | 1.39 | 0.02 | 1 | 0.34 | 1516.4 |
| | High_sem | 4.28 | 0.05 | 1 | 0.14 | 1506.53 | | 1 | 0.03 | 1 | 0.62 | 1511.47 |
| | Low_phon | 1 | 0.04 | 1 | 0.38 | 1513.49 | | 1 | 0.04 | 1 | 0.46 | 1513.16 |
| | Low_sem | 1 | 0 | 0.98 | 0.54 | 1526.2 | | 1 | 0.01 | 1 | 0.14 | 1523.71 |
| | RT_high_phon | 1 | 0.01 | 1 | 0.43 | 1361.36 | | 1.46 | 0.02 | 1 | 0.38 | 1360.1 |
| | RT_high_sem | 1 | 0.01 | 1 | 0.2 | 1361.88 | | 1 | 0.01 | 1 | 0.5 | 1360.15 |
| | RT_low_phon | 1.87 | 0.04 | 1 | 0.72 | 1354.22 | | 1 | 0.03 | 0.99 | 0.76 | 1354.99 |
| | RT_low_sem | 1 | 0.02 | 1 | 0.03 | 1355.05 | | 1 | 0.01 | 0.75 | 0.7 | 1360.04 |
| | Proverb | 2.3 | 0.17 | 1 | 0.08 | 1452.48 | | 1 | 0.16 | 1 | 0.63 | 1459.05 |
| | Syn_comp | 1.94 | 0 | 1 | 0.8 | 1439.9 | | 1 | 0 | 1 | 0.04 | 1434.11 |
| | Sem_comp | 1 | 0.01 | 1 | 0.02 | 1429.86 | | 2.58 | 0.04 | 1 | 0.01 | 1419.12 |
| | RT_Syn_comp | 1 | 0.01 | 1 | 0.39 | 1109.89 | | 1 | 0.01 | 0.34 | 0.87 | 1110.15 |
| | RT_Sem_comp | 4.39 | 0.01 | 1 | 0.38 | 1127.76 | | 1.06 | - 0.01 | 0.22 | 0.97 | 1129.52 |
| | ToT | 2.14 | 0.14 | 1 | 0.42 | 1446.29 | | 1 | 0.15 | 1 | 0.11 | 1446.05 |
| **LParsT_RParsT** | VF | 1.77 | 0.23 | 1 | 0.51 | 1462.4 | **LParsT_RSTG** | 1 | 0.24 | 0.85 | 0.57 | 1458.92 |
| | Vocab | 2.42 | 0.23 | 1 | 0.12 | 1469.89 | | 1.36 | 0.21 | 0.56 | 0.94 | 1478.25 |
| | High_phon | 1.68 | 0.02 | 1 | 0.37 | 1517.32 | | 1.32 | 0.03 | 1 | 0.83 | 1513.25 |
| | High_sem | 2.04 | 0.04 | 1 | 0.27 | 1510.1 | | 1 | 0.04 | 1 | 0.01 | 1505.17 |
| | Low_phon | 1.55 | 0.04 | 1 | 0.27 | 1512.42 | | 2.16 | 0.05 | 1 | 0.29 | 1510.25 |
| | Low_sem | 2.94 | 0.02 | 1 | 0.42 | 1524.02 | | 1 | 0.02 | 0.56 | 0.75 | 1518.57 |
| | RT_high_phon | 1.68 | 0.01 | 1 | 0.74 | 1361.76 | | 2.48 | 0.02 | 1 | 0.13 | 1358.25 |

| | | | | | | | | | | | | | |
|---|---|---|---|---|---|---|---|---|---|---|---|---|---|
| | RT_high_sem | 1 | 0.01 | 1 | 0.16 | 1358.96 | | | 4.88 | 0.04 | 1 | 0.02 | 1353.32 |
| | RT_low_phon | 1 | 0.03 | 1 | 0.13 | 1352.86 | | | 3.1 | 0.05 | 1 | 0.04 | 1346.58 |
| | RT_low_sem | 1 | 0.02 | 0.75 | 0.72 | 1358.16 | | | 2.85 | 0.03 | 1 | 0.07 | 1352.37 |
| | Proverb | 2.35 | 0.17 | 1 | 0.18 | 1453.94 | | | 1 | 0.16 | 1 | 0.51 | 1454.5 |
| | Syn_comp | 2.97 | 0.01 | 1 | 0.03 | 1431.75 | | | 1 | 0 | 1 | 0.5 | 1437.02 |
| | Sem_comp | 1 | 0 | 0.95 | 0.46 | 1435.31 | | | 1 | 0.01 | 1 | 0.12 | 1430.36 |
| | RT_Syn_comp | 1 | 0.01 | 1 | 0.64 | 1110.37 | | | 1 | 0.01 | 1 | 0.54 | 1108.96 |
| | RT_Sem_comp | 1 | -0.01 | 0.73 | 0.68 | 1130.18 | | | 1 | -0.01 | 0.73 | 0.65 | 1129.75 |
| | ToT | 1.25 | 0.15 | 1 | 0.77 | 1441.46 | | | 1 | 0.14 | 1 | 0.67 | 1444.43 |
| **LSTG_RSTS** | VF | 2.24 | 0.24 | 1 | 0.12 | 1460.02 | | **LSTG_RMTG** | 1 | 0.24 | 1 | 0.12 | 1456.49 |
| | Vocab | 1 | 0.21 | 0.56 | 0.79 | 1473.82 | | | 1.46 | 0.22 | 1 | 0.63 | 1478.86 |
| | High_phon | 1 | 0.01 | 1 | 0.35 | 1516.37 | | | 1 | 0.02 | 1 | 0.63 | 1512.05 |
| | High_sem | 1 | 0.03 | 0 | 0.99 | 1511.42 | | | 1 | 0.04 | 1 | 0.24 | 1508.88 |
| | Low_phon | 1 | 0.04 | 1 | 0.05 | 1509.61 | | | 2.63 | 0.04 | 1 | 0.48 | 1513.66 |
| | Low_sem | 1 | 0.01 | 1 | 0.28 | 1524.25 | | | 3.52 | 0.01 | 1 | 0.47 | 1524.26 |
| | RT_high_phon | 1 | 0.02 | 0 | 0.95 | 1362.2 | | | 2.82 | 0.03 | 1 | 0.06 | 1355.03 |
| | RT_high_sem | 1 | 0.01 | 1 | 0.46 | 1361.68 | | | 1 | 0.01 | 1 | 0.53 | 1362.45 |
| | RT_low_phon | 1.35 | 0.03 | 1 | 0.67 | 1354.91 | | | 4.68 | 0.04 | 1 | 0.32 | 1353.81 |
| | RT_low_sem | 1 | 0.01 | 0.3 | 0.9 | 1360.31 | | | 1.54 | 0.02 | 1 | 0.36 | 1359.19 |
| | Proverb | 1 | 0.16 | 1 | 0.74 | 1453.44 | | | 3.07 | 0.17 | 1 | 0.28 | 1454.26 |
| | Syn_comp | 1 | 0 | 0.32 | 0.82 | 1438.66 | | | 1.92 | 0.05 | 1 | 0.06 | 1418.55 |
| | Sem_comp | 1 | 0 | 1 | 0.37 | 1435.62 | | | 1 | -0.01 | 0 | 0.99 | 1436.66 |
| | RT_Syn_comp | 1 | 0.01 | 0 | 0.98 | 1110.86 | | | 1 | 0.01 | 0 | 1 | 1110.27 |
| | RT_Sem_comp | 1 | 0 | 1 | 0.37 | 1129.26 | | | 1 | -0.01 | 0.36 | 0.84 | 1129.31 |
| | ToT | 1 | 0.14 | 1 | 0.4 | 1444.84 | | | 1 | 0.14 | 1 | 0.7 | 1445.45 |
| **LSTG_RParsO** | VF | 3.36 | 0.25 | 1 | 0.03 | 1453.62 | | **LSTG_RParsT** | 1 | 0.23 | 0.45 | 0.72 | 1464.92 |
| | Vocab | 2.06 | 0.22 | 1 | 0.49 | 1473.45 | | | 1.88 | 0.21 | 1 | 0.47 | 1475.18 |
| | High_phon | 2.32 | 0.02 | 1 | 0.79 | 1516.61 | | | 2.29 | 0.02 | 1 | 0.17 | 1514.7 |
| | High_sem | 1 | 0.04 | 1 | 0.03 | 1502.95 | | | 1 | 0.04 | 1 | 0.55 | 1511.53 |
| | Low_phon | 1.65 | 0.04 | 1 | 0.89 | 1514.89 | | | 1.91 | 0.05 | 1 | 0.18 | 1508.47 |
| | Low_sem | 1 | 0.02 | 0.98 | 0.54 | 1516.55 | | | 1.45 | 0 | 1 | 0.6 | 1525.3 |
| | RT_high_phon | 1 | 0.02 | 1 | 0.12 | 1356.55 | | | 2.2 | 0.02 | 1 | 0.16 | 1358.98 |
| | RT_high_sem | 1 | 0.02 | 1 | 0.14 | 1357.11 | | | 1.87 | 0.02 | 1 | 0.14 | 1360.36 |
| | RT_low_phon | 1.08 | 0.03 | 1 | 0.76 | 1353.1 | | | 1.37 | 0.03 | 1 | 0.78 | 1355.1 |
| | RT_low_sem | 1.49 | 0.02 | 0.75 | 0.87 | 1358.83 | | | 1 | 0.01 | 1 | 0.43 | 1359.57 |
| | Proverb | 3.96 | 0.17 | 1 | 0.57 | 1457.73 | | | 1.88 | 0.17 | 1 | 0.42 | 1451.19 |
| | Syn_comp | 1 | 0 | 1 | 0.34 | 1437.84 | | | 1 | 0 | 0.09 | 0.92 | 1438.4 |
| | Sem_comp | 1 | -0.01 | 0 | 0.96 | 1436.65 | | | 1 | 0 | 0.15 | 0.9 | 1436.96 |
| | RT_Syn_comp | 1 | 0.01 | 1 | 0.57 | 1109.21 | | | 1.01 | 0.01 | 1 | 0.54 | 1109.21 |
| | RT_Sem_comp | 2.13 | 0.01 | 1 | 0.1 | 1126.16 | | | 3.13 | 0.01 | 1 | 0.28 | 1128.65 |
| | ToT | 1.19 | 0.14 | 1 | 0.72 | 1450.12 | | | 1 | 0.14 | 1 | 0.39 | 1444.81 |
| **LSTG_RSTG** | VF | 1 | 0.23 | 0.16 | 0.87 | 1461.18 | | | | | | | |
| | Vocab | 1.15 | 0.21 | 0.87 | 0.82 | 1479.81 | | | | | | | |
| | High_phon | 1.33 | 0.02 | 1 | 0.58 | 1516.33 | | | | | | | |
| | High_sem | 1 | 0.04 | 1 | 0.1 | 1507.64 | | | | | | | |

| | | | | | | |
|---|---|---|---|---|---|---|
| | Low_phon | 2.56 | 0.05 | 1 | 0.09 | 1508.65 |
| | Low_sem | 2.09 | 0.01 | 1 | 0.09 | 1521.7 |
| | RT_high_phon | 5.37 | 0.04 | 1 | 0.07 | 1355.06 |
| | RT_high_sem | 3.88 | 0.05 | 1 | 0 | 1347.28 |
| | RT_low_phon | 2.51 | 0.04 | 1 | 0.55 | 1354.72 |
| | RT_low_sem | 4.08 | 0.03 | 1 | 0.14 | 1356.08 |
| | Proverb | 1 | 0.16 | 1 | 0.32 | 1455.69 |
| | Syn_comp | 6.2 | 0.05 | 1 | 0.16 | 1428.12 |
| | Sem_comp | 1 | 0 | 0.11 | 0.92 | 1432.35 |
| | RT_Syn_comp | 1.55 | 0.02 | 1 | 0.25 | 1106.66 |
| | RT_Sem_comp | 2.76 | 0.01 | 1 | 0.41 | 1128.81 |
| | ToT | 1 | 0.14 | 1 | 0.4 | 1444.91 |
